# Dicer is essential for proper maturation, composition, and function in the postnatal retina

**DOI:** 10.1101/2025.07.30.635135

**Authors:** Seoyoung Kang, Daniel Larbi, Eik Bruns, Konstantin Hahne, Alireza Khodadadi-Jamayran, Chaitra Sreenivasaiah, Mariana Lima Carneiro, Monica Andrade, Khulan Batsuuri, Shaoheng Chen, Julia Jager, Suresh Viswanathan, Brian S. Clark, Stefanie G. Wohl

**Affiliations:** Department of Biological and Vision Sciences, The State University of New York, College of Optometry, New York, New York, USA; Applied Bioinformatics Laboratories, Office of Science and Research, New York University School of Medicine, New York, NY, USA; Indiana University School of Optometry, Bloomington, Indiana, USA; John F Hardesty, MD Department of Ophthalmology and Visual Sciences and Department of Developmental Biology, Washington University School of Medicine, St. Louis, MO, USA

**Keywords:** Müller glia, rod photoreceptors, bipolar cells, amacrine cells, miR-25, miR-20, Elavl3, EdU, HuC/D, ERG, OCT, Luciferase, gene ontology, target prediction

## Abstract

microRNAs (miRNAs) play a pivotal role during the early phases of retinal development, but their impact on late-phase retinogenesis is unknown. We depleted miRNAs in late retinal progenitor/precursor cells (RPCs/PCs) via a conditional Dicer knock-out. Optical coherence tomography (OCT), electroretinography (ERG), histological, and transcriptional analyses were conducted in young and adult mice. Alterations in gene expression of late-born cells were observed as early as postnatal day 7 (P7), resulting in impaired rod function, a significantly reduced number of rod bipolar cells and their associated function, and a decreased Müller glia population at adult age. These defects appear to be caused by a delay in differentiation/ incomplete maturation, as indicated by an enlarged progenitor/precursor population at young ages that persists into adulthood. Notably, an increased population of HuC/D+ amacrine cells was found. Luciferase assays led us to speculate that this increase may be due to the absence of *Elavl3* suppression via RPC-miRNAs. This suggests that Dicer/miRNAs in late RPC/PCs are essential for the proper formation and maturation of late RPC progenies and may also play a role in regulating cell state.

**Graphical abstract:** 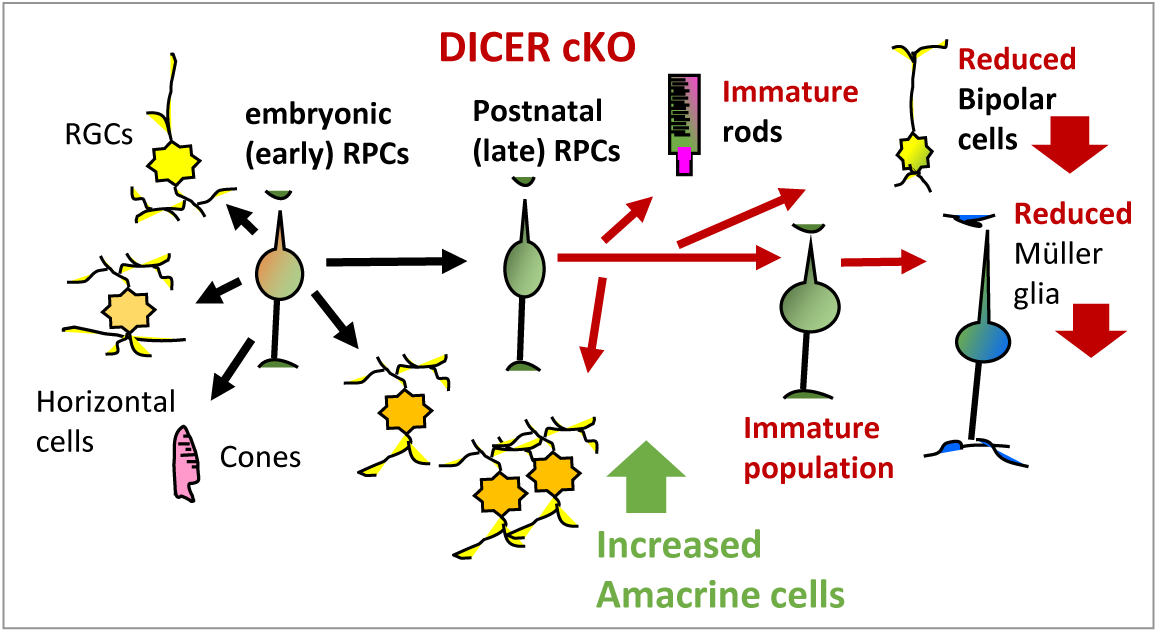

**Summary statement:** Late-retinal progenitor microRNAs are essential for proper postnatal retinogenesis and retinal function.

## Introduction

Retinogenesis is a complex and highly regulated process across all vertebrates. There are two principal phases in which all six retinal neuron types are generated: First, retinal ganglion cells (RGCs), cone photoreceptors, horizontal cells (HCs), and early-born amacrine cell types (ACs) arise from early (embryonic) retinal progenitor cells (eRPCs); second: rod photoreceptors, bipolar cells (BCs), late-born ACs, and Müller glia (MG), the only non-neuronal cell type, are generated from late, postmitotic (p) RPCs (Cook, 2003, Jin and Xiang, 2017, Marquardt and Gruss, 2002, Rapaport et al., 2004, Xiang, 2013, Andreazzoli, 2009, Hoang et al., 2020, Hahn et al., 2023, Young, 1985a, Prada et al., 1991, LaVail et al., 1992, Cepko et al., 1996, Brzezinski et al., 2011). All retinal cell types are found in designated retinal layers, forming distinct connections to neighboring cells, and express their cell-type-specific markers by postnatal day (P) 12. Full tissue maturation is, however, not completed until P28 (full visual acuity) (Prusky et al., 2004).

Various molecules, primarily specific combinations of transcription factors, have been identified in the past decades that play a role in neuro- and gliogenesis in the nervous system, including the neural retina. Dicer/microRNAs (miRNAs) are known to regulate cell proliferation, cell fate decisions, cell differentiation, and maturation in various tissues, including the brain and the neural retina (Bian et al., 2013, Foshay and Gallicano, 2009, Lee et al., 2018, Lu et al., 2012, He and Hannon, 2004, Fairchild et al., 2019, La Torre et al., 2013, Akerblom and Jakobsson, 2013, Ambros, 2011, Amini-Farsani and Asgharzade, 2019, Bueno and Malumbres, 2011, Coolen et al., 2013, Corbin et al., 2009, Sun et al., 2013, Vo et al., 2010). These short (18-22 nucleotides long), non-coding RNAs act as translational repressors (Lee et al., 1993, Gurtan and Sharp, 2013, Sundermeier and Palczewski, 2012, Zuzic et al., 2019). They are encoded in genes of exons or introns and processed by the ribonuclease III (RNAse III) enzyme complexes called Drosha- and Dicer-complex. The Drosha-complex generates a precursor molecule from the primary transcript, while the Dicer-complex processes the precursor molecule into the mature functional miRNA (Gurtan and Sharp, 2013, Sundermeier and Palczewski, 2012, Zuzic et al., 2019). A very efficient method to study the role of tissue- or cell-type-specific miRNAs is therefore via loss-of-function experiments by conditionally deleting the enzyme (or its function) that generates mature miRNAs, Dicer. To our knowledge, Dicer-conditional knock-out (cKO) studies have only been carried out exclusively during embryonic retinal development, resulting in mostly severe outcomes including microphthalmia, rosette formation, and massive cell loss, leading to severe functional defects (Damiani et al., 2008, Decembrini et al., 2008, Iida et al., 2011, Pinter and Hindges, 2010). Intriguingly, the loss of Dicer/ mature miRNAs in eRPCs (E12/14) led to an overproduction of early-born cells (RGCs, HCs, ACs) and a failure to produce late-born cells (rods, BCs, MG), as shown by two independent laboratories (Georgi and Reh, 2010, Davis et al., 2011). However, most of these studies were performed over a decade ago. Many techniques have advanced over the years, now allowing for defined manipulation and evaluation at the single-cell level. Furthermore, to date, no data are available regarding the role of Dicer and miRNAs in postnatal RPCs/PCs in terms of the generation, maturation, and functionality of their progeny cells, specifically rods, BCs, late-born ACs, and MG.

This study aimed to analyze the impact of miRNA loss in late (postnatal) RPCs/PCs, with a focus on the adult retina. Retinal composition, function, and overall health were assessed using spectral-domain optical coherence tomography (SD-OCT), electroretinograms (ERGs), histological approaches, and single-cell as well as bulk RNA sequencing (RNA-seq). We utilized the late RPC-specific *Ascl1CreERT:tdTomato* reporter mouse (Brzezinski et al., 2011, Kim et al., 2011, Kim et al., 2007). Ascl1 (<u>A</u>chaete-<u>Sc</u>ute homolog 1, also known as Mash1), a pro-neural basic helix loop helix [bHLH] transcription factor, is expressed in late RPCs/PCs that generate the late-born retinal cells. It is not expressed in differentiated neurons or glia (Brzezinski et al., 2011, Nelson et al., 2011). This reporter mouse was crossed with the Dicer^fl/fl^ mouse. Reporter expression/ Dicer loss was induced at postnatal days (P) 1-3, i.e., the peak of the second phase of retinogenesis (Young, 1985b, Young, 1985a). Cell and molecular analyses were performed at P7 and P14; structural/functional *in vivo* evaluations, in combination with histology, were conducted at P28, P56, 3 months, and 4 months of age.

Our data show that loss of Dicer/mature miRNAs in postnatal RPCs/PCs results in prolonged cell division events and a delay of maturation in the young postnatal retina. This delay leads to functional impairments and a reduction of the overall number of late RPC/PC progenies in the adult retina, predominantly in the nasal periphery. The most affected were central rod BCs and peripheral MG. Overall, the process of retinogenesis appeared incomplete, resulting in a residual immature population in the adult retina. The failure to mature correctly results in functional impairments. Subsequent retinal degeneration becomes evident as early as two months after Dicer deletion, but progresses very slowly. Moreover, Dicer-cKO_RPC_ retinas have an increased HuC/D+ AC population. We hypothesized that increased *Elavl3* expression, an upregulated RPC-miRNA target and the gene encoding HuC protein, might cause this. Overall, postnatal RPC-miRNAs appear to be essential for the proper maturation of late-born retinal populations.

## Results

### Dicer loss in late RPCs/precursors results in maturation delays

To label, trace, and analyze the progenies of late RPCs and investigate the impact of Dicer loss in the second phase of retinogenesis (Fig. 1A), we used the Ascl1-CreERT:tdTomato mouse (referred to as wildtype) and crossed it with the Dicer-cKO_RPC_ mouse (referred to as cKO, Fig. 1B). *Cre* induction at P1-3 resulted (Fig. 1C) in sufficient reporter expression throughout the wildtype retina. At P4, the vast majority of cells in the neuroblastic layer (NBL) were reporter+ (Fig. 1D). At P7, many Tomato+ cells were found in the inner and outer nuclear layers (INL/ONL); hence, most late-born cells, including rod photoreceptors, were labeled. At P21, very bright Tomato+ cells were found throughout the INL. Reporter signals in photoreceptors in the ONL became fainter over time (Fig. 1D), very likely due to the inactivation of Tomato expression in the *Rosa26* locus of mature neurons (Godecke et al., 2017, Long and Rossi, 2009).

**Figure 1.**
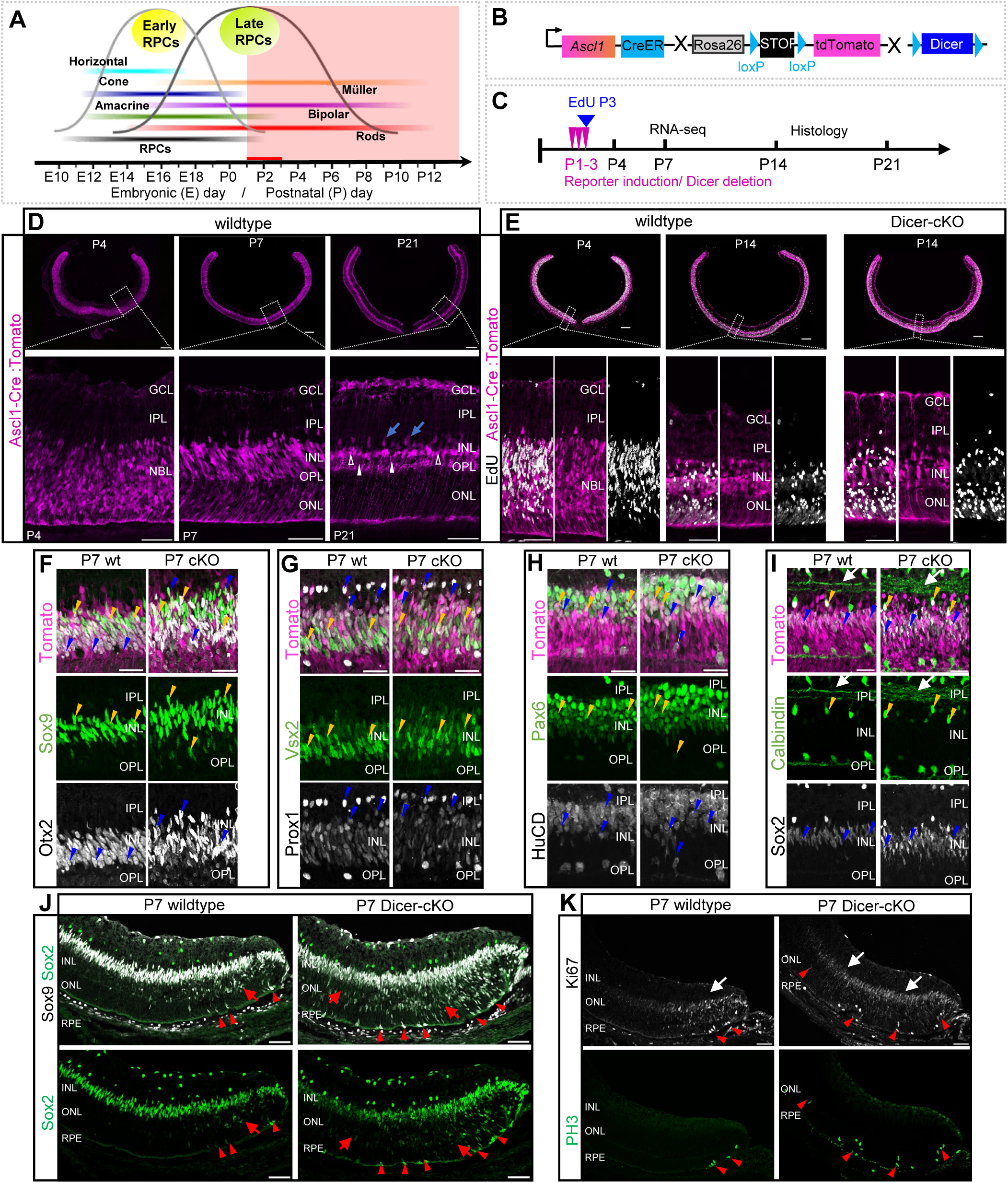
Dicer-cKO retinas have an enlarged progenitor/precursor cell population. **A:** Mouse retinogenesis schematic with time point of manipulation (red line, after *Jin et al., 2016*). **B:** Schematic of retinal progenitor cell (RPC) reporter mouse (wildtype, wt) and Dicer-cKO mouse. **C:** Experimental design with time points of *Cre* induction and EdU labeling. **D-E:** P4, P7, P14, and P21 wildtype and cKO retinal cross sections with magnified insets visualizing endogenous reporter expression of RPC progenies labeled at P1-3, that underwent cell division at P3; blue arrows in D show presumptive amacrine cells, white arrowheads presumptive bipolar cells, unfilled arrowheads presumptive Müller glia; EdU+ progenies are shown in E . **F-I:** Labeling with antibodies against Sox9, Otx2 (F), Vsx2, Prox1 (G), Pax6, HuCD (H), and Calbindin, Sox2 (I) of the INL of P7 wildtype or Dicer-cKO central retinal cross sections to characterize tdTomato+ RPC progenies (arrowheads). Arrows indicate stratification in the INL. **J-K:** Sox2 and Sox9 (J) or Phosphohistone 3 (PH3) and Ki67 labeling (K) of P7 peripheral retinal wildtype or Dicer-cKO cross sections; red arrows show migrating RPCs, white areas indicate Ki67+ zones; arrowheads indicate dividing RPCs. P4 wt n=4, P4 cKO n=2, P7 wt n=4, P7 cKO n=4, P14 wt n=4, P14 cKO n=3, P21 wt n=5. Scale bars in D, E: 200 μm, insets 50 μm; F-I: 25 μm, J-K: 50 μm. GCL: ganglion cell layer, IPL: inner plexiform layer, INL: inner nuclear layer, OPL: outer plexiform layer, ONL: outer nuclear layer, NBL: neuroblastic layer, RPE: retinal pigment epithelium.

*Cre* activation in the Dicer-cKO mouse at P1-3 led to the successful excision of exon 23 of the Dicer1 gene, resulting in a functionally inactive Dicer enzyme. This was evident in P4 retinal lysates as well as in all FACS-purified P7 reporter+ cKO RNA-seq samples (reporter-negative or reporter+ wildtype samples had intact *Dicer1*, Figs. S1A, B). To evaluate whether reporter+ cells were actively dividing RPCs, a single EdU pulse was administered at P3 (Fig. 1C). At P4, the vast majority of Tomato+ cells (∼75%) were EdU+ in both wildtype and cKO retinas, hence proliferating RPCs (Figs. 1E, S1C). No drastic difference was seen in the center and periphery of both conditions. However, it appeared the cKO had slightly more EdU+ cells in the lower NBL. We then traced these cells until P14. EdU+ Tomato+ progenies were predominantly found in the ONL (PRs) and the lower and center INL (BCs/ACs/MG) of the central retina of wildtypes and cKOs. Very few cells were also seen in the upper INL and GCL, and were very likely ACs. Overall, the central and peripheral P14 cKO retinas exhibited a similar pattern of EdU+ cell distribution to that of the wildtypes. Some areas displayed slightly more cells, but that was not consistent in all sections analyzed (Figs. 1E, S1C).

We next performed an initial histological characterization of Tomato+ progenies (indicated via yellow/blue arrowheads) at P7 using the markers Otx2 and Sox9, Prox1 and Vsx2, HuC/D and Pax6, as well as Sox2 and Calbindin (Figs. 1F-I, S2). At this stage, these markers label various cell populations, including RPCs/PCs, young BCs and MG, as well as early and late ACs, and HCs, which are predominantly found in the INL. Many cells were Sox9+ and Otx2+, indicating that at this stage, RPCs, BC precursors/young BCs, and/or young MGs were present. In the cKO, some Tomato+Sox9+Otx2 cells were also found in the ONL, likely representing migrating RPCs/BC precursors or PR precursors/young PRs. Most Tomato+ ONL cells, however, exhibited a very faint, doughnut-like Otx2 expression pattern, characteristic of the typical photoreceptor expression pattern (and were Sox9-, Figs. 1F, S2A).

Vsx2 and Prox1, both RPC/BC markers, also labeled the majority of the central INL in both conditions. The cKO, however, appeared to have more cell rows than the wildtype (Figs. 1G, S2B). Prox1 also labels differentiated ACs found as a single layer in the upper INL, adjacent to the IPL, and HCs in the lower INL, adjacent to the OPL. These populations appeared to be minimally affected in the cKO. However, slightly more, yet fainter, small, round Prox1+ cells were observed in the upper cKO INL, possibly representing young ACs. Additional AC markers, such as Pax6 and HuC/D, were also co-expressed by Tomato+ progenies, in the INL, with more cell rows in the cKO (Figs. 1H arrowheads, S2C). It also appeared that some HuC/D cells in the cKO were still migrating up to their location in the INL.

The Calbindin/Sox2 combination revealed that Tomato+ Calbindin+ ACs were found in the upper INL in both conditions. Some co-expressed Sox2, hence being cholinergic ACs (Whitney et al., 2014) (Figs. 1I, S2D). However, the cKO seemed to have some additional small Calbindin+ Sox2-cells in the upper INL. Tomato+ Calbindin+ HCs or RGCs were, as expected, not found in the wildtype nor cKO. Nevertheless, the stratification in the IPL, which in the wildtype showed two distinct synapse lines, was different in the cKO. Three less-defined lines were present, suggesting synaptic alterations early on (Figs. 1I, white arrows). Moreover, most Sox2+ progenies were found in the INL, exhibiting a similar pattern to Sox9, and are probably RPCs/young MG. We therefore labeled young MG using Sox9 in combination with glutamine synthetase (GS, Fig. S2E). GS was only found in retinal astrocytes in the GCL, which also expressed Sox2 and Sox9. GS was not found in MG at this age, in agreement with previous reports (Clark et al., 2019), and also not ectopically expressed in the cKO.

Since the retinal center is more differentiated, we next evaluated Sox9/Sox2 expression in the periphery. In the wildtype, many still migrating Sox9+Sox2+ RPCs/PCs with typical elongated nuclei were present in the outermost periphery adjacent to the ciliary body (Fig. 1J, arrows). A few of them were still found in the vicinity of the RPE, where RPCs divide (arrowheads). In the Dicer-cKO_RPC_, however, the peripheral Sox9+ Sox2+ population was larger, with more migrating cells (Fig. 1J, arrows). Co-labeling with the proliferation markers Ki67 and phosphohistone 3 (PH3, Fig. 1K) showed a greater number of Ki67+ cells (arrows) and approximately 2-3 times more PH3+ cells in the periphery of cKO retinas compared to wildtypes (arrowheads). Similar results were found by (Davis et al., 2011) in *αCre* mice at embryonic stages, suggesting extended cell division events after miRNA loss.

Since the different markers used are not cell-type specific at this age and histological characterizations are limited, we performed single-cell and bulk RNA-seq to decipher better the P7 cell populations/subpopulations and gene expression patterns (Figs. 2, S3). We FACS-purified P7 Tomato+ wild-type and cKO progenies, as well as collected the corresponding reporter-negative fractions of both conditions for scRNA-seq. Cell types/populations were identified based on known marker expression for retinal cell types (Clark et al., 2019, Blackshaw et al., 2004, Macosko et al., 2015) (Fig. 2A). The predominant cell populations found in the FACS-purified reporter+ wildtype retinas included the expected late-born RPC progenies, i.e., rods, BCs, and MG. We also found BC/photoreceptor precursors, neurogenic RPCs, and RPCs (Fig. 2B). Another separate, quite large population consisted of ACs. HCs, RGCs, and Starburst ACs, i.e., early-born cells, were not present or only as a tiny population within the tdTomato-fraction (Fig S3A).

**Figure 2.**
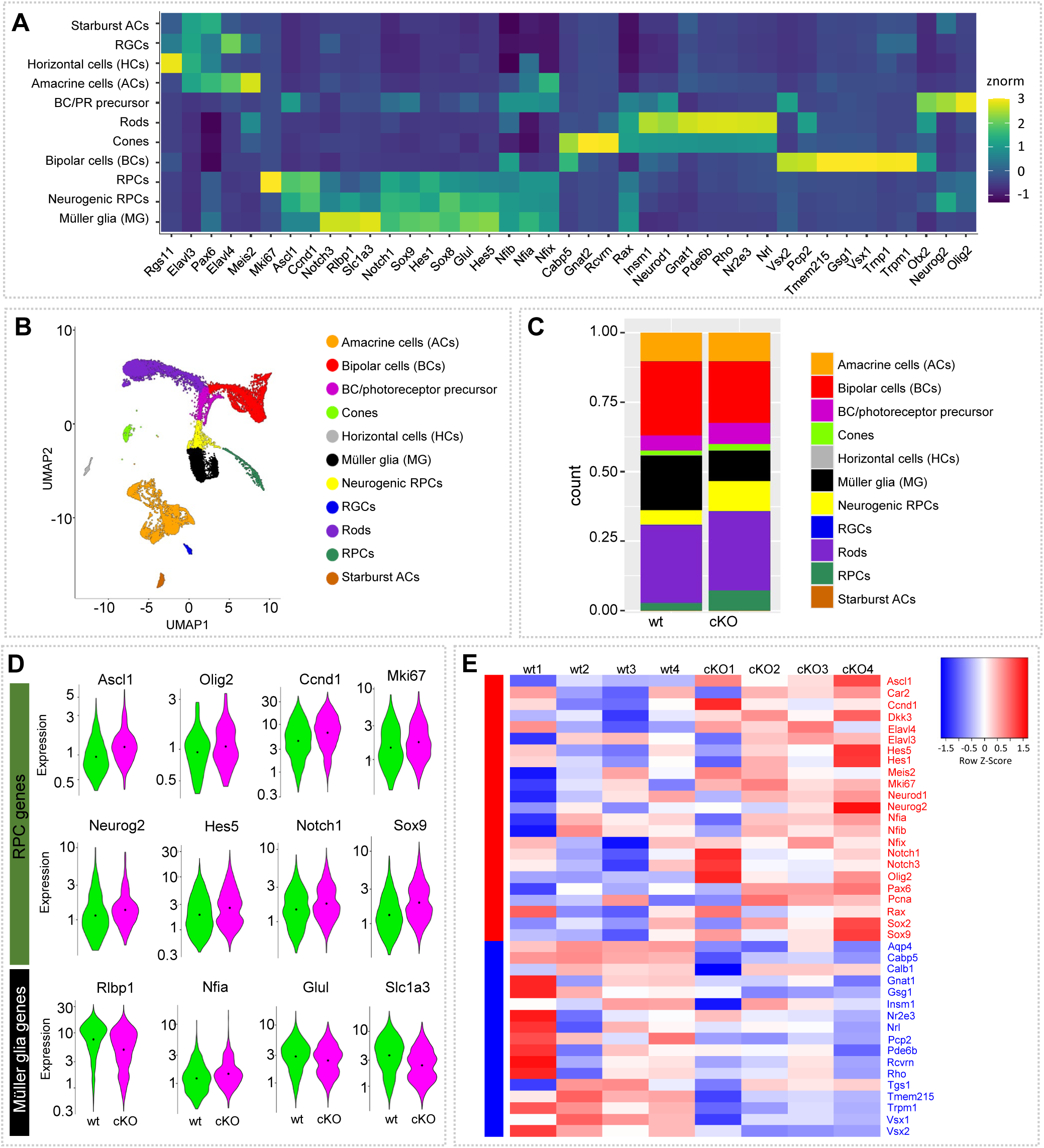
Dicer loss in late RPCs results in less mature progenies. **A-D:** scRNA-seq of FACS-purified P7 tdTomato+ wildtype (wt) and Dicer-cKO (cKO) progenies showing (A) a heatmap of expressed genes in annotated cell types, (B) UMAP-dimension reduction, and (C) cell proportions, colored by annotated cell type as determined by marker gene expression; (D) Violin plots of the cellular expression of marker genes of RPCs and Müller glia (MG), facetted by genotype; WT: n=3 mice, pooled, cKO: n=3 mice, pooled, one technical replicate. **E:** Heatmap of selected 40 up- and downregulated genes of bulk RNA-seq FACS-purified P7 Tomato+ wildtype and cKO progenies; wt: n=4 biological replicates, cKO: n=4 biological replicates, 4 independent experiments, normalized counts. RPCs: retinal progenitor cells, RGCs: retinal ganglion cells, PRs: photoreceptors.

We next compared the cell proportions found in P7 wildtype progenies with those of the cKO progenies. Overall, scRNA-seq confirmed our histological findings. The cKO had larger proportions of immature cells, i.e., RPCs, neurogenic RPCs, and BC/rod precursors (Figs. 2C, S3A). Fractions for mature BCs and MG were smaller than those of the wildtype, suggesting maturation deficits of these two populations. The fractions of rod and AC populations were, however, similar in both conditions. Wildtype non-progenies (tomato-negative cells) contained a large AC fraction, Starburst ACs, HCs, and RGCs. cKO non-progenies (tomato-negative cells) contained a smaller number of ACs, only a few HCs, RGCs, and Starburst ACs (Fig. S3A). This suggests that these cell populations may be indirectly affected by the loss of Dicer in late RPCs (e.g., missing propped connection), potentially impacting their health as well. We also found a small fraction of cones among our wildtype and cKO progenies. Since cones are only born during embryonic stages, we believe that this result is due to cell carryover. MGs are very tightly associated with/wrap both PR types, resulting often in some PR contamination of FACS-purified cells with cytoplasmic reporter proteins (Wohl and Reh, 2016).

We next analyzed and compared the expression patterns in the specific populations of wildtype and cKO progenies, starting with RPC/precursor genes and MG genes. This differentiation is not always easy since MG express many RPC genes (Jadhav et al., 2009, Roesch et al., 2008). Interestingly, *Ascl1, Olig2, Neurog2,* and *Sox9*, all RPC/precursor genes as well as cell cycle genes, such as *Mki67* and *Ccnd1,* were upregulated in the cKO, (Fig. 2D, Fig. S3B). By contrast, known MG markers, including *Rlbp1*, *Nfia*, *Glul*, and *Slc1a3* (Clark et al., 2019, Macosko et al., 2015) showed reduced expression levels.

In addition to scRNA-seq, we also conducted bulk RNA-seq of FACS-purified P7 progenies. Although this dataset represents a heterogeneous population, bulk RNA-seq allows a better sequencing depth and validation. The heatmap in Fig. 2A shows 30 selected genes known to be specific to RPCs, BCs, PRs, MG, and ACs (Clark et al., 2019, Akimoto et al., 2006, Macosko et al., 2015, Shekhar et al., 2016, Kim et al., 2016a). This selection also includes genes whose protein expression we have evaluated over time via histological analysis. We identified RPC/precursor genes (e.g., *Ascl1, Hes1, Rax, Dkk3*), including cell cycle genes (e.g., *Ccnd1, Mki67, Pcna*), which were upregulated (Fig. 2E, Table S1). Also upregulated were *Meis2*, *Pax6, Cdkn1b (encodes for* p57kip2) genes known to play a role in RPC state as well as AC fates (Bumsted-O’Brien et al., 2007, Dupacova et al., 2021, Yan et al., 2020, Dyer and Cepko, 2001a, Dyer and Cepko, 2001b). Furthermore, we found *Elavl3* and *Elavl4* upregulated, both genes with essential functions during development and AC formation (Lu et al., 2021, Wu et al., 2021, Bronicki and Jasmin, 2013, Wutikeli et al., 2025, Yokoi et al., 2017). In contrast, genes of the late-born cells, including MG (*Aqp4*), BCs (*Cabp5, Insm, Vsx1, Tmem215, Gsg1*), and rods (*Gnat2, Rcvrn, Nrl, Nr3e3, Rho*), were downregulated in the cKO compared to the wildtype (Figs. 2E, Tables S1). Apoptosis makers, including Caspases or Bax, were not found to be upregulated in P7 cKOs, suggesting no increased apoptosis at least at this stage (Tables S1). This is different from observations made in embryonic Dicer loss studies (Iida et al., 2011, Georgi and Reh, 2010).

### P56 cKO mice have fewer Müller glia and an unknown putative immature population

Since MG seemed to be affected and play a crucial role in overall retinal health, we next examined adult cKO retinas concerning their cellular composition of the glia. To evaluate the total number of MG and the fractions of RPCs that differentiated into MG, we labeled P56 tissue with antibodies against glutamine synthetase (GS) and Sox9 (Figs. 3A-D). Wildtype MG somata were almost exclusively located in the center of the INL (arrowheads), in central and peripheral areas. The MG processes spanned the entire retina to form the inner and outer limiting membranes (arrows), confirming that the vast majority of MG was labeled using this reporter mouse (Figs. 3A-B).

**Figure 3.**
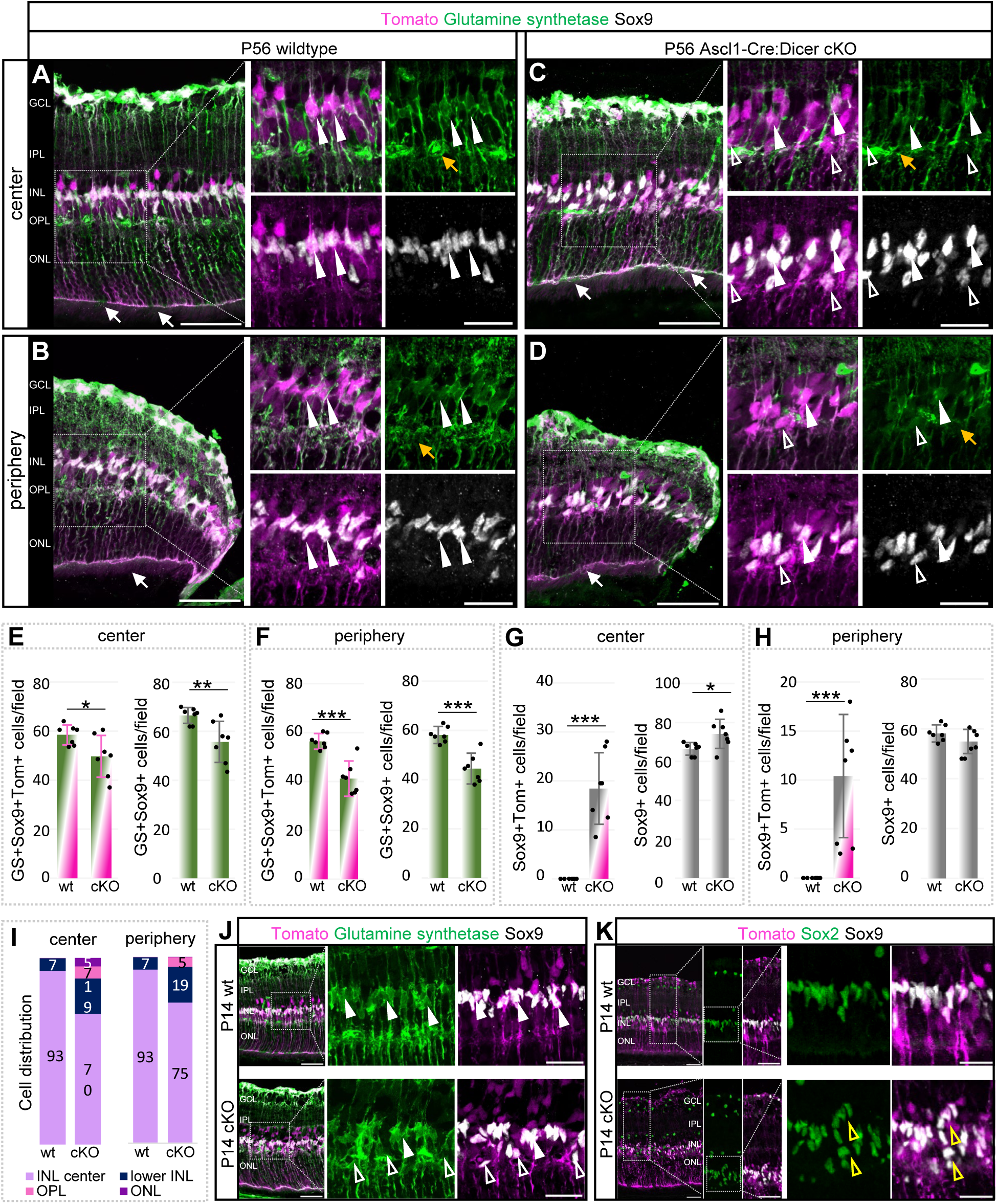
Dicer loss in late RPCs results in reduced Müller glia population and a new, unknown population. **A-D:** Müller glia (MG) labeling with antibodies against glutamine synthetase (GS) and Sox9 of central or peripheral P56 wildtype (wt) or cKO retinas, with insets shown in high magnification; white arrowheads indicate GS+ Sox9+ MG, unfilled arrowheads GS- Sox9+ cells, white arrows the external limiting membrane (ELM), yellow arrows alterations in MG processes. **E-F:** Absolute number of Tomato+ GS+ Sox9+ MG and total GS+ Sox9+ MG per field in the central or peripheral retina. **G-H:** Absolute overall number of Tomato+ Sox9+ cells and total number of Sox9+ (GS-) per field of the central or peripheral retina. **I:** Tomato+ Sox9+ cell distribution (percentage) in the center INL, lower INL, OPL, and ONL of the central and peripheral wildtype and cKO retina. **J-K:** GS and Sox9 (J) and Sox2 and Sox9 (K) labeling of P14 wildtype and cKO central retinas, with higher magnification insets; white arrowheads show Sox9+ GS+ MG, unfilled arrowheads Sox9+ GS- cells, yellow arrowheads stacked cells; wt: n=7, cKO: n=7, mean ± S.D., Mann-Whitney-U-test: *: p ≤ 0.05; **: p ≤ 0.01; ***: p ≤ 0.005. Scale bars in A-D, O-R: 50 µm, insets: 25 µm. The layer description is given in Figure 1.

In the Dicer-cKO_RPC_, central and peripheral retinal areas had many displaced and scattered Sox9+ cells throughout the INL, OPL, and ONL (Figs. 3C-D). Most cells were GS+ MG (arrowheads). These glia, however, did not have fine, branched processes in the OPL (particularly in peripheral areas, indicated by yellow arrows), suggesting possible maturation deficits. Nevertheless, the OLM was formed and seemed intact (white arrows). Cell counts of the total number of Tomato+ MG as well as overall MG numbers in the central and peripheral cKO retina showed a ∼15% and ∼25% decline of both populations, respectively, compared to the wildtype (Figs. 3E-F). 90% of all MG found in the wildtype or cKO were reporter+, and the fraction of Tomato-labeled MG was similar in wildtype and cKO, with approximately 90-97%.

Some of the displaced Sox9+ cells (ONL) in the cKO had elongated nuclei and were GS-negative (Figs. 3C-D, unfilled arrowheads), suggesting a putative immature cell population that did not migrate to its intended destination (residual RPCs). This population comprised approximately one-fourth of all Sox9+ cells in the center and about one-sixth in the periphery. The overall number of Sox9+ cells in the cKO was, however, not much different from that in wildtypes, suggesting that these displaced cells are very likely immature cells that failed to differentiate into glia, resulting in their ectopic location in the outer retina (18-23% displaced cells, Figs. 3G-H, the layer distribution (%) is shown I). Interestingly, a similar phenotype, i.e., dislocated Sox9+ cells, mostly GS-MG, has been reported previously in an MG-specific Dicer-cKO (Wohl et al., 2017). Although in the MG study, young differentiated MG were targeted and manipulated, which differs from the present study in which RPC/PCs, including MG progenitors/precursors, were targeted and manipulated, similar dedifferentiation events could cause this outcome, at least in the central (more mature) retina of this RPC-Dicer-cKO study.

Since we could not detect any GS expression in P7 MG (Fig. S2E), we next analyzed P14 tissue to ascertain whether these Sox9+ GS-negative cells were present early on (Figs. 3J, unfilled arrowheads). We found displaced Tomato+Sox9+ cells (GS-negative) in the P14 cKO, but not in the wildtype. These displaced cells were also Sox2+, and were partly arranged in stacks (Figs. 3K, unfilled arrowheads), presumably representing remaining resident RPCs. Interestingly, similar progenitor stacks/columns were also seen in embryonic Dicer-cKO studies (Davis et al., 2011).

### miRNA loss in late retinal progenitor cells results in impaired rod photoreceptor function but does not affect cones

To investigate the observed impact of this maturation delay on structural and functional outcomes in the adult cKO mouse, we performed SD-OCT and ERGs, paired with histological evaluations at P28, P56, as well as 3 and 4 months of age (Fig. 4A). SD-OCT was conducted at the central retinas of approximately 650 μm radius from the optic nerve (ON). Measurements were performed across the nasal-temporal and superior-inferior axes (Figs. 4B-C, S4-S5). The OCT scans of wildtypes across the different ages showed intact retinas with a total thickness of about 220 μm, in accordance with previous reports (Ferreira et al., 2021, Ferguson et al., 2014, Larbi et al., 2025). Somewhat surprisingly, the OCT of P28 Dicer-cKO_RPC_ mice appeared normal, and the overall retinal thickness was also not much altered later on. However, initial alterations were found at the layer level, starting at P28 and affecting the OLM-RPE area (i.e., outer/external limiting membrane (OLM/ELM), inner and outer segments (IS/OS) of photoreceptors, and the RPE). There was a ∼10% reduction in thickness compared to wildtypes, further decreasing with age, suggesting impairment of photoreceptor segments (Fig. 4C). In P56 Dicer-cKO_RPC_ mice, an initial thinning of the ONL was found (∼14% decrease), suggesting photoreceptor loss, which, however, did not further progress over time. No alterations were found in the inner retinal layers.

**Figure 4.**
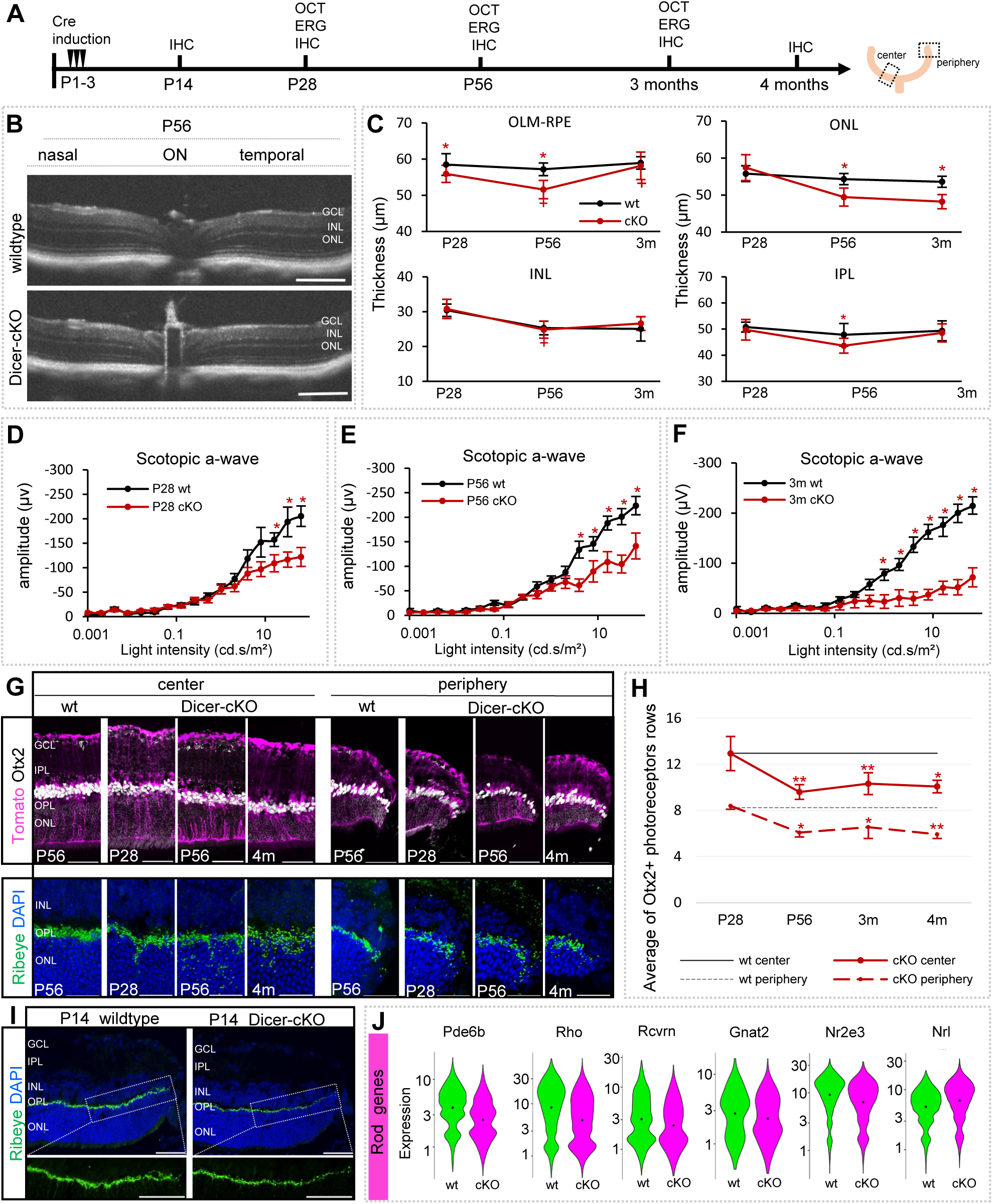
Dicer loss in late RPCs leads to reduced rod function. **A:** Experimental design and timeline. **B:** Optical coherence tomography (OCT) images of wildtype (wt) and cKO center retinas at the nasal-temporal axis. **C:** Timeline plots showing specific layer thickness changes in OCT for P28 wt (n=13) and cKO (n=10), P56 wt (n=7) and cKO (n=9), and 3-month wt (n=5) and cKO (n=4). **D-F:** Full-field scotopic electroretinogram recordings showing a-wave amplitudes of wt and cKO mice at P28 (wt: n=9 vs. cKO: n=10, D), P56 (wt: n=10 vs. cKO: n=6, E), and 3 months (wt: n=5 vs. cKO: n=5, F), mean ± S.E.M. **G:** Antibody labeling against Otx2 or Ribeye as well as DAPI nuclear staining of central or peripheral wt or cKO retinas at P28, P56, and 4 months. **H:** Time course of the number of Otx2+ photoreceptor (PR) rows (10 averaged values per image) in the central or peripheral ONL in wt (n=8, baseline), P28 (n=4), P56 (n=6), 3-month (n=4), and 4-month-old cKO mice (n=4), mean ± S.D. **I:** Antibody labeling against Ribeye of P14 wt (n=4) or cKO (n=3) peripheral retinas with higher magnified insets. **J:** Violin plots of the cellular expression of marker genes of rod photoreceptors in scRNA-seq FACS-purified P7 progenies (n=3 mice per condition, on technical replicate). Mann-Whitney-U-test: *p ≤ 0.05, ** p ≤ 0.01. Scale bars in B: 200 µm, in G, J: 50 µm, insets 25 µm. Layer description see Figure 1.

To measure rod photoreceptor function, scotopic ERGs were conducted (Figs. 4D-F, S6). Wildtypes displayed normal healthy responses, with a scotopic a-wave amplitude of about -200 μV. Scotopic a-wave amplitude recordings in Dicer-cKO_RPC_ mice showed a significant reduction to ∼130 μV at P28, indicating impairment of photoreceptor function. Amplitudes remained unchanged in P56 cKOs mice but further dropped in 3-month-old mice (∼75 μV).

Since the P28 Dicer cKO exhibited no structural impairments in the ONL (unchanged thickness via OCT), all photoreceptors (cell bodies, histology) were generated; however, they apparently were not fully functional (differentiated). Furthermore, the additional functional loss at 3 months of age could suggest the onset of secondary neuronal degeneration, as reported before in an embryonic Dicer loss study (Damiani et al., 2008). This functional loss may be caused by the immaturity of the cells, as no cell loss was detected via OCT.

To analyze photoreceptor impairment at the histological level, we labeled retinal cross sections of various ages with the markers Otx2 (photoreceptor nuclei) and Ribeye (ribbon synapses, Figs. 4G-I, S7A-D). We first evaluated the number of Otx2+ photoreceptor rows in the central and peripheral regions of the retina. While P28 cKO retinas appeared intact, a 26% reduction in the ONL was found at P56 in both center and periphery. However, no further cell loss was evident up to 4 months of age (Fig. 4H), confirming our OCT data.

The evaluation of ribbon synapses showed dense synapses restricted to the OPL in the center and the peripheral wildtype retina (Figs. 4G, S7B) In the P28 Dicer-cKO_RPC_, however, ribbon synapses were more diffuse, not only found in the OPL but also in the ONL. In P56 up to 4-month-old cKO mice, they seemed more and more diffuse and scattered throughout the OPL and ONL, predominantly in the central retina. This suggests remodeling or rewiring events. To determine whether these synaptic impairments were already present at a younger age, we evaluated P14 tissue (Fig. 4I, S7C-D). In the wildtype, Otx2 expression appeared normal, and a clear band of ribbon synapses was formed in the central OPL. In the periphery of Dicer-cKO_RPC_ retinas, Otx2 also seemed to be normal. Ribeye, however, exhibited reduced or absent signals. This suggests that peripheral ribbon synapses were not correctly formed, which may have led to progressive structural impairments and functional defects in the adult retina that spread from the periphery towards the center. Our transcriptomic analysis supported these findings. As mentioned before, the overall proportions of rods in the scRNA-seq datasets were strikingly similar (Fig. 2C). Nevertheless, rod gene transcripts were found reduced in bulk RNA-seq FACS-purified P7 cKO progenies (Fig. 2E, Tables S1). scRNA-seq further confirmed this finding, showing reduced levels of essential rod genes including *Pde6b*, *Rho, Rcvrn, Gnat2, Nr2e3 or Nrl* (Clark et al., 2019, Akimoto et al., 2006, Macosko et al., 2015, Shekhar et al., 2016, Kim et al., 2016a) in the P7 cKO rod progenies (Figs. 4J).

Next, we evaluated cone photoreceptor health using peanut agglutinin (PNA) and M-opsin. The density and pattern of PNA+ and/or M-Opsin+ cone segments in wildtype and P56 or 3-month Dicer- cKO_RPC_ retinas were not different (Figs. 5A-B, S7E). Furthermore, the photopic b-wave in P28 and P56 cKOs was similar to that of wildtypes (about 150 μV), suggesting intact, healthy cones (Figs. 5C-E, S6). Hence, early-born cones were not affected in our Dicer-cKO. In 3-month-old Dicer-cKO_RPC_ mice, however, initial functional cone impairments were detected, possibly as a response to the malfunctioning and/or reduced rods at P56 or due to MG impairment (Larbi et al., 2025), or both.

**Figure 5.**
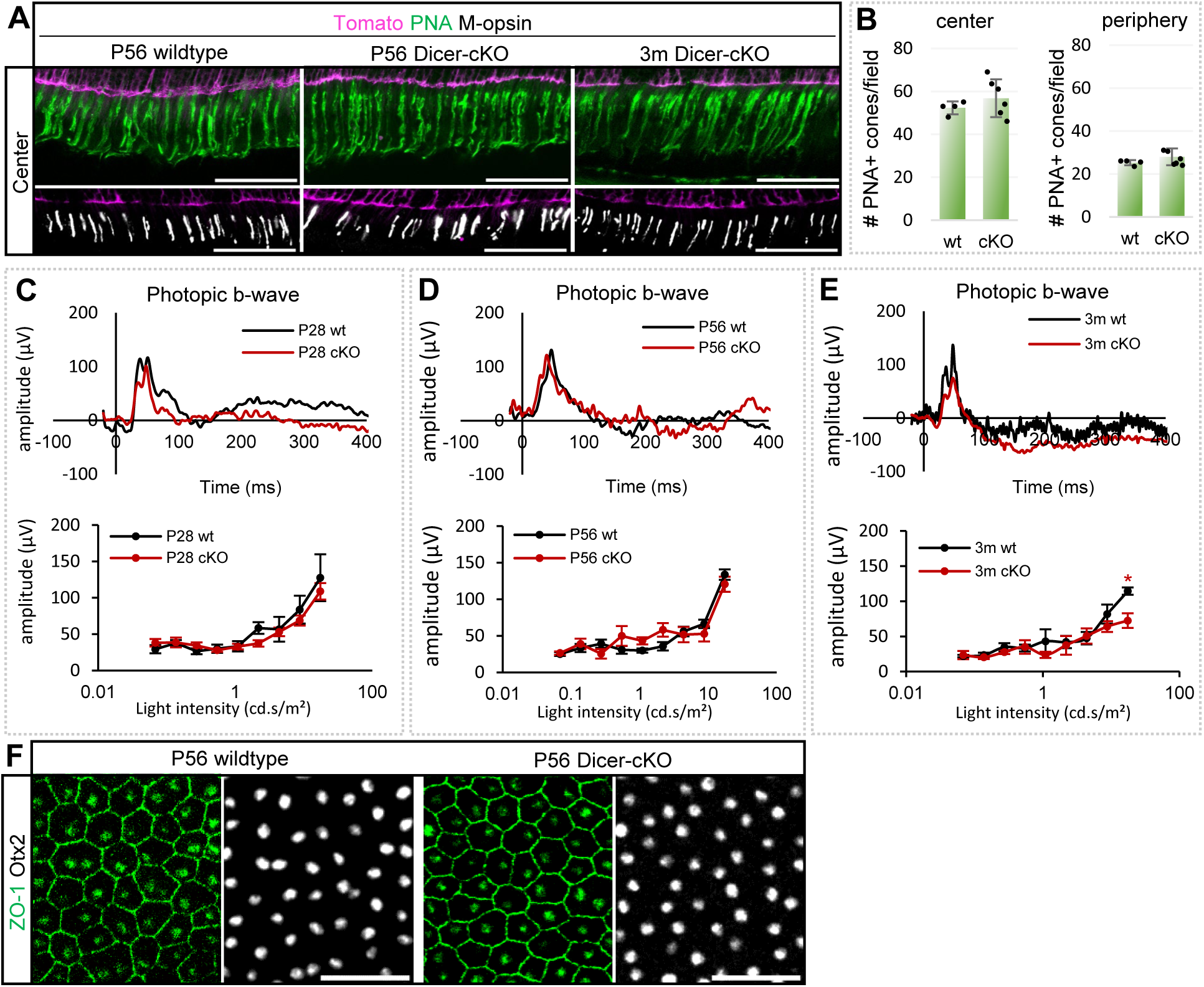
Cone photoreceptors display normal function in young Dicer-cKO mice. **A:** Antibody labeling against peanut agglutinin (PNA) or M-opsin of central P56 wt or cKO retinas. **B:** Absolute number of PNA+ cone segments per field in P56 wt (n=6) and cKO retinas (n=6), mean ± S.D., p= 0.454 and 0.412, respectively. **C-E:** Full-field photopic electroretinogram recordings showing b-wave amplitudes as selected wave forms and intensity dependent graphs for P28 (C, wt: n=9 vs cKO: n= 10), P56 (D, wt: n=10 vs cKO: n= 6), and 3-month old (E, wt: n=5 vs cKO: n= 5) wt and cKO mice, mean ± S.E.M., Mann-Whitney-U-test *: p ≤ 0.05. **F:** ZO-1 and Otx2 antibody labeling of P56 wt or cKO RPE. Scale bars: 50 µm. Layer description see Figure 1.

To exclude the possibility that the RPE contributed to the observed photoreceptor impairment phenotype as seen in RPE-specific (Ohana et al., 2015, Sundermeier et al., 2017) and MG-specific Dicer-cKO studies (Larbi et al., 2025), we examined flat-mounted RPE at P56. We used ZO-1, a marker for the structural and functional integrity of tight junctions (TJs) in the RPE, and Otx2, a marker for RPE nuclei (Fig. 5F). There was no reporter expression found in the RPE of wildtype and Dicer-cKO mice. The RPE of P56 Dicer-cKO mice was intact and similar to wildtype RPE; hence, a direct contribution of the RPE in photoreceptor loss was ruled out.

### Dicer loss in late retinal progenitor cells leads to a reduction in bipolar cell number and functional deficits

We next analyzed BCs, which are also located in the INL and which form between P0 and P7. According to our P7 transcriptomic data, the cKO BC precursor population was increased at the expense of the mature BC population (Fig. 2C), very likely resulting in maturation delays and functional impairments. Furthermore, as mentioned before, BC gene transcripts were reduced in the cKO (bulk RNA-seq, Fig. 2E).

Although OCT scans did not reveal any cell loss in the INL, we evaluated the overall BC population histologically using antibodies against Otx2 and PKC (Figs. 6A-D). In the adult retina, Otx2 is a generic BC marker, while PKC labels specifically rod BCs, which constitute approximately one-third of the overall BC population (Greferath et al., 1990, Wassle et al., 1991). In the wildtype and Dicer-cKO_RPC_ retinas, Otx2+ BCs were found in the lower INL, adjacent to the OPL, in the central and peripheral retina. The vast majority (88%) of all BCs were reporter+, in both areas. However, the total number of Tomato+Otx2+ BCs was reduced in the cKO compared to the wildtype (-26%), and a trend towards reduction was seen in the periphery as well (Figs. 6E-F). 15-20% of all Otx2+ BCs were reporter-negative, possibly due to incomplete reporter recombination, cells not being affected by the Dicer-cKO, or loss of their fluorescence tag. Nevertheless, the decrease in BC progenies resulted in a reduction in the overall BC population in the cKO (center: approximately -26%, periphery: approximately -23%; Fig. S8).

**Figure 6.**
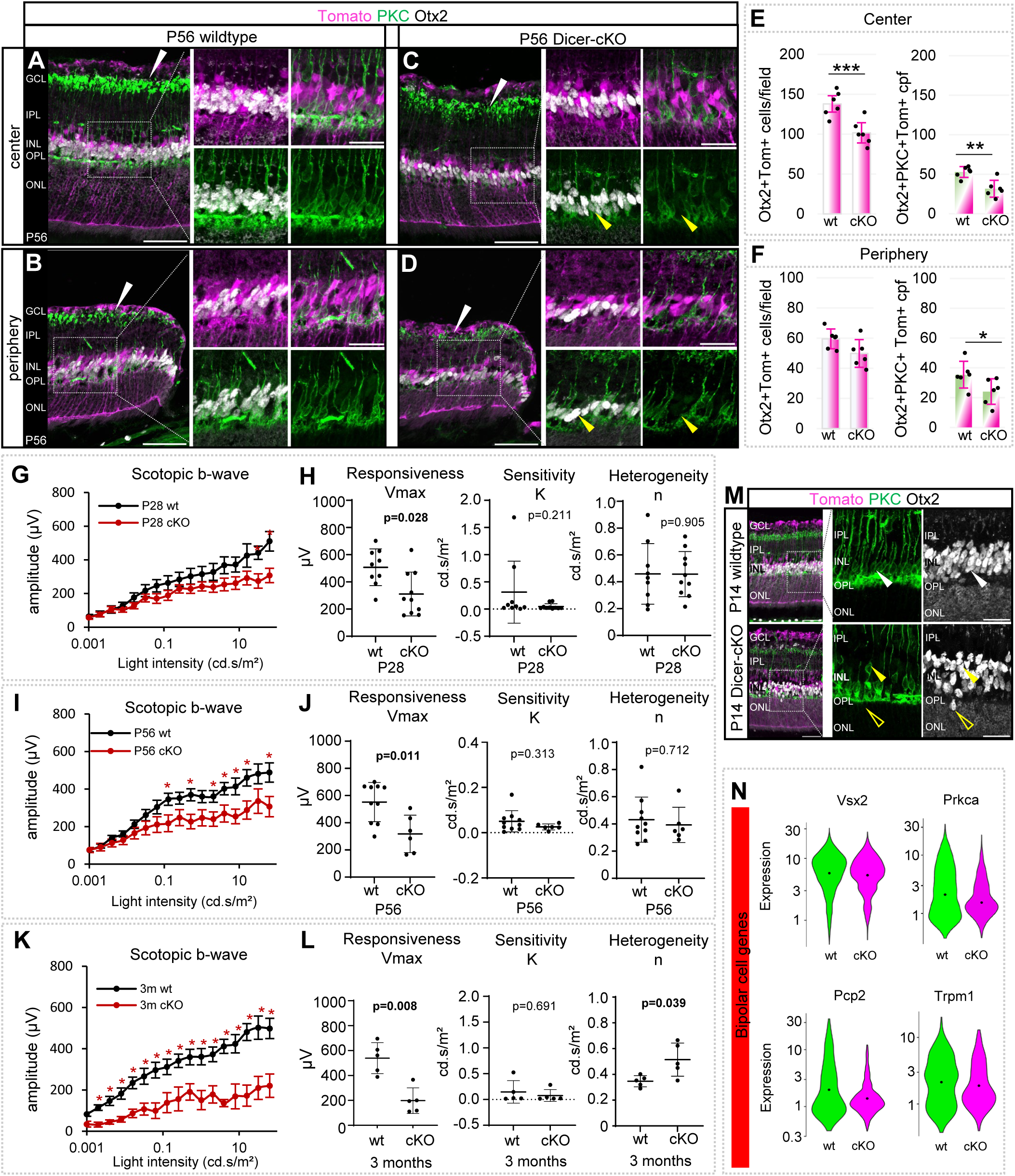
Dicer loss in late RPCs leads to a reduced bipolar cell population and functional defects in the inner retina. **A-D:** Antibody staining to label Otx2+ overall bipolar cells (BCs) and PKC+ rod BCs of central or peripheral P56 wt or cKO retinas, with insets showing higher magnification; white arrowheads show PKC+ dendrites in IPL; yellow arrowheads absent rod BCs. **E-F:** Absolute number of Tomato+ Otx2 BCs and PKC+ rod BCs per field in the central or peripheral retina; wt: n=6, cKO: n=6, mean ± S.D. **G-L:** Full-field scotopic electroretinogram recordings showing b-wave amplitudes of P28 (G), P56 (I), and 3-month-old (K) wt and cKO mice. Estimated saturated amplitudes (Vmax, responsiveness), semi-saturation (K, sensitivity), and slope (n, heterogeneity) using the Naka-Rushton equation for P28 (H), P56 (J) and 3-month old mice (L), P28 wt: n=9 vs cKO: n= 10; P56 wt: n=10 vs cKO: n= 6; 3m wt: n=5 vs cKO: n= 5, mean ± S.E.M. **M:** PKC and Otx2 antibody labeling of central and peripheral P14 wt (n=4) or cKO (n=3) retinas with insets showing higher magnification; white arrowhead shows wt rod BC, yellow arrowheads displaced rod BCs; unfilled yellow arrowheads displaced PKC-negative Otx2+ cells. **N:** Violin plots of the cellular expression of marker genes of BCs in scRNA-seq FACS-purified P7 progenies. Significant differences are indicated, Mann-Whitney-U-test: *: p ≤ 0.05; **: p ≤ 0.01; ***: p ≤ 0.005). Scale bars: A-D, M: 50 µm; insets: 25 µm. Layer description see Figure 1.

The most affected BC type appeared to be the rod BCs. Rod BC somata (PKC+Otx2+) are located in the lower INL. Their processes and dendrites, which are beautifully visualized via PKC, extend to the GCL as well as to the OPL, in both the central and peripheral retina (Figs. 6A-B, white arrowheads). In the P56 Dicer-cKO_RPC_, rod BCs were present, but their number appeared reduced and some were dislocated (yellow arrowheads). Furthermore, their processes appeared less dense and less developed (Figs. 6C-D, white arrowheads). The quantification of the overall number of Tomato+PKC+Otx2+ rod BCs revealed a 40% and 32% reduction in the center and periphery, respectively (Figs. 6E-F). This deficit resulted in an overall rod BC reduction of 42% in the center and 35% in the periphery in P56 Dicer-cKO_RPC_ retinas (Fig. S8).

We next evaluated rod BC function (Figs. 6G-L, S6). In the Dicer-cKO_RPC_, as early as P28, the scotopic b-wave was reduced (∼300 μV vs. wt: ∼550 μV), very likely due to the reduced observed a-wave response. An in-depth analysis using the Naka-Rushton equation revealed a significantly reduced Vmax, i.e., reduced responsiveness. Sensitivity, reflected by the semi-saturation constant (K) as well as heterogeneity, reflected by the slope (n), were, however, surprisingly like wildtype values at P28 and P56 (Fig. 6H, J). In 3-month-old Dicer-cKO_RPC_ mice, the scotopic b-wave amplitudes were approximately 200 μV, indicating a trend toward further functional reduction. 3-month-old cKO mice had a reduced Vmax and a higher slope n, suggesting a potential secondary neuronal degenerative event in the INL at this age, or MG malfunction (Fig. 6L).

To determine whether this phenotype onset was observed early, P14 retinal cross sections were evaluated (Fig. 6M). Overall, P14 wildtype retinas displayed a similar Otx2 and PKC expression pattern as seen in adult wildtypes. P14 cKO retinas, however, had less dense Otx2+ cells (very likely due to cell loss), and some cells were found in ectopic locations, in the ONL. Rod BCs were reduced, and some were misplaced (Fig. 6M, arrowheads) and PKC- (unfilled arrowheads). The PKC+ dendrites in the IPL were less dense/less developed, overall displaying the phenotype seen at P56 and confirming developmental defects. Hence, the aforementioned downregulated BC genes (e.g., *Cabp5)* in P7 cKO progenies (bulk RNA-seq, Fig. 2E) seemed to be the cause of this outcome. scRNA-seq of P7 BC progenies confirmed these findings, showing a reduction of important BC genes, including *Prkca* (encodes for PKC*), Vsx2*, *Trpm1*, *Pcp2,* as well as *Cabp5, Insm1*, and *Gag1* in the cKO BC population. Figs. 6N, S3B).

### P56 Dicer-cKO retinas have increased amacrine cell numbers

Since we found some additional faint Prox1 or Calbindin+ cells in the upper INL in P7 cKOs, we next investigated the ACs in the adult retina. Adult ACs were stained with antibodies against HuC/D and Pax6, both markers that are also in retinal ganglion cells (RGCs) in the GCL (Figs. 7A-D). Similarly, as in P7 retinas, P56 cKO retinas had 1-2 more Pax6+ and HuC/D+ cell layers in the INL (wt center 2-3 Pax6+ HuC/D+ cell layers, wt periphery: 1-2 Pax6+ HuC/D+ cell layers). All HuC/D+ cells were Pax6+, while very few Pax6+ wells were HuC/D-negative and likely MG (Kato et al., 2022, Joly et al., 2011, Bernardos et al., 2007). Many of these ACs were Tomato+ (arrowheads) and may have partially compensated for INL cell loss (BCs), which could explain why OCT scans did not show any reduction in the INL. The quantification of the total number of Tomato+ HuC/D+ Pax6+ ACs in the INL showed a 60% increase in reporter+ ACs in the central cKO retina, and a trend towards an increase (10%) in the periphery compared to wildtypes (Figs. 7E-F). The overall amacrine cell population, however, was stable with a trend towards an increase, probably because the vast majority of ACs (∼67%) were generated before the manipulation.

**Figure 7.**
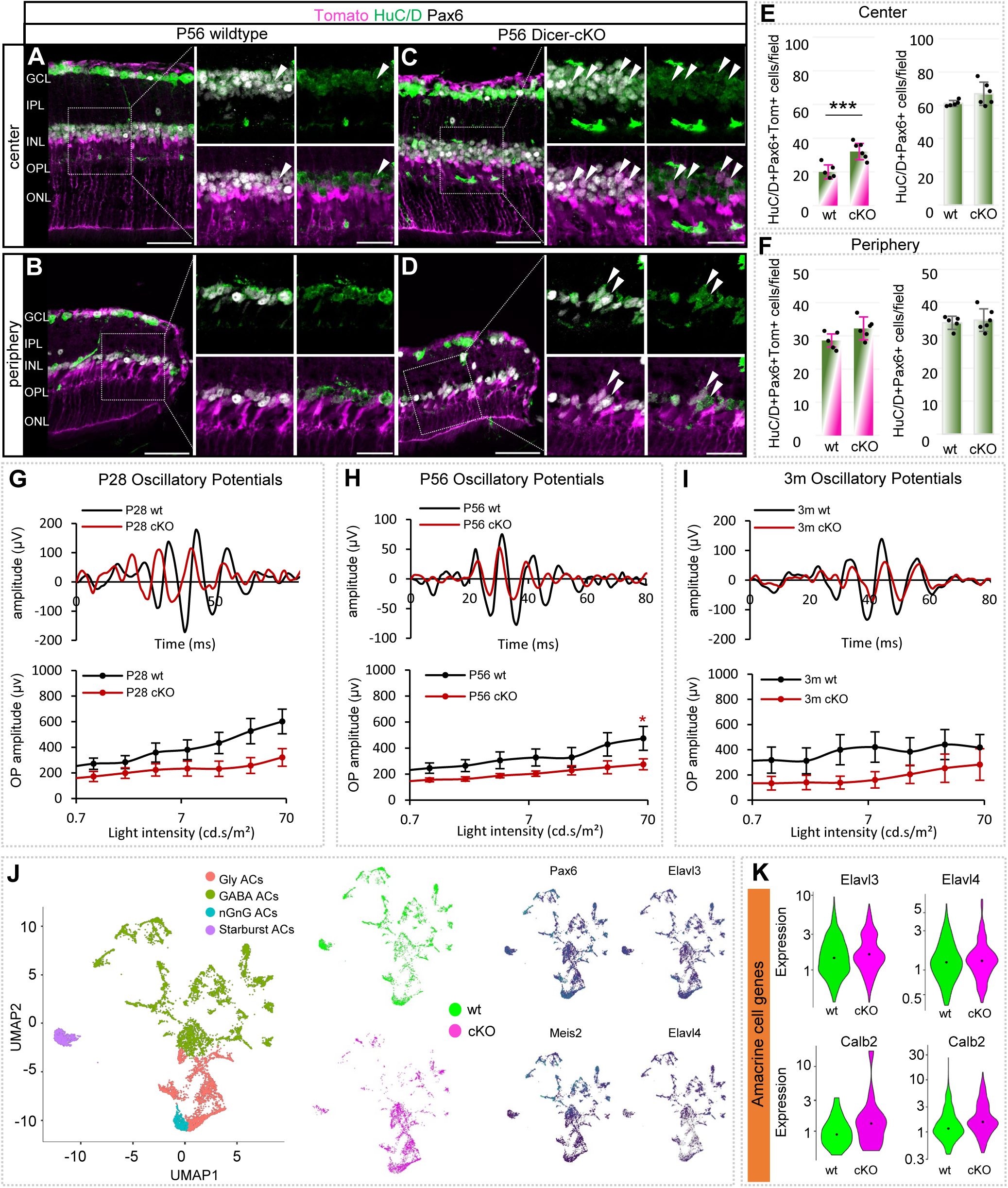
Dicer loss in late RPCs results in an increased amacrine cell population. **A-D:** Antibody staining against HuC/D and Pax6 to label amacrine cells (ACs) in the central or peripheral P56 wt or cKO retina, with insets in higher magnification; white arrows show AC progenies, partially arranged in clusters in the cKO. **E-F:** Absolute number of Tomato+ Pax6+ HuC/D+ ACs and the overall number of Pax6+HuC/D+ ACs per field in the central or peripheral retina; wt: n=6, cKO: n=6, mean ± S.D. **G-I:** Oscillatory potential (OP) amplitudes of full-field scotopic electroretinogram recordings showing selected wave forms and intensity-dependent graphs for P28 (G, wt: n=9, cKO: n= 10), P56 (H, wt: n=10, cKO: n= 6), and 3-month old wt and cKO mice (I, wt: n=5, cKO: n= 5), mean ± S.E.M. **J:** UMAP-dimension reduction of scRNA-seq (one technical replicate) FACS-purified integrated P7 Tomato+ wildtype (WT, n=3) and Dicer-cKO (cKO, n=3) progenies, colored by annotated cell population or cell type as determined by marker gene expression, with representation of selected AC transcripts. **K:** Violin plots of the cellular expression of selected AC transcripts in scRNA-seq P7 progenies. Significant differences are indicated, Mann-Whitney-U-test: *: p ≤ 0.05; **: p ≤ 0.01; ***: p ≤ 0.005. Scale bars in A-D; insets 25 µm. Layer description see Figure 1.

To investigate whether this ectopic cell population impacts retina function, oscillatory potentials (OPs) were evaluated (Figs. 7G-I, S6). OPs reflect inner retinal activity, predominantly neuronal synaptic activity in inhibitory feedback pathways mediated by ACs. They exhibit rapid changes in adaptation and represent both photopic and scotopic processes. The OPs of wildtypes across all ages analyzed were approximately the same, with amplitudes ranging between 400 and 600 μV. The OPs in the cKO mice did not show any significant differences; however, a trend towards a reduced amplitude (300 μV) was observed at all ages. This could suggest possible amacrine cell impairment/rewiring events.

Since we identified some ectopic ACs in our cKOs, we extracted the AC-progeny population from the scRNA-seq dataset to evaluate its composition in the wildtype and cKO (Fig. 7J-K). The major populations found within the P7 wildtype AC fraction based on marker expression (Fig. S9A) included glycinergic (born E16-P1) and GABAergic ACs (born E14-P0) as well as cholinergic Starburst ACs (born E8-P5) and non-GABAergic non-glycinergic ACs (nGnG, born postnatally (Balasubramanian and Gan, 2014), Fig. 7J). Interestingly, the comparison of cell counts and proportions found in cKOs showed alterations in both the absolute and relative numbers (Fig. S9B-C). The predominant cKO AC populations were glycinergic and nGnG ACs (Fig. 7J, S9B-C), both of which were late-born. Furthermore, overall, the GABAergic (early-born) ACs were reduced, but not only within the progenies (tomato+) but also within the non-progenies (tomato-, Fig. S9B). This could suggest not only a direct but also an indirect impairment of GABA-signaling in the cKO. Glycinergic (late-born) ACs, on the other hand, were the dominating cKO AC population. nGnG ACs though, (also late-born) seemed reduced (Fig. S9B-C).

As mentioned earlier, among the genes identified as upregulated in the P7 cKO bulk RNA-seq dataset were the AC genes *Elavl3* and *Elavl4* (Fig. 2E), which encode HuC and HuD, respectively. Not much is known about *Elavl3/4* in the retina. However, *Elavl* genes are highly conserved across species (Perron et al., 1999, Good, 1995, Amato et al., 2005) and play a role in neuronal differentiation, maintenance, and axogenesis in the brain, as well as AC subtype differentiation during retinogenesis for maintaining normal retinal function in the retina (Wu et al., 2021, Wutikeli et al., 2025).

*Elavl3* was expressed in all AC subtypes together with *Pax6*, *Meis2*, and *Calb2* (encodes for Calretinin, Fig. 7J, S9A). It seemed to have somewhat higher levels in the glycinergic and nGnG ACs, though. *Elavl4* was also found across all AC populations. Yet, it appeared to be more restricted to GABAergic ACs, which co-expressed Slc6a1 (encoding GABA transporter 1, GAT1) and Gad1 (encoding glutamic acid decarboxylase (GAD) 67). A quantification of the cell counts and proportions revealed that the major populations among the wildtype progenies were glycinergic (largest), GABAergic, and nGnG ACs (Fig. 7K, S9B). In the cKO, a larger glycinergic but smaller nGnG population was found (Fig. 7K, S9B-C). Therefore, although the overall AC fraction/population in the P7 cKO was similar to that in wildtypes (Fig. 2C), the composition of cKO AC subtypes was different. This suggests a role of Dicer/mature miRNAs in AC subtype specification. Moreover, the overall number of captured ACs in the cKO (Tomato+ and Tomato- cells) was lower than in the wildtype (Fig. S9C). Since we found higher cell numbers histologically, the cKO ACs (even those without the reporter) may be more fragile/impaired and have died during cell sorting. Nevertheless, the evaluation of the AC gene expression in these P7 cKO AC-progenies showed increased levels of *Elavl3 and Elavl4*, as well as *Calb1* and *Calb2* (Fig. 7K), confirming our bulk RNA-Seq data (Fig. 2E)

Since we found ectopic Calbindin+ cells and increased transcripts in the P7 cKO retina (Fig. 1I, 7K), we next evaluated this population in P56 retinas. In the P56 wildtype, Calbindin+ ACs are located in the upper INL, with about 40% of them expressing ChAT (Figs. 8A-B). Calbindin labeling also beautifully visualizes the synaptic interactions in the IPL, which appear as three distinct synapse lines in the central retina (arrows). ChAT+ cells were found in the upper INL and GCL with synaptic connections in the IPL, overlaying the first and third Calbindin synapse layer. In the P56 cKO retinas, similar to the observations made at P7, many Calbindin+ cells were found in the upper INL. These cells were partly Tomato+ but mostly ChAT-negative (Figs. 8C-D, Tomato+ arrowheads, Tomato-unfilled arrowheads). The quantification of the number of Tomato+Calbindin+ cells showed that numbers were doubled/tripled in the central cKO retina, compared to the wildtype (Figs. 8E-F). This led to an overall increase in the Calbindin+ AC population in the upper INL in the cKOs. Calbindin+ ACs, however, constitute only a minority of the overall population (∼20-30%). A cell increase was, however, not observed in the peripheral cKO retina; yet, a trend towards increased Tomato+Calbindin+ ACs and overall Calbindin+ ACs was noted (Figs. 8E-F). Nevertheless, the increased presumptive AC population seen at P7 was still present at P28 (Fig. 8G arrowheads) and remained until P56. The rather loose-appearing processes in the IPL seen at P7 were still present at P28 and P56 and were also ChAT+ (Figs. 8C-D, G).

**Figure 8.**
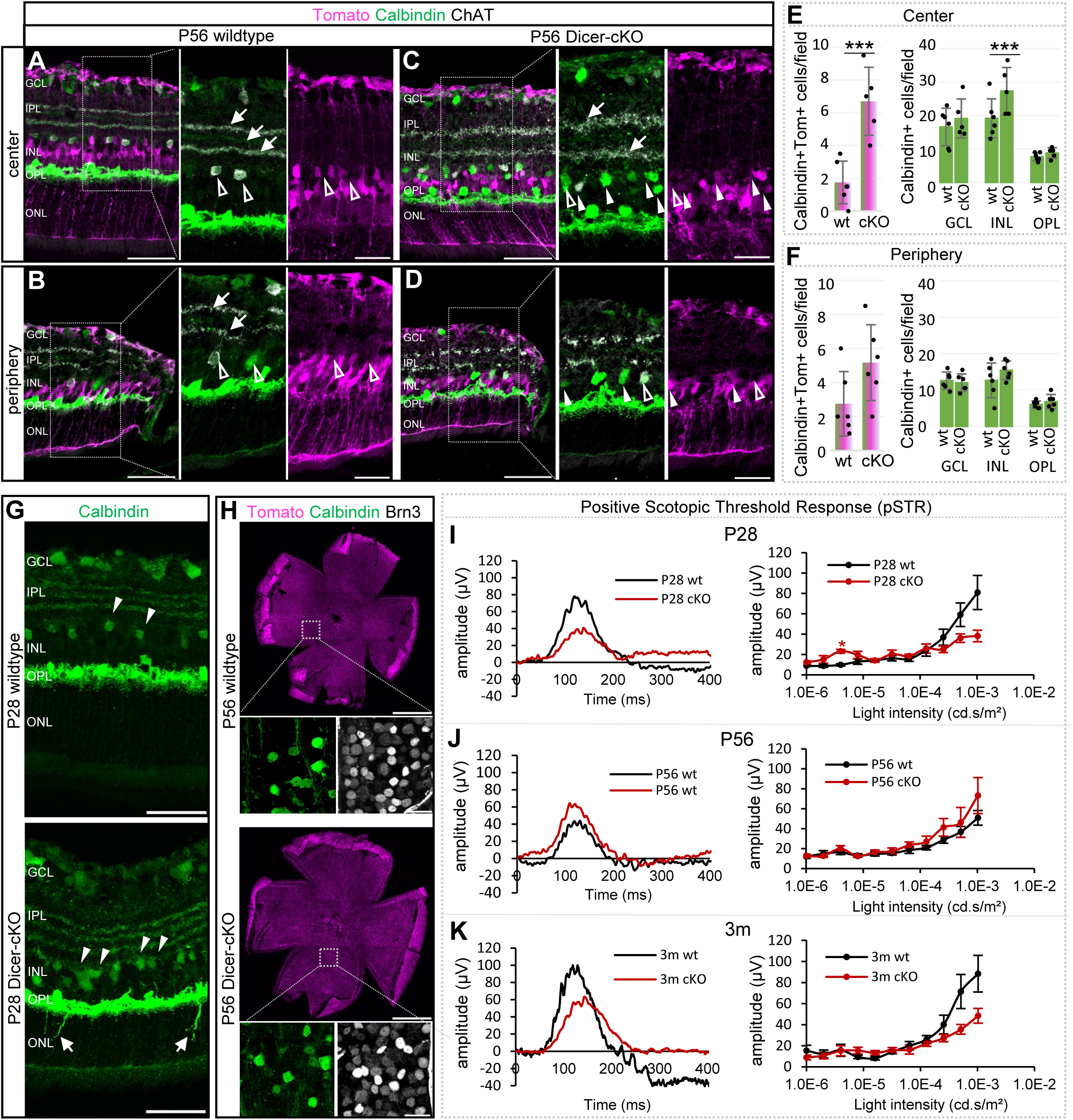
Dicer loss in late RPCs leads to an increased Calbindin+ amacrine cell population but does not affect ganglion cells. **A-D:** Antibody staining against Calbindin and Choline Acetyltransferase (ChAT) to label amacrine cells (ACs) in central or peripheral P56 wt or cKO retinas, with insets in higher magnification; arrows show stratification in IPL, white arrowheads Tomato+ ACs; unfilled arrowheads indicate tdTomato- ACs. **E-F:** Absolute numbers of Tomato+ Calbindin+ cells per field in the INL and the overall number of Calbindin+ cells in the GCL, upper INL, and lower INL/OPL of central and the peripheral retinas; wt: n=6, cKO: n=6, mean ± S.D. **G:** Calbindin labeling of P28 wt (n=4) and cKO (n=4) retinal cross sections showing ectopic horizontal cell neurites (arrows) in regions with high amacrine cell density (arrowheads) in cKO retinas**. H:** Calbindin and Brn3 labeling of retinal flatmounts to visualize ACs and ganglion cells in the GCL of P56 wt and cKO mice, with insets showing higher magnifications. **I-K:** Selected wave forms and intensity graphs of positive scotopic threshold responses (pSTRs) of full-field scotopic electroretinogram recordings of P28 (I, wt: n=9, cKO: n= 10), P56 (J, wt: n=10, cKO: n= 6), and 3-month old wt and cKO mice (K, : n=5 cKO: n=5), mean ± S.E.M. Significant differences are indicated, Mann-Whitney-U-test: *: p ≤ 0.05; **: p ≤ 0.01; ***: p ≤ 0.005. Scale bars in A-D, G,H: 50 µm, insets: 25 µm, for retinal flatmounts in H: 1mm. Layer description see Figure 1.

### Horizontal cells and retinal ganglion cells are not directly affected by miRNA loss in late RPCs

Since Calbindin is also expressed in RGCs and displaced ACs in the GCL, as well as HCs in the lower INL, adjacent to the OPL, we included these cell types in our analysis. RGCs and HC are early-born cell types that should not be affected by Dicer loss in postnatal RPCs/precursors. Indeed, HCs and RGCs were reporter-negative in both conditions, and cell numbers were similar in both conditions (Calbindin+ cell number counts in GCL and OPL see Figs. 8E-F). However, although HCs were reporter-negative, they exhibited some morphological abnormalities, specifically ectopic neurites extending into the ONL (Fig. 8G, arrows). This was particularly evident in areas with a higher AC density, suggesting some neuronal remodeling. A similar phenotype was reported in the *Chx10Cre*: *Dicer-cKO*, which, however, had its onset at embryonic stages (Damiani et al., 2008). On a different note, some of these ectopic Calbindin+ ACs had cell bodies that somewhat resembled the typical MG somata (Fig. 8G)

We next examined RGC pattern and density in retinal flatmounts and used antibodies against Calbindin in combination with antibodies against Brn3 (Figs. 8H). We did not detect any density difference in all 4 quadrants in the wildtype and Dicer-cKO, confirming our observation in cross sections. We concluded that RGCs were not affected (Brn3: wt 105 ± 2 cells/field vs. cKO: 104 ± 3 cells/field, Calbindin: wt: ∼13 ± 4 cells/field vs. cKO: 12 ± 1 cells/field, wt n=3, cKO n=3).

To measure RGC function, the positive scotopic threshold response (pSTR) was evaluated in P28, P56, and 3-month-old wildtypes and Dicer-cKO_RPC_ mice. The pSTR reflects activity in RGCs and a subpopulation of ACs (Lagnado, 1998). At all ages analyzed, we could not detect any significant differences between wildtype and cKO pSTRs (Figs. 8I-K, S6D). There was, however, a trend towards reduced amplitudes in P28 and 3-month-old cKOs, which is likely a result of the already decreased input. The slightly improved waveforms at P56 are unexpected and interesting. This could indicate a possible compensatory mechanism in the cKO at this stage and should be analyzed further in subsequent experiments.

### Possible miRNA regulators of the Dicer-cKO**_RPC_** phenotype

We next aimed to identify possible miRNA candidates that might be involved in the observed Dicer-cKO phenotype. We performed and evaluated the data of small RNA-seq, which includes miRNAs, of FACS-purified P7 wildtype and Dicer-cKO_RPC_ retinas. Dicer deletion results in interrupting the miRNA maturation of most, not all, miRNAs (Cheloufi et al., 2010, Chong et al., 2010, Kim et al., 2016b). Out of a total of 1200 miRNAs found in our dataset, 285 (∼24%) had detectable expression levels in the P7 wildtype. 216 miRNAs out of these 285 miRNAs (∼76%) were downregulated in the cKO (up to 72% reduction), confirming successful Dicer deletion and impairment of miRNA biogenesis. The top 20 P7 wildtype miRNAs and their expression levels in the cKO are shown in a heatmap in Fig. 9A and are listed in Table S2. Among these top miRNAs were the let-7 family (with let-7b having the highest expression levels), as well as miR-183 and miR-204. We next used the Diana Tool microT to identify predicted targets of the top 20 miRNAs. We found about 12,660 hits with an interaction score of 0.7-1. We then compared this list of predicted targets of the top 20 downregulated miRNAs in the P7 cKO with the list of upregulated genes in the P7 cKO (at least 20% increase). We found 164 matches. Gene Ontology (https://pantherdb.org) identified the biological processes in which these genes are involved (Fig. 9B, Table S3). Major processes included synapse formation, signaling, and axon guidance, which might explain some of the observed differentiation deficits of differentiating neurons. Nucleokinesis and radial glia migration/cell motility were also prominent biological processes and a possible explanation for the displaced Sox2+/Sox9+ cells in the cKO. Processes related to photoreceptor and amacrine cell differentiation were also found, which could be the reason for the robust rod cell status in the cKO and the ectopic AC population we found. The genes associated with these major processes and their targeting miRNAs were *Sun1* (Sad1 And UNC84 Domain Containing 1) and *Syne2* (Spectrin Repeat Containing Nuclear Envelope Protein 2), involved in glial/nuclear migration. They are regulated by miR-181c/182/484 (Fig. 9C, Table S1). *Sox9* and *Casz1* (Castor zinc finger 1) were associated with photoreceptor differentiation and are targeted by the let-7 family and miR-151, respectively. *Hes1* and *Casz1* again play a role in ACs differentiation. *Hes5* is regulated by miR-182 (Fig. 9C, Table S1). Notably, *Casz1*, which appeared in two different processes, is a well-studied gene. It not only regulates photoreceptor differentiation but also maintains late RPC identity (Mattar et al., 2015, Mattar et al., 2021, Mattar et al., 2018). *Sox9* and *Hes1* are key factors in embryonic and postnatal brain/retinal development, as well as essential for glial development (Poche et al., 2008, Zhu et al., 2013, Tomita et al., 1996, Kageyama et al., 1997, Furukawa et al., 2000, Takatsuka et al., 2004, Wohl, 2024).

**Figure 9.**
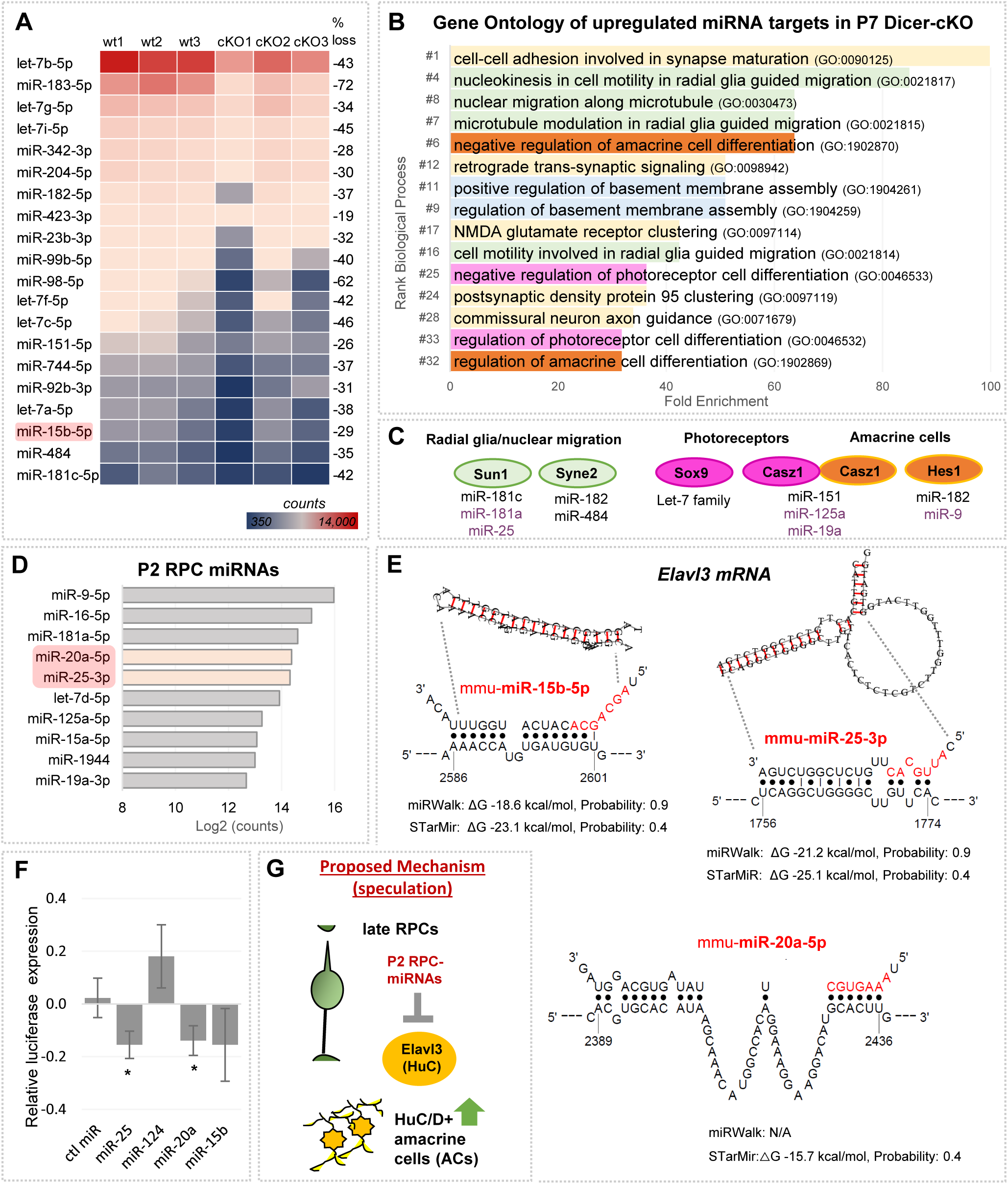
Processes regulated by RPC miRNAs include amacrine cell genes. **A:** Heatmap of the top 20 miRNAs highly expressed in FACS-purified P7 wildtype progenies and their expression levels in the cKO, with emphasis on miR-15b (small RNA-seq, n=3 per condition). **B:** Biological processes (Gene Ontology, GO, fold enrichment) of genes targeted by RPC-miRNAs found upregulated in P7 cKO FACS-purified progenies (>20%, bulk RNA-Seq dataset). **C**: Selected genes associated with major GO biological processes and their targeting P7 and P2 (purple) RPC miRNAs. **D:** miRNAs highly expressed in P2 RPCs (log2 counts from *Wohl et al., 2019*), not found in the P7 miRNA dataset with emphasis on miR-20a/miR-25. **E:** Predicted binding sites for miR-15b-5p, miR-25-3p, and miR-20a-5p in the 3’UTR of the *Elavl3* mRNA using STarMir and miRWalk. miRNA seed sequences are indicated in red. Guide values for ΔG (hybrid stability): < -15 kcal/mol, probability: ≥ 0.5 indicate high hybrid stability/probability. **F:** Normalized relative luciferase activity of *Elavl3* 3’UTR reporter plasmids co-transfected with control miRNAs (ctl miR) or RPC mimics, 3 technical replicates, 6 independent experiments, mean ± S.D., Mann-Whitney U-Test: *: p <0.05. **G:** Schematic of proposed possible mechanism for AC overproduction: loss of late RPC-miRNAs miR-20a/25 might result in an increase of *Elavl3* mRNA leading to ectopic HuC+ ACs.

A somewhat unexpected finding was that many miRNAs known to be expressed by RPCs/PC, including miR-9, miR-16, miR-25, miR-20a, miR-15a, and miR-19a (La Torre et al., 2013, Genini et al., 2014, Wohl and Reh, 2016, Hackler et al., 2010) were not detectable in our P7 dataset (or only at very low levels, Table S4). It was reported several years ago that P0-P2 RPCs, including miR-16, 15a/b, and miR-25, decline rapidly within the first postnatal days (Hackler et al., 2010). However, this fact could also be due to the different methodology used and/or RNA degradation, as our yields of captured miRNAs were relatively low (25%). To rule out the possibility of missing essential miRNA candidates that might play a role in the cKO phenotype, we plotted the top 10 miRNAs found in P2 RPCs (Fig. 9D, Table S4). Interestingly, several of them also target *Sun1, Casz1*, and *Hes*2 (in Fig. 9C, shown in purple font), among them miR-25. Interestingly, miR-25, together with miR-15 and other P2 RPCs miRNAs were shown to induce RPCs features in primary MG when overexpressed (Wohl et al., 2019). These RPCs subsequently adopt BC features, but not AC features. Since our RPC-miRNA loss leads to an almost opposite outcome, we hypothesized that these P2 miRNAs might be involved in the observed BC/AC phenotype.

One major AC gene that caught our attention was *Elavl3,* which encodes HuC. HuC, together with HuD, is only expressed in RGCs (which are not affected in our model) and ACs, but not in BCs, HCs, or MG. Furthermore, *Elavl3* was found to be upregulated in P7 cKO progenies (bulk and scRNA-Seq) and may be a contributing factor to the higher number of HuC/D+ ACs observed in the cKO. To explore the possibility that P2 RPC-miRNAs target Elavl3 mRNA, we utilized MirWalk and STarMIr, two comprehensive mRNA:miRNA target prediction databases. These two tools, based on AGO-Clip data, provide extensive information about and visualize the binding sites (Figs. 9E, Table S5). Diana Tools and TargetScan were also used, although they do not utilize the most current mRNA transcripts. We identified three P2 miRNAs, namely miR-15b, miR-20a, and miR-25, which are predicted to target *Elavl3*. The predominant binding sites were at the 3’UTR, but also sites at the CDS (coding sequence, Figs. 9E, S10, Table S5, ΔG should be less than -15 kcal/mol and the probability for binding should be higher than 0.5 and close to 1 (Rennie et al., 2014)). To test whether these predictions are resulting in genuine miRNA:mRNA interaction, we performed dual luciferase assays. We transfected miR-25-3p, miR-20a-5p, miR-15-5p, as well as miR-124-3p as a reference miRNA (Lu et al., 2021), or control mimics, together with a dual luciferase plasmid containing the 3’UTR of *Elavl3* mRNA. We found that miR-25 and miR-20a significantly reduced luciferase activity. miR-15b showed a trend towards reduction (Fig. 9F). This supports our hypothesis that the reduction/ loss of these miRNAs around P2-P4 via Dicer loss could have contributed to the increase in their target *Elavl3* mRNA (Fig. 9G).

## Discussion

In the past decades, a variety of studies showed that Dicer and proper miRNA biogenesis are required for interspecies developmental processes of many organs and systems, including the central nervous system (De Pietri Tonelli et al., 2008, Davis et al., 2008, Kawase-Koga et al., 2010, McLoughlin et al., 2012, Zindy et al., 2015, Rajaram et al., 2014, Konar et al., 2020, Lagos-Quintana et al., 2002, Decembrini et al., 2008). In the neural retina, miRNAs have been shown to play a fundamental role during the embryonic phase, regulating the competence of early RPCs and the fates of their progenies (Davis et al., 2011, Georgi and Reh, 2010, La Torre et al., 2013, Iida et al., 2011, Pinter and Hindges, 2010). Whether Dicer and miRNAs also play a role in postnatal retinogenesis has not been explored *in vivo* yet.

Here, we demonstrate that Dicer in postnatal RPCs/PCs is essential for ensuring the proper maturation and differentiation of all late-born progeny. Overall, we observed five primary outcomes: a reduction in BCs (1) and MG number (2), impaired rod maturation and function (3), the presence of an ectopic immature population (4), and an ectopic AC population (5). Although somewhat similar observations regarding BC, rods, MG, and immature cells have been reported before, to our knowledge, AC overproduction has not been documented in any of the embryonic RPC Dicer-cKO studies. This cell addition, however, may be one of the reasons for the relatively slow degeneration we observed in the adult retina. We identified *Elavl3,* which encodes HuC, and miR-20a/25 as possible key players and speculate that their interaction may be a possible regulatory mechanism underlying this AC phenotype. We also present and discuss additional possible key players that may be involved in the observed maturation delays affecting central and peripheral progenitor/precursor cells and their progeny.

### The vital role of Dicer in embryonic and postnatal retinogenesis

Dicer and miRNAs are crucial during the embryonic stages of retinogenesis. The first study was conducted by the Strettoi and Harfe Labs (Damiani et al., 2008), using the *Chx10-Cre* mouse (Rowan and Cepko, 2004). *Chx10*, also known as *Vsx2*, is induced early, around E11.5 (Belecky-Adams et al., 1997, Burmeister et al., 1996, Green et al., 2003, Rowan and Cepko, 2004, Sigulinsky et al., 2008). Although this manipulation did not cause a severe phenotype due to mosaic expression of *Chx10-Cre*, which was not traceable, one notable abnormality was the temporary formation of rosettes at P16. These rosettes appeared to cause dysfunction in rod and cone photoreceptors and ongoing degeneration in adult mice (up to 5 months). Signs of retinal remodeling, such as neurite sprouting of HCs, were observed, similar to our findings. Using the inducible Ascl1CreER transgenic mouse (Kim et al., 2011, Brzezinski et al., 2011), we induced Dicer loss after birth, specifically targeting postnatal RPCs/PCs. Unlike Damiani et al., we identified only functional deficits in rod photoreceptors (not cones) in young adult mice. Cone dysfunction was only seen later on, indicating secondary impairment due to rod malfunction (Gargini et al., 2007) and/or MG dysfunction (Larbi et al., 2025). We did not see rosettes; however, HC neurite sprouting suggests similar remodeling mechanisms are present in both embryonic and postnatal Dicer- cKO models.

A second RPC Dicer-cKO study was conducted at the embryonic stage (E10.5) using the *αPax6Cre*:R26EYFP mouse. This model allowed for a more detailed investigation of Dicer-cKO cells through lineage tracing (Georgi and Reh, 2010). In this mouse, only peripheral RPCs lost Dicer, enabling extensive tissue analysis into adulthood. Unlike Damiani et al. (2008), a notably severe and interesting phenotype was observed in the affected periphery: early-born RGCs and HCs were produced in abundance, whereas late-born cell types (rods, MG, and BCs) were not formed. Late RPC genes, including Ascl1, were downregulated, and cell proliferation was slightly reduced at E16 and P1 (Georgi and Reh, 2010). This contrasts with our findings in postnatal RPCs, where we observed an increase in *Ascl1* transcripts and cell proliferation. This suggests that the specific progenitor state at which Dicer is deleted is essential, as that particular cell state seems to be maintained or extended. Additionally, embryonic and postnatal RPCs express somewhat different sets of miRNAs (Hackler et al., 2010), which likely regulate different genes (targets) and biological processes, including RPC state and competence (La Torre et al., 2013)).

A subsequent embryonic Dicer-cKO study, conducted by the Ashery-Padan Lab (Davis et al., 2011), confirmed the overproduction of RGCs at the expense of late-born retinal cells (HCs, PRs, BCs, ACs, MG), as seen by (Georgi and Reh, 2010). Davis et. al., focused on neuroepithelial cells, including those of the iris and ciliary body, and used different *Cre* lines, including the αCre that labels optic cup progenitors (E10/11). Interestingly, similar to our findings, an increase in proliferation markers (i.e., Ki67, PH3, Ccnd1) and Sox2+ cells (a progenitor marker) was observed, resulting in columns of progenitor cells that persisted until P8 (Davis et al., 2011). An ectopic progenitor population was also found within the ciliary margins. Although our ectopic immature population was not restricted to the periphery and the ciliary marginal zones in our Dicer-cKOs did not show any abnormalities, these outcomes are comparable. This may suggest shared mechanisms for maintaining cells in an immature state. Interestingly, Davies et al. reported a differentiation failure of ACs and PRs in the *αCre-*cKO at E18.5, just before birth. This was due to the absence of Syntaxin1 and altered expression of Bhlhe22 and Tfap2a. Both genes are found in differentiating ACs (Jin et al., 2015, Park et al., 2023, Perez-Leon et al., 2022). However, Foxn4, a marker of cycling AC and HC precursors (Li et al., 2004), and Ptf1, a marker expressed in postmitotic AC precursors (Fujitani et al., 2006), were present (Davis et al., 2011). This led the authors to conclude that miRNAs are dispensable for AC type specification but essential for their differentiation (Davis et al., 2011). In our study, all later-born AC types were present, but their proportions were altered. We also observed additional, ectopic Calbindin+Pax6+HuC/D+ ACs, which persisted at least until P56. This suggests some differences related to the RPC stage at the time Dicer is deleted (continued discussion below).

A very severe phenotype, with most cells gone by P14, was reported by the Watanabe Lab using the *Dkk3-Cre* mouse (Iida et al., 2011). *Dkk3-Cre* is active during the embryonic stage (E10.5) (Sato et al., 2007), but throughout the entire retina. As a result, massive RPC death during embryonic phases was found, leading to microphthalmia. GCL and IPL were not formed. However, specific differentiation markers were expressed, including the MG/astrocyte marker GS (scattered throughout the Dicer-cKO retina). This is interesting since *Dkk3* is known to be expressed in MG (Gallina et al., 2016, Sato et al., 2007, Zhu et al., 2018), hence one would expect them to be affected most. Since that was not the case, MG progenitors/precursors appear to be relatively resilient cell types and may be less affected by Dicer/miRNA loss than neuronal progenitors/precursors. This could partly explain why also in our study, MG were formed. However, Dicer loss in young MG leads to MG impairment, which, nevertheless, takes several months to become evident (Larbi et al., 2025, Wohl et al., 2017).

Nonetheless, cells most affected in the *Dkk3-Cre: Dicer-cKO* were BCs (rod BCs), showing weak PKC expression both *in vivo* and *ex vivo* (after electroporating P0 retinas and maintaining them in culture for 12 days (Iida et al., 2011)). Since we too found impaired BCs with predominantly impaired PKC+ rod BCs, this suggests that BC progenitors/precursors are more susceptible to Dicer/miRNA loss. The P0 explants also displayed an unchanged rod number, consistent with our findings, a slight increase in MG number, and a ∼66% reduction in Islet+ cells (RGC/ACs). Impaired ACs, indicated by weak Pax6/HuC/D expression, were also present *in vivo* (Iida et al., 2011). Hence, ACs seem to be affected differently in the *Dkk3-Cre* study. A possible explanation could involve the *Cre* lines used. Although *Dkk3* and *Ascl1* are both expressed in RPCs, some MGs, and mature ACs (Muranishi and Furukawa, 2012), their expression patterns do not appear to overlap. This is likely due to their opposing roles, as Ascl1 inhibits Dkk3, as demonstrated during fish regeneration (Ramachandran et al., 2011). As a result, different subpopulations of RPC/precursors may have been manipulated in the Iida et al. study and our study. Additionally, the Dicer phenotype can be accelerated *ex vivo* (Wohl et al., 2017); therefore, a culture experiment does not necessarily reflect the same timeline observed *in vivo*.

### RPC miRNAs and RPC state - peripheral versus central retinal impairments

One of the other significant findings in our study was a delay in tissue maturation. This was very likely due to increased and/or sustained transcripts associated with the cell cycle (more Ki67+ PH3+ cells in cKOs). A key miRNA regulator that plays a fundamental role during development by controlling cell cycle dynamics is let-7. let-7 (lethal-7) is a family of miRNAs consisting of let-7a, b, c, d, e, f, g, and i, which are expressed in mouse RPCs, MG, and neurons (La Torre et al., 2013, Wohl and Reh, 2016). During retinal development, let-7 accumulates in RPCs, and its levels and activity oscillate, mediating cell cycle exit in a concentration-dependent manner (Fairchild et al., 2019). One well-known target of let-7 is CyclinD1 (Shu et al., 2012, Song et al., 2020, Tu et al., 2022, Wang and Cai, 2020), Cyclin D1 is encoded by the *Ccnd1* gene, a transcript we found upregulated in the cKO mouse. Ccnd1 is a major regulator of cell division and is downregulated during cell cycle exit and retinal cell maturation (Barton and Levine, 2008, Trimarchi et al., 2008, Dyer and Cepko, 2000a). Alterations in *Ccnd1* have been shown to significantly effect cell fates, and consequently the cellular composition of the retina (Das et al., 2009, Das et al., 2012). Besides regulating the RPC cell cycle, let-7 also appears to play a role in primary mouse MG proliferation events (Wohl et al., 2019). Therefore, the decline of members of the let-7 family might have contributed to the prolonged cell cycle events we observed. Furthermore, in addition to cell cycle control, let-7 also negatively regulates Ascl1 expression. That occurs in RPCs (La Torre et al., 2013) as well as in fish MG after initiating on a natural regeneration program in response to injury (Ramachandran et al., 2010, Goldman, 2014, Konar et al., 2020). An induction of Ascl1 was also observed in the aforementioned primary MG study after let-7 inhibition (Wohl et al., 2019). Since the P7 cKO-progenies in our study displayed increased Ascl1 transcripts, the loss of let-7 might be responsible for this outcome as well.

Because the retina matures and differentiates from the center to the periphery, we conducted thorough analyses of both central and peripheral regions. Compared to findings from embryonic studies (Georgi and Reh, 2010, Iida et al., 2011), our results were somewhat unexpected, such as the still relatively normal size of the overall retina, which meant no drastic cell loss of the affected cells; a normal OCT, even though this was misleading since the cellular composition differed, and ectopic immature and AC populations also found in the central retina. Moreover, despite being impaired, the retina was still functional for at least up to 4 months of age in the mouse. These are fascinating results that will require further investigation, and we can discuss them here only to a limited extent. Part of the explanation might, however, lie in the cell state of the affected cells.

Furthermore, although most Tomato+ progenies were identified as proliferating RPCs, we also affected more committed precursor cells, especially in the central retina (Young, 1985b, Kageyama et al., 1997, Belecky-Adams et al., 1997, Hojo et al., 2000, Turner and Cepko, 1987, Cepko et al., 1996). As a result, these BC, rod, MG, and AC precursor cells may respond differently to Dicer/miRNA loss than late RPCs, leading to even more diverse outcomes. It appears that the impact of Dicer/miRNA loss on RPCs is faster than in more mature cells. This is probably because RPC miRNAs fluctuate and can change within a few days (Hackler et al., 2010). Hence, their rapid turnover might enhance the Dicer-cKO phenotype. Conversely, miRNAs of specific differentiating cells appear relatively stable, at least under normal (injury-free) conditions. This may allow survival for several months, even if most cells are affected (Sundermeier et al., 2014, Wohl et al., 2017, Sundermeier et al., 2017, Aldunate et al., 2019). Therefore, the stability/half-lives of Dicer (both protein and mRNA), as well as miRNA, are also influential. This means that, although Dicer was impaired by P4, the mRNA and protein produced up to that point remained present and functional. The half-lives of miRNAs are very considerable, ranging from hours to weeks, especially when bound to other factors like argonaut proteins (Grosshans and Chatterjee, 2010, Gantier et al., 2011, Winter and Diederichs, 2011, Perez-Ortin et al., 2013, Kingston and Bartel, 2019). Consequently, not only the miRNA itself but also its half-life can be cell-type specific (Kingston and Bartel, 2019). As a result, the dynamics of miRNA may differ across retinal cell types, leading to distinct outcomes. Moreover, there is a Dicer-independent miRNA pathway (Cheloufi et al., 2010, Yang and Lai, 2011). Although only a small number of miRNAs are processed this way, these miRNAs could still influence the canonical pathway.

Another aspect we want to address is the immature Sox9+Sox2+GS-negative population found in both the peripheral and central cKO retina. While the peripheral cells might indeed be remaining RPCs that stay undifferentiated, the central cells could belong to a different population. Although we observed these cells at P7 and P14, they may already have been more committed MG precursor cells, possibly even young glia, by the time the miRNA loss occurred. Furthermore, some of the ectopic ACs seen in the central retina could even be cells that originated from impacted MG precursors. This is because some of their nuclei were somewhat triangular and MG-like rather than round. Furthermore, Dicer/miRNA loss in P11 MG leads to proliferation and migration of the MG, which also lose GS expression over time (Wohl et al., 2017). This could suggest a similar mechanism. The impairment of MG precursor/young glia could also be part of the events resulting in the relatively moderate degeneration we observed. It is conceivable that the glial reactivity of young glia in response to cell loss leads to less secondary neuronal loss.

### miRNAs of postnatal RPCs as possible regulators of amacrine cell populations

As mentioned earlier, another major outcome of our postnatal RPC/PC study was the ectopic AC population, a result not previously reported. We therefore decided to focus on these ACs, not only because it provides a more straightforward way to analyze upregulated genes in a Dicer-cKO and identify direct miRNA targets, but also because the relatively slow degeneration we observed in the adult retina may indicate the presence of compensatory mechanisms. Additional neurons might explain this phenomenon (neuroplasticity).

So, what is causing the generation of these additional ACs? We do not have a complete answer to that question yet, and it is very likely a complex event involving many factors and miRNAs. For instance, early factors such as Pax6, Sox2, Meis2, and Cdkn1c (which encodes p51kip2), all found to be upregulated in P7 cKO progenies, are known to influence the competence of retinal progenitors and significantly affect the development of ACs from the early progenitor pool. (Balasubramanian and Gan, 2014, Remez et al., 2017, Bumsted-O’Brien et al., 2007, Conte et al., 2010, Dupacova et al., 2021, Yan et al., 2020, Dyer and Cepko, 2001a, Dyer and Cepko, 2001b, Dyer and Cepko, 2000b). In particular, Pax6 and Meis2 are required for the generation of late-born glycinergic AC types (Marquardt et al., 2001, Remez et al., 2017, Bumsted-O’Brien et al., 2007, Oron-Karni et al., 2008). Meis2 is also an early marker for postmitotic GABAergic ACs (Bumsted-O’Brien et al., 2007), which is targeted by miR-204 (Conte et al., 2010). miR-204 was found in the P2 and our P7 progeny population, with declining levels in the cKO. Hence, the increase of ACs might also be partially regulated via the miR-204 and Meis2 (Conte et al., 2010).

Sox2, expressed in cholinergic and GABAergic subsets of ACs in the inner INL of the adult retina (Li et al., 2021), also influences AC fates. Overexpression of Sox2 causes progenitor cells to shift toward an AC fate (Li et al., 2021), while its knockdown results in a switch from amacrine to bipolar fates (Li et al., 2021). Since we observed an increase in Sox2 and ACs, but a loss of BCs, Sox2 may act as an upstream regulator promoting AC overproduction at the expense of BC development. p57Kip2 was shown to regulate AC proportions and appears involved in determining GABAergic AC fates. Loss of p57Kip2 led to an increase in Calbindin+ ACs, although this p57Kip2-expressing AC population does not colocalize with markers like Calbindin, Calretinin, ChAT, or others (Dyer and Cepko, 2001b). Since we found that the transcripts of these genes increased in our P7 cKO bulk RNA-seq data, they may contribute to the observed phenotype, potentially under the regulation of RPC miRNAs.

Nevertheless, we would like to propose another possible mechanism, introducing new potential key players: miR-20a/25/15 and Elavl3. This remains speculative at this point, but it also provides a thought-provoking starting point for further in-depth analysis. Elavl3 encodes HuC and was found to be expressed across all AC populations and upregulated in cKO progenies. Furthermore, the ectopic AC population in the P56 retina was HuC/D+. HuC/D is exclusively expressed in ACs and RGCs, but not in HCs, BCs, or MG. Because of that, and since RGCs are unaffected in our model, this marker serves as a relatively specific AC marker in our study. Additionally, Elavl3 remains a relatively unexplored gene, despite initial studies being conducted over two decades ago. It is known that the Hu family of proteins contains RNA-binding proteins related to the Drosophila protein ELAV (embryonic lethal abnormal visual system) and are essential for nervous system development and maintenance (see review (Soller and White, 2004, Perrone-Bizzozero and Bolognani, 2002).

There are four Hu family members in vertebrates with HuB, HuC, and HuD being neuron-specific, encoded by *Elavl2*, *Elavl3,* and *Elavl4,* respectively (Amato et al., 2005). These factors are first expressed in the GCL and later in the inner part of the INL, which contains ACs (Good, 1995, Perron et al., 1999). *Elavl* genes play a vital role in neuronal differentiation, maintenance, and axogenesis in the brain (Yokoi et al., 2017, Mulligan and Bicknell, 2023, Bronicki and Jasmin, 2013). Recently, *Elavl* genes have been reported to be involved in AC subtype differentiation during retinogenesis to maintain normal retinal function (Wu et al., 2021, Wutikeli et al., 2025). Therefore*, Elavl3* might be involved in AC fate, as well as overall tissue maturation processes. Two of the targeting miRNAs identified and validated via luciferase assays in this study are miR-20a and miR-25. These are miRNAs known to be expressed in P2-4 RPCs (Hackler et al., 2010, Wohl et al., 2019). Furthermore, overexpression of these RPC-miRNAs (in cocktails or separately) in young primary MG predominantly drives BC fates, but not AC fates (Wohl et al., 2019). ACs are, however, the other primary cell type usually generated in MG reprogramming experiments (Ueki et al., 2015, Yao et al., 2016, Le et al., 2023). This led us to hypothesize that these miRNAs might influence BC/AC fate determination. Although *Elavl3* hasn’t been examined in these reprogramming studies, it could be a direct target of miR-20a and miR-25. This regulation isn’t limited to the 3’UTR. We also observed binding sites for these miRNAs in the coding sequence (CDS), which results in the inhibition of *Elavl3* translation (Tastsoglou et al., 2023, Brummer and Hausser, 2014). Additionally, it’s important to note that there are multiple binding sites for miR-25. One appears to be a target-directed miRNA degrading site (TDMD site), leading to the degradation of the miRNA itself (Buhagiar and Kleaveland, 2024). Therefore, other regulatory events might also occur. To better understand these complex interactions, further experiments will be necessary.

Taken together, late RPC Dicer/miRNAs seem to regulate processes essential for the proper differentiation and maturation of all three major late-born cell types: rods, BCs, and MG. The loss of miRNA primarily affected rod BCs; therefore, BC progenitors and precursors appear to be the most vulnerable, as observed also during the embryonic stage. In contrast, rod and MG progenitors and precursors appear to be quite resilient. AC progenitors and precursors, however, show opposing reactions depending on the timing of Dicer/miRNA loss and appears beneficial in the postnatal retina. We speculate that a potential partial explanation for the observed AC phenotype might involve *Elavl3*, which encodes HuC and is targeted by miR-25/20a. Overall, Dicer and mature miRNAs play a vital role in maintaining and ensuring the survival of not only embryonic but also postnatal RPCs and their progeny in the adult retina.

### Limitations of the study

This study provides the first comprehensive report on postnatal RPC/PC miRNA loss, combining structural and functional in vivo analysis with histology, bulk RNA, and single-cell RNA sequencing (scRNA-seq). The main goal was to identify molecular changes in early postnatal late RPC/PC progeny and to assess how these changes affect overall retinal health, including structure and function, in adult mice. Extensive transcriptomics analyses were performed at P7, a relatively early stage. We chose this time point intentionally, not only to confirm the observed phenotypes but also to identify potential key players and mechanisms. Additional transcriptomics at later stages are beyond the scope of this study. Moreover, many potential regulators were identified and partially introduced. To validate these putative key regulators and further understand their networks, future studies involving detailed tissue culture experiments are needed, which are beyond this study’s scope. The major findings were based on data from P7 (transcriptomics) and P56 (structure and function). Lastly, small RNA-seq currently cannot be performed at single-cell resolution, so this data reflects a heterogeneous cell population with an unknown cellular composition.

## Materials and Methods

### Animals, Cre induction, EdU labeling

All mice (species *mus musculus*) used in this study were housed at the State University of New York, College of Optometry, following the Institutional Animal Care and Use Committee-approved protocols (IACUC). The *Ascl1-creERT2* strain (ID 12882, generated by Dr. Jane Johnson, (Kim et al., 2011), the *R26-stop-flox-CAG-tdTomato* strain (also known as Ai14, ID 007908), as well as the Dicer conditional knockout strain (*Dicer^f/f,^* ID 006001 (Harfe et al., 2005), were obtained from the Jackson Laboratory to create the *Ascl1CreER: stop^f/f^-tdTomato* strain further referred to as wildtype (wt) and the *Ascl1CreER: Dicer^f/f^*: *stop^f/f^-tdTomato* further referred to as Ascl-Dicer-cKO strain (cKO). The homozygous Dicer-cKO_RPC_ mouse will have a functionless Dicer enzyme in late RPCs due to the excision of exon 23 of the *Dicer1* gene via Cre recombinase (Harfe et al., 2005). Heterozygous mice (Dicer^fl/wt)^were not used in this study. Males and females were used, with experimental groups consisting typically of more than one litter. Cre negative littermates, as well as S129 background wildtype mice, were used as reference controls. *Cre*-positive not tamoxifen-treated mice were checked for reporter induction. Genotyping was done using the primers listed in Table 1. Successful deletion of exon 23 of *Dicer1* was confirmed via PCR for retinal lysates at P4, P7, and adult mice. Tamoxifen (Sigma, St. Louis, MO) was administered intraperitoneally at 75 mg/kg in corn oil for three consecutive days to initiate the recombination of the floxed alleles for inducing tdTomato at postnatal day (P) 1-3, P4-6, P9-11, or P13-15, or to initiate Dicer deletion at P1-3. Labeling of dividing cells using 5-Ethynyl-2’-deoxyuridine (EdU) was conducted at P3. EdU (Santa Cruz, cat. sc-284628) was reconstituted in sterile PBS and 25 μl injected subcutaneously into the back (skinfold) of the P3 old pups (10 mg/mL)(Kaufman et al., 2021). Animals were analyzed at P4, P7, P14, P28, P56, or at 3- and 4 months of age.

**Table 1:**
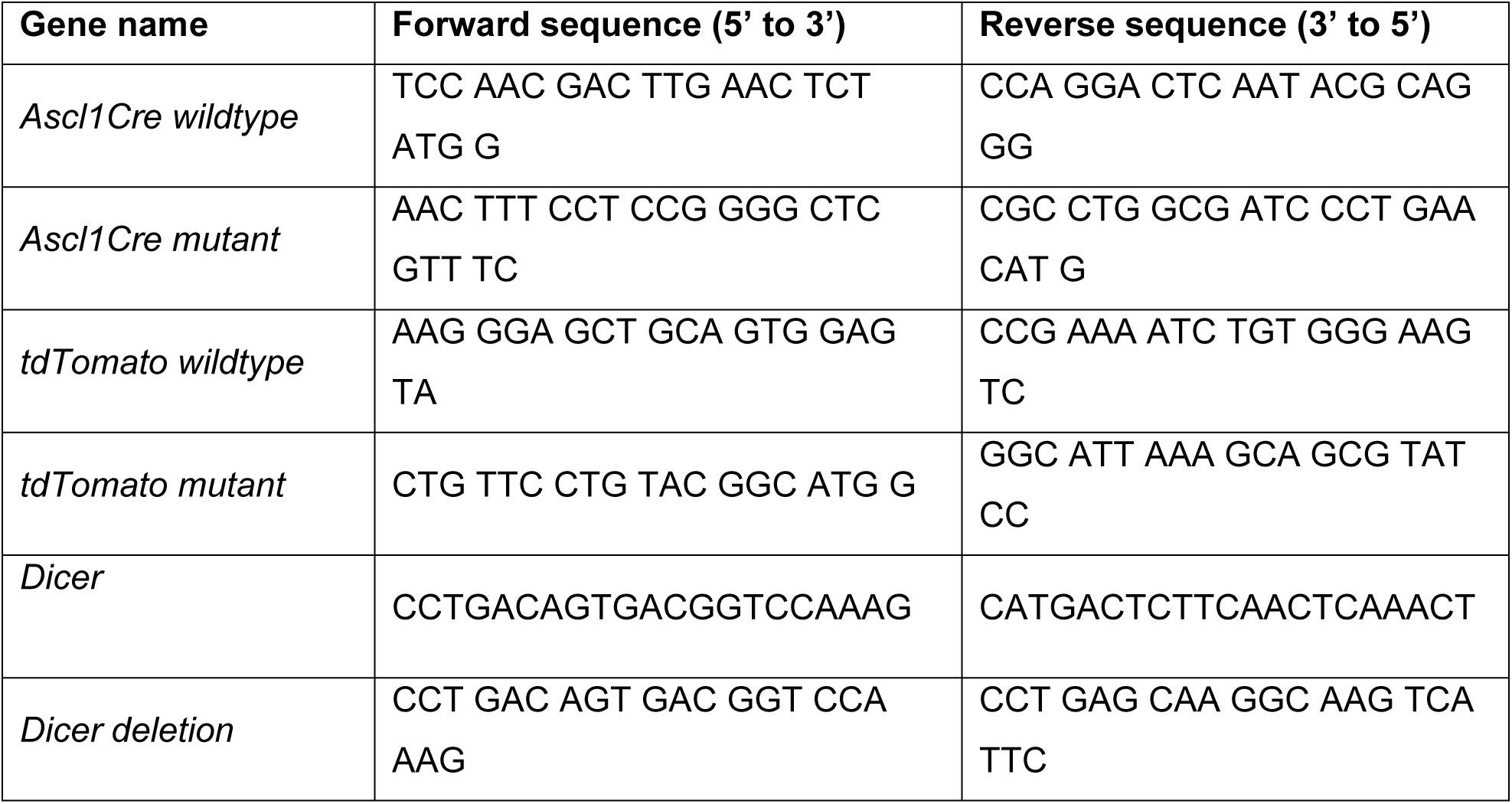
Genotyping Primers.

### Retinal spectral-domain optical coherence tomography (SD-OCT) imaging

*In vivo*, retinal structure and integrity were assessed using spectral-domain optical coherence tomography (SD-OCT) imaging using the Envisu R2200 SD-OCT device (Bioptigen, Durham, NC). Animals were anesthetized using 75-100 mg/kg Ketamine and 5-10 mg/kg Xylazine dissolved in sterile saline, injected intraperitoneally. Pupils were dilated with phenylephrine hydrochloride (2.5%) and tropicamide (0.5%). Methylcellulose was used to keep the corneas lubricated during the scans. Rectangular and radial volume scans were obtained while centered on the optic nerve head (1.4 x 1.4 mm, 1000 A-scans/B-scan X 15 frames/B-scan). For image analysis and measurements, retinal layers were manually segmented in a 9 x 9 plot in four regional quadrants (superior-inferior, nasal-temporal). Thicknesses were measured using DiverRelease_2_4 software. The total retinal thickness was defined as the distance between the inner border of the retinal nerve fiber layer (RNFL or NFL) and the outer border of the retinal pigment epithelium (RPE). All nuclear and plexiform layers were analyzed, including the nerve fiber layer /ganglion cell layer (NFL/GCL), inner plexiform layer (IPL), inner nuclear layer (INL), outer plexiform layer and outer nuclear layer (ONL). The inner and outer segments as well as the RPE were measured individually. The distance between the outer border of the outer limiting membrane (OLM, also known as external limiting membrane, ELM) to the outer border of the RPE were examined as whole allows for more accurate measurement of overall outer retinal thickness, which can be a valuable biomarker for various retinal conditions (Karampelas et al., 2014, Pandya et al., 2024). Thickness measurements were conducted in the central areas, within a radius of approximately 650 μm from the optic nerve (ON).

### Electroretinogram (ERG) recordings

For electroretinography, mice were dark-adapted overnight and anesthetized the following day via an intraperitoneal injection of 75-100 mg/kg Ketamine and 5-10 mg/kg Xylazine dissolved in sterile saline. Recordings were performed using the Espion Electrodiagnostic rodent system (Diagnosys LLC, Lowell, MA, USA) in a darkroom. Pupils were dilated with 1% tropicamide ophthalmic solution, and the eyes were lubricated using 1% methylcellulose. During the recording, mice were kept on a heat pad to maintain a constant body temperature. A gold wire electrode served as the recording electrode, while needles inserted into the cheek and back of the mouse served as reference and ground electrodes, respectively. A handheld, portable stimulator was used as the light source. Full-field 5-ms flashes were used to elicit responses. Dark-adapted ERGs were obtained with light intensities ranging from 0.001 to 64 cd.s/m^2^. To assess photoreceptor functionality, scotopic and photopic ERG a- and b-wave amplitudes were measured from the baseline to the a-wave trough and from the a-wave trough to the b-wave peak, respectively. For an in-depth analysis of the scotopic b-wave, the Naka-Rushton equation was employed, where V(I) = Vmax (In) / (In + Kn), as shown in Equation 1. The intensity–response amplitudes were fitted using the generalized equation: “V” is the amplitude at stimulus intensity “I”. “Vmax” is the maximum/saturated amplitude and an indirect measurement for responsiveness. “K” is the semi-saturation constant representing the stimulus intensity I at which half of the saturated amplitude Vmax is reached. Hence, K is an indirect measurement of sensitivity. “n” is the slope of the fit curve and an indirect measurement of heterogeneity (Severns and Johnson, 1993, Evans et al., 1993).

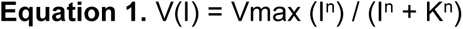

Generalized Naka-Rushton equation for estimating fit parameters.

To assess the function of the inner retinal layers, oscillatory potentials (OPs) and positive scotopic threshold responses were analyzed. OPs are low amplitude wavelets superimposed on the ascending b-wave and are believed to be generated by the inhibitory feedback interactions between amacrine, bipolar and ganglion cells (Cobb and Morton, 1952, Liao et al., 2023, Wachtmeister, 1998, Wachtmeister and Dowling, 1978). Total OP amplitude was determined by adding the amplitudes of each OP wavelet (OP1, OP2, OP3, and OP4). A single OP wavelet amplitude was calculated from the trough to the peak of that wavelet, as previously described (Akula et al., 2007). Scotopic threshold responses (STRs) were generated in the proximal retina and are believed to reflect the activity of retinal ganglion and amacrine cells (Naarendorp and Sieving, 1991, Sieving et al., 1986). Positive scotopic threshold responses are waveforms with positive polarity similar to b-wave but are elicited at light levels too low to generate the b-wave produced by bipolar cells (Saszik and Bilotta, 2001). The pSTR amplitude was evaluated from the pre-stimulus baseline to the peak of the positive deflection.

### Tissue preparation for immunofluorescent labeling

Mice were euthanized, and eyes were marked at the nasal side, enucleated, and transferred to pre-chilled phosphate-buffered saline (wash, PBS, Fisher Scientific, Hampton, NH, USA). <u>For retinal cross sections</u>, the eyeball was fixed in cold 4% paraformaldehyde (PFA, VWR International, Radnor, PA, USA, 4°C) for 20 min. The cornea, lens, iris, and vitreous body were removed (video available in (Kang and Wohl, 2022), and eyecups were fixed for an additional 20 min in cold 4% PFA. After fixation, eyecups were subsequently washed for 10 min in pre-chilled PBS, and incubated with 30% sucrose (Fisher Scientific, Hampton, NH, USA) in PBS overnight at 4°C. The tissue was embedded in O.C.T. embedding medium (Sakura Finetek, Torrance, CA, USA) with nasal-temporal orientation and frozen at -80°C. The frozen tissue was cross-sectioned into 12 μm-thick sections for subsequent immunofluorescent labeling. For labeling, frozen sections were dehydrated at 37°C for 20 min, then fixed for 20 min with 4% PFA, and subsequently washed in PBS. Sections were incubated in blocking solution (10% horse serum, Fisher Scientific, Hampton, NH, USA, in PBS; with 0.5% Triton-X100, Sigma-Aldrich, St. Louis, MO, USA) for at least 1 hour at RT, then incubated with primary antibodies (Table 2) overnight. Secondary antibody incubation was done on the following day for 1 hour at RT. All antibodies were diluted in 10% horse serum. 4,6-Diamidino-2-phenylindole (DAPI, Sigma-Aldrich, St. Louis, MO, USA, 1:2,000) was used for counterstaining. For EdU labeling, the Click-iT™ EdU Alexa Fluor™ 647 Imaging Kit (Catalog #C10340, Fisher Scientific) was used, following the manufacturer’s instructions. Slides were mounted with cover glasses and mounting medium (Invitrogen, Waltham, MA, USA). Only validated antibodies were used in this study. Validation was based on item information given by the vendor as well as publications.

**Table 2:** Primary and Secondary antibodies.

| Antibody | Concentration | Company, Catalog # | Reference |
| --- | --- | --- | --- |

| **Primary antibodies/dyes** |  |  |  |
| --- | --- | --- | --- |
| mouse anti Brn3a | 1:500 | Millipore, MAB1585 | (Schneider et al., 2001) |
| mouse anti Calbindin | 1:500 | Millipore, ABN 2192 | (Wassle et al., 1998) |
| goat anti ChAT | 1:250 | Millipore, AB144P | (West Greenlee et al., 1998) |
| mouse anti glutamine synthetase (GS) | 1:500 | Millipore, MAB302 | (Grossman et al., 1994) |
| rabbit anti HuC/D | 1:250 | Fisher, A-21271 | (Marusich et al., 1994) |
| rabbit anti Ki67 | 1:250 | Abcam, ab15580 | (Gerdes et al., 1984) |
| rabbit anti M-Opsin | 1:300 | Sigma-Millipore, AB5407 | (Applebury et al., 2000) |
| goat anti Otx2 | 1:250 – 1:500 | R&D Systems, AF1979 | (Martinez-Morales et al., 2004) |
| rabbit anti Pax6 | 1:250 | Invitrogen, 42-6600 | (Marquardt et al., 2001) |
| Peanut agglutinin (PNA) fluorescein (a lectin) | 1:300 | Vector labs, FL-1071 | (Blanks and Johnson, 1983) |
| rat anti Phosphohistone3 (PH3) | 1:250 | Novus, 102133-030 | (Dhomen et al., 2006) |
| mouse anti PKC | 1:500 | Millipore, P5704 | (Greferath et al., 1990) |
| guinea pig anti Prox1 | 1:250 | SYSY antibodies, 509 005 | (Dyer et al., 2003) |
| rat anti RFP (tdTomato) | 1:1000 | Antibodies online, ABIN334653 | validation based on previous work (Wohl et al., 2019, Wohl et al., 2017) |
| rabbit anti Ribeye | 1:500 | SYSY antibodies, 192 103 | (Schmitz et al., 2000) |
| goat anti Sox2 | 1:500 | R&D Systems, AF2018 | (Catena et al., 2004) |
| rabbit anti Sox9 | 1:250 | Millipore, AB5535 | (Poche et al., 2008) |
| sheep anti Vsx2 | 1:250 | Fisher, 50-210-1941 | (Burmeister et al., 1996) |
| rabbit anti ZO-1 | 1:300 | Invitrogen, 61-730-0 | (Konari et al., 1995) |
| **Secondary antibodies** |  |  |  |
| Alexa Fluor 488 -<br>AffiniPure F(ab')<sub>2</sub><br>Fragment Donkey Anti-Goat IgG (H+L) | 1:500 | Jackson<br>ImmunoResearch<br>Laboratories, Inc<br>705-546-147 | validation based on<br>previous work (Wohl et<br>al., 2019, Wohl et al.,<br>2017) |
| Alexa Fluor 647 -<br>AffiniPure F(ab')<sub>2</sub><br>Fragment Donkey Anti-Goat IgG (H+L) | 1:500 | Jackson<br>ImmunoResearch<br>Laboratories, Inc.<br>705-606-147 | validation based on<br>previous work (Wohl et<br>al., 2019, Wohl et al.,<br>2017) |
| Alexa Fluor 488-AffiniPure F(ab')<sub>2</sub><br>Fragment Donkey Anti-Mouse IgG (H+L) | 1:500 | Jackson<br>ImmunoResearch<br>Laboratories, Inc.<br>715-546-150 | validation based on<br>previous work (Wohl et<br>al., 2019, Wohl et al.,<br>2017) |
| Alexa Fluor 488 -<br>AffiniPure F(ab')<sub>2</sub><br>Fragment Donkey Anti-Rabbit IgG (H+L) | 1:500 | Jackson<br>ImmunoResearch<br>Laboratories, Inc.<br>711-546-152 | validation based on<br>previous work (Wohl et<br>al., 2019, Wohl et al.,<br>2017) |
| Alexa Fluor 647 -<br>AffiniPure F(ab')<sub>2</sub><br>Fragment Donkey Anti-Rabbit IgG (H+L) | 1:500 | Jackson<br>ImmunoResearch<br>Laboratories, Inc.<br>711-606-152 | validation based on<br>previous work (Wohl et<br>al., 2019, Wohl et al.,<br>2017) |
| Rhodamine Red 570 -<br>AffiniPure F(ab')<sub>2</sub><br>Fragment Donkey Anti-Rat IgG (H+L) | 1:1000 | Jackson<br>ImmunoResearch<br>Laboratories, Inc.<br>712-296-150 | validation based on<br>previous work (Wohl et<br>al., 2019, Wohl et al.,<br>2017) |

<u>For retinal flatmounts and RPE preparation</u>, muscles and connective tissues on the sclera were removed from the eyeball to facilitate the tissue flattening. The cornea, iris, lens, and vitreous body were removed, and the RPE (with attached retina) was carefully transferred onto filter paper. The tissue was cut radially (4 times), and the retina gently detached and fixed with cold 4% PFA for at least 30 min up to 1 hour at RT. The RPE (with sclera) was fixed with cold 4% PFA for 1-2 hours at RT. After fixation, the tissue was washed three times in PBS at RT for 10 min and stained following the protocol as described above and the markers listed in Table 2.

### Microscopy and cell count

Retinal eye cups were imaged using a confocal laser scanning microscope (FM1200 and FV3000, Olympus, Shinjuku City, Tokyo, Japan) and the Fluoview FV31S software for whole tissue imaging. Whole cup stitched images were taken with 10 x and 20 x objectives. Images of peripheral areas (close to the ora serrata) and central areas (within 500 μm distance from the optic nerve [ON]) were taken with 20 x or 40 x objectives as z-stacks. For confocal analysis, image parameters were defined by control/wildtype groups and used (reloaded) for all conditions. Images were processed and analyzed using Adobe Bridge, Adobe Photoshop, and Affinity. Cells per field (field = full image) or photoreceptor segments per field (field = full image) were counted for at least 4 images per mouse per staining protocol from central and peripheral cross-sections. The number of rod columns was counted for 10 areas per image for at least 4 images of central areas per mouse. For P56 analysis, at least 6 mice per group were evaluated per group and condition (power analysis 85%); the exact n is given in the legends. For color-blind friendly figures, either gray scale or green/magenta images were chosen or generated using Photoshop.

### Fluorescence-activated cell sorting

P7 wildtype and Dicer-cKO retinas were checked for successful recombination and reporter induction using a fluorescence microscope (Keyence BZ-X810, KEYENCE, Elmwood Park, NJ, USA). For each sort, the retinas of 1-3 reporter+ mice were pooled and dissociated in DNase/Papain (50 µl/ 500 µl respectively, Worthington Biomedical, Lakewood, NJ, USA) for 20 min at 37⁰C on the shaker (Thermomixer, Fisher Scientific, Hampton, NH, USA), triturated, mixed with Ovomucoid (500 µl, Worthington Biomedical, Lakewood, NJ, USA), and centrifuged for 10 min at 300 x g. The pellet was resuspended in 800 µl DNase/ Ovomucoid/ Neurobasal without phenol red medium (Gibco, Hampton, NH, USA) solution per retina (1: 1: 10 respectively). Cells were filtered through a 35-µm filter, sorted using a 100-micron nozzle (BD Melody, BD Bioscience, Franklin Lakes, NJ, USA), and collected into two chilled FBS-coated tubes containing Neurobasal medium without phenol red. Cells with the brightest fluorescence were considered “positives” (RPC-progeny fraction), and cells with no fluorescence (“negatives”) were considered as the non-progeny fraction. Debris was excluded. After collection, the tdTomato^+^ fraction and the tdTomato^−^ fraction were post-sorted, and a fraction of each sort was plated in a 48-well plate to confirm and validate purity and viability using 7-AAD viability staining (BD Bioscience). After the sort, cells were spun for 10 minutes at 300 × g at 4 °C. Pellet was either resuspended in HBSS for scRNA-seq and cell numbers were ascertained using an automated cell counter (Countess, Thermo Fisher), or homogenized in RLT lysis buffer (miRNeasy Tissue/Cells Advanced Micro Kit, 217684, Qiagen, Hilden, Germany) for subsequent RNA extraction.

### Single-cell RNA Sequencing and Data Analysis

Single-cell RNA sequencing was performed using pre-templated instant partitions (PIP) sequencing (PIPseq, (Lee et al., 2024)). The T20 kit (≤ 20,000 cells output, FBS-SCR-T20-4-V4.05, Fluent Biosciences) was used according to the manufacturer’s instructions. In brief, tubes containing template beads were thawed on ice, and the required number of input cells was added per tube (i.e., approximately 5,000-5,500 live cells for the T2 or 25,000–30,000 live cells for the T20 kit). The suspension was mixed gently by pipetting and processed further according to the manufacturer’s instructions. After the cDNA synthesis, purification, and library preparation, the concentration, integrity, and quality were assessed using the Agilent 2100 Bioanalyzer System and the high-sensitivity DNA kit (Agilent Technologies, Santa Clara, CA, USA). Illumina-compatible libraries were sequenced using the NovaSeq X+ Sequencing System (Illumina, 40,000 reads per cell) at the NYU Genome Technology Center. FASTQ files were preprocessed using PIP Seeker, a comprehensive analysis solution that performs barcode matching, quality control, mapping with the STAR aligner40, deduplication, transcript counting, cell calling, clustering, differential expression, and cell type annotation. The resulting matrices from PIPseeker were imported into Monocle3 (Cao et al., 2019, Qiu et al., 2017, Trapnell et al., 2014) for downstream analysis. Matrices were aggregated and first-pass filtering for cell quality was performed, keeping cells with < 25% reads coming from mitochondrial transcripts and displaying total transcript numbers between 500 and 10,000 transcripts per cell. Dimension reduction was performed using the top 3234 transcripts displaying high variance across the data using UMAP dimension reduction (McInnes et al., 2020) on the top 17 PCAs (umap.min_dist = 0.1, umap.n.neighbors = 15,umap.metric = “euclidean”). Cells were clustered within the reduced dimension space using the cluster_cells function (reduction_method = “UMAP”, cluster_method = “louvain”, num_iter = 5, partition_qval = 0.01). Cell type calls were performed based on enrichment of transcripts within individual clusters using established markers (Clark et al., 2019). Similar analyses were utilized to determine major amacrine cell subtypes after sub-setting of amacrine cells from the total datasets. Fastq are deposited at Gene Expression Omnibus (GEO, https://www.ncbi.nlm.nih.gov/geo), accession number GSE288231.

### RNA extraction and RNA sequencing

Total RNA was isolated using the RNeasy Plus Mini Kit (Qiagen, Hilden, Germany) per the manufacturer’s instructions. RNA quantity and quality were assessed via NanoDrop 2000 (Thermo Scientific, Waltham, MA, USA) as well as the RNA 6000 Pico Kit (Agilent Technologies, Santa Clara, CA, USA) for 2100 Bioanalyzer Systems. All RNA samples were submitted to the NYU Langone Health Genome Technology Center for processing.

The SMART-Seq HT Kit (Takara) was utilized to generate complementary DNA (cDNA) from 1 ng of total RNA input. The synthesis process included 13 PCR cycles. The quality and concentration of the stock RNA were assessed using an Agilent Pico Chip on the Bioanalyzer system, ensuring high-quality RNA inputs. For library preparation, cDNA was quantified using the Invitrogen Quant-iT system and subsequently diluted to a concentration of 0.3 ng/μL for library preparation. Libraries were prepared using the Nextera XT DNA Library Preparation Kit, employing a total of 12 PCR cycles to amplify and uniquely barcode the samples. The quality of the generated libraries was assessed using the Agilent High Sensitivity TapeStation to verify fragment sizes and integrity. Additionally, the concentration of libraries was measured with the Invitrogen Quant-iT system to ensure accurate normalization prior to pooling. For sequencing, libraries were pooled in equimolar ratios and sequenced on the Illumina NovaSeq X+ platform using paired-end 100-cycle reads.

Small RNA libraries were prepared from 10 ng of purified miRNA using the SMARTer® smRNA-Seq Kit for Illumina (Takara Bio), following the manufacturer’s protocol. The kit utilizes a ligation-free method based on SMART (Switching Mechanism at the 5′ end of RNA Template) technology to generate high-quality libraries from low-input small RNA. After adapter ligation and cDNA synthesis, libraries were amplified using PCR and purified using AMPure XP beads according to the protocol specifications. The quality and size distribution of the final libraries were assessed using the Agilent High Sensitivity D1000 ScreenTape on the TapeStation 4200 system (Agilent Technologies) to confirm the presence of expected small RNA insert sizes (∼140–150 bp for miRNA). Library concentrations were measured using the Quant-iT™ dsDNA High Sensitivity Assay Kit (Invitrogen) to ensure accurate quantification. Libraries were normalized to equimolar concentrations based on fluorometric measurements and pooled accordingly. Sequencing was performed on the Illumina NovaSeq X+ platform using paired-end 100-cycle reads, providing high-depth coverage suitable for small RNA transcriptome analysis.

### Bulk RNA-seq data processing and analysis

All sequencing data were processed by the Applied Bioinformatics Laboratories (ABL) at New York University School of Medicine using standard bioinformatics pipelines. Cutadapt (v5.0) (Martin, 2011) was used to cut the adaptors from the small RNA-seq data. Sequencing reads from both small and large RNA-Seq projects were mapped to the reference genome (mm10) using the STAR aligner (v2.5.0c) (Dobin et al., 2013). Alignments were guided by a Gene Transfer Format (GTF) file. The mean read insert sizes and their standard deviations were calculated using Picard tools (v.1.126) (http://broadinstitute.github.io/picard). The read count tables were generated using HTSeq (v0.6.0) (Anders et al., 2015), normalized based on their library size factors using DEseq2 (Love et al., 2014), and differential expression analysis was performed. The Read Per Million (RPM) normalized BigWig files were generated using BEDTools (v2.17.0) (Quinlan and Hall, 2010) and bedGraphToBigWig tool (v4). Gene set enrichment analysis was performed using GSEA tool (PMID: 16199517). To compare the level of similarity among the samples and their replicates, we used two methods: principal-component analysis and Euclidean distance-based sample clustering. All the downstream statistical analyses and generating heatmaps and plots were performed in R environment (v3.1.1) (https://www.r-project.org/). Because of the possibility of a global effect in mature miRNA levels in our Dicer1 knockout (cKO) model, we made count tables for all genes and used the none miRNA genes as our spike-in factor. Spike-in normalization is generally used when gene expression is affected globally. We then used Geometric Library Size Factor (GLSF) normalization as described in DESeq R package for our spike-in factors and applied it to normalize the miRNA genes. Raw files (fastq) and RPM-normalized BigWig files were used to visualize genomic data in interactive genomic browsers (IGV). Fastq files and Bigwig files were deposited at Gene Expression Omnibus (GEO, https://www.ncbi.nlm.nih.gov/geo), accession number GSE288231.

### miRNA-target interaction analysis and prediction tools to identify RPC miRNA targets

Different prediction tools and genome browsers were used to find putative targets (correct and latest transcript) of late RPC miRNAs (Nanostring data set from (Wohl et al., 2019), available at GEO (GSE135835, under SuperSeries accession number GSE135846), including miRWalk, STarMir, and microT (Diana tools) (Tastsoglou et al., 2023). miRWalk (http://mirwalk.umm.uni-heidelberg.de) is a database of predicted, as well as experimentally validated, miRNA-target interaction pairs within the complete sequence of genes resulting from the TarPmiR algorithm. TarPmiR is a random-forest-based approach that was trained with 13 features, including Seed match, Accessibility, AU content, exon preference, and the length of the target mRNA region. It also integrates results from other databases with predicted and validated miRNA-target interactions (Sticht et al., 2018); STarMir web server (http://sfold.wadsworth.org/starmir.html) is an application module of the Sfold RNA package (http://sfold.wadsworth.org), which predicts miRNA binding sites on a target mRNA (Rennie et al., 2014). STarMir is a logistic prediction model developed with miRNA binding data from CLIP studies which predicts miRNA binding sites and computes comprehensive sequence, thermodynamic and target structure features for any given gene. The sequence information for several miRNAs (miRNA IDs) and our single target mRNA (RefSeq ID) were entered manually and the HITS-CLIP mouse model for prediction was selected. The miRNA-mRNA binding sites results predicted from the STarMir web server range from one up to around 40 hybrid conformations, which were cross-referenced with those from miRWalk. In contrast to TargetScan, these tools utilize the most current and up-to-date mRNA transcript (https://mart.ensembl.org/info/genome/ genebuild/canonical.html), resulting in updated matches of miRNA:mRNA interactions. However, TargetScan was also used as a reference.

### Luciferase Assay

The 3’ UTR of *Elavl3* (obtained from UCSC Genome Browser, https://genome.ucsc.edu/), was cloned into the IV-80 pmiRGLO vector (gift from Dr. Volker Busskamp, University of Bonn, Addgene plasmid #78128, (Eichelser et al., 2014). The restriction enzymes NheI and XbaI were used with specific primers for the 3’ UTR region to generate the construct (Table 3) as previously described (Kutsche et al., 2018). The empty IV-80 vector (without a 3’UTR) was a negative control for the assay. These constructs were transfected with mimics (Dharmacon Horizon, Lafayette, CO, USA) for the RPC miRNA miR-25-3p, miR-20a-5p, miR-15a-5p, and the proneuronal miRNA miR-124-3p as well as a control mimic (C.elegans, Table 4). Four independent experiments were conducted with three technical replicates using HEK293T cells (gift from Dr. Ching-Hwa Sung, Cornell University) and Lipofectamine 3000 (L3000015, Invitrogen, Waltham, MA, USA). 48 h after transfection, Dual-Glo Luciferase reagents were added to the cells (E2920, Promega, Madison, WI, USA). Luciferase activity (Firefly and Renilla) was measured using a plate reader (FilterMax F3, Molecular Devices, San Jose, CA, USA). Data was normalized against negative controls (Plasmid only), and RPC-miRNA treatments were compared to control mimic transfected conditions.

**Table 3:**
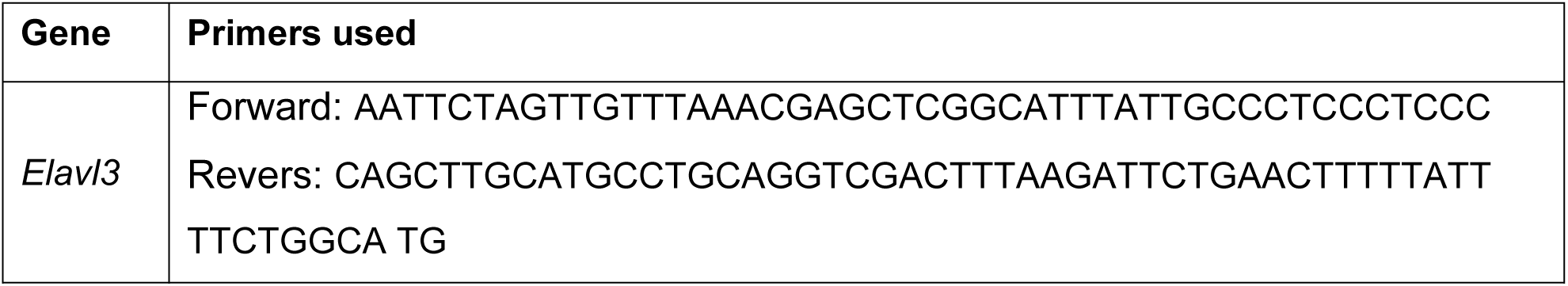
Luciferase assay primers for 3‘ UTR amplification.

**Table 4:** miRNA Mimics.

| Name | Catalog number | microRNA ID |
| --- | --- | --- |
| *Negative control mimic* | CN-001000-01-50 | **MIMAT0000039** |
| *mmu-miR-25-3p mimic* | C-310564-05-0020 | **MIMAT0000652** |
| *mmu-miR-20a-5p mimic* | C-310514-05-0002 | **MIMAT0000529** |
| *mmu-miR-15a-5p mimic* | C-310510-05-0020 | **MIMAT0000526** |
| *mmu-miR-124-3p mimic* | C-310390-05-0020 | **MIMAT0000134** |

### Statistical Analysis

Statistical analyses were performed using SPSS or Sigma Plot by using the Mann-Whitney U-Test, a non-parametric test for independent samples. The Mann–Whitney U test is preferable to the t-test when the data are ordinal but not interval scaled, hence the spacing between adjacent values of the scale cannot be assumed to be constant. Holm-Bonferroni method was used to correct for multiple comparisons. Based on power analysis (85% power, alpha 0.5) and previous studies, an n of at least 5 mice per group were used. Only animals that could not be analyzed properly, i.e. methodological complications (for instance, anesthesia complications, noisy ERG recordings, missing or insufficient reporter labeling) were excluded, which was rarely the case. For randomization and to reduce badge effect, animals of several litters were analyzed. The genotype was withheld from the investigator evaluating cell numbers or expression levels at the time point of analysis to guarantee unbiased results.

## Acknowledgment

The authors would like to thank Dr. Adriana Heyguy and her outstanding team from the NYU Langone Health Genome Technology Center for handling our miRNA-seq and bulk RNA-seq samples, as well as for their exceptional support. We thank Dr. Aristotelis Tsirigos from the NYU Applied Bioinformatics Laboratories (ABL) for his support. We thank Autumn Cholger from FluentBio/Illumina for excellent training and in-depth troubleshooting. We thank Prof. Dr. Volker Busskamp and his team from the University of Bonn for providing protocols, reagents, and advice for the Luciferase assays. We thank Dr. Joseph A. Brzezinski IV from the University of Colorado for sharing the EdU injection protocol. We thank Dr. Ching-Hwa Sun from Weill Cornell Medicine for sharing HEK 293T cells. We also thank Dr. Miruna Ghinia-Tegla for suggesting the Ribeye antibody.

## Data and resource availability

All relevant data and resources can be found within the article and its supplementary information. scRNA-Seq as well as bulk RNA (long RNA) and miRNA (short RNA) data files were deposited at Gene Expression Omnibus (GEO), accession number GSE288231.

