## Supplementary Figures and Tables for "Dicer is essential for proper maturation, composition, and function in the postnatal retina"

Supplementary
Figures and
Tables

Kang et al., 2025

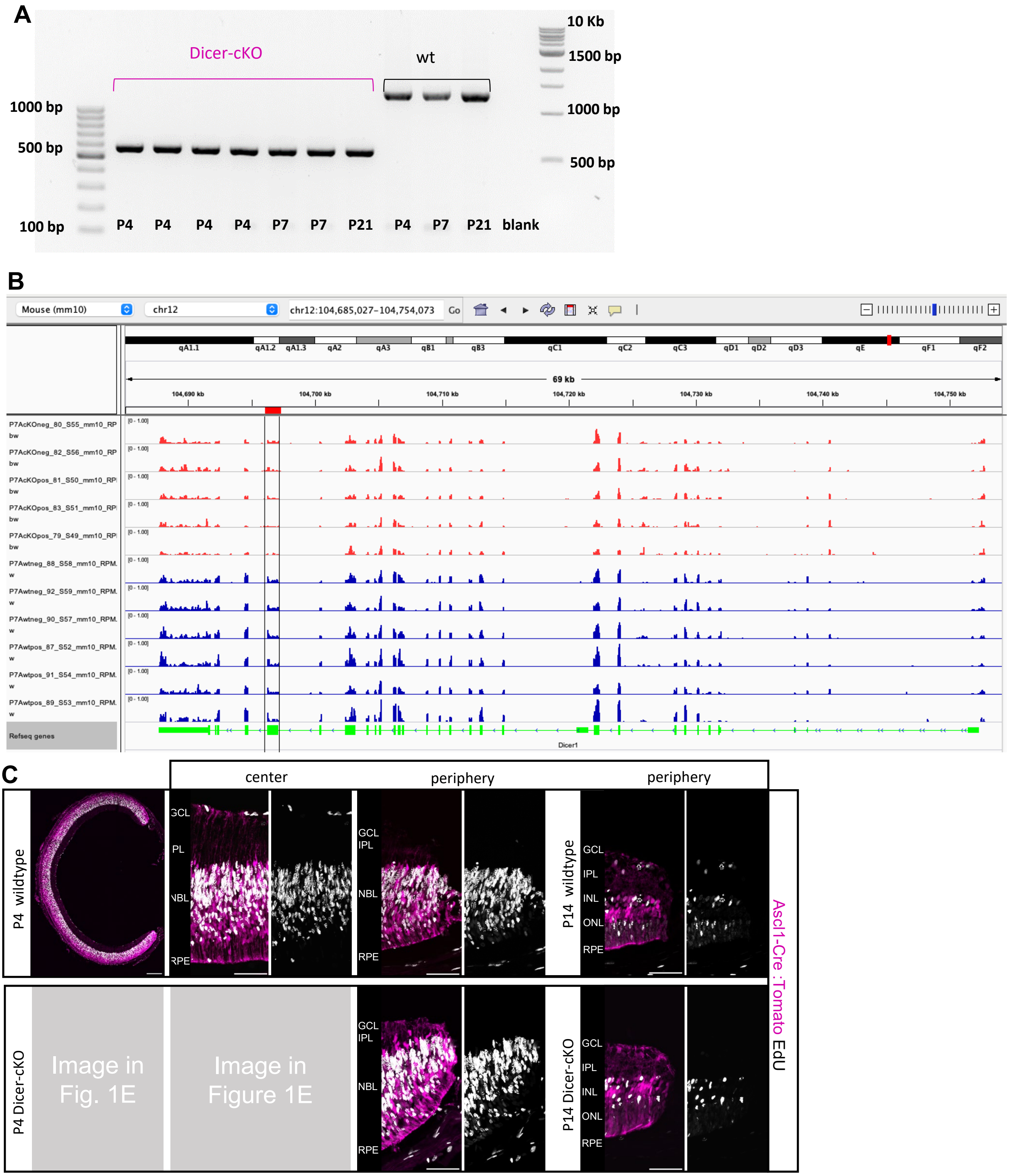


**Supplementary Figure 1. Dicer depletion in late proliferating retinal progenitor cells. A:** Gel image visualizing successful exon 23 deletion of *Dicer1* (~550 bp transcript) in P4, P7, and P21 retinal lysates of Ascl1-Cre:Dicer cKO mice (Dicer-cKO) mice as well as unaffected Dicer genes (1300 bp transcript) in P4, P7 and P21 wildtype (wt) retinal lysates. **B**: Screenshot of bulk RNA-seq data (BigWig files using Integrative Genomics Viewer IGV) showing exon 23 loss in P7 Dicer-cKO (AcKOpos) FACS-purified tomato+ cells as well as unaffected exons in wildtype (Awtpos) and reporter-negative cells (AcKOneg). **C:** P4 and P14 retinal cross sections with insets in higher magnification of central and peripheral areas, visualizing endogenous reporter expression of RPC progenies labeled at P1-3 as well as EdU-labeled cells at P3. Scale bars 200 µm, insets: 50 µm. GCL: ganglion cell layer, IPL: inner plexiform layer, INL: inner nuclear layer, OPL: outer plexiform layer, ONL: outer nuclear layer, NBL: neuroblastic layer, RPE: retinal pigment epithelium.

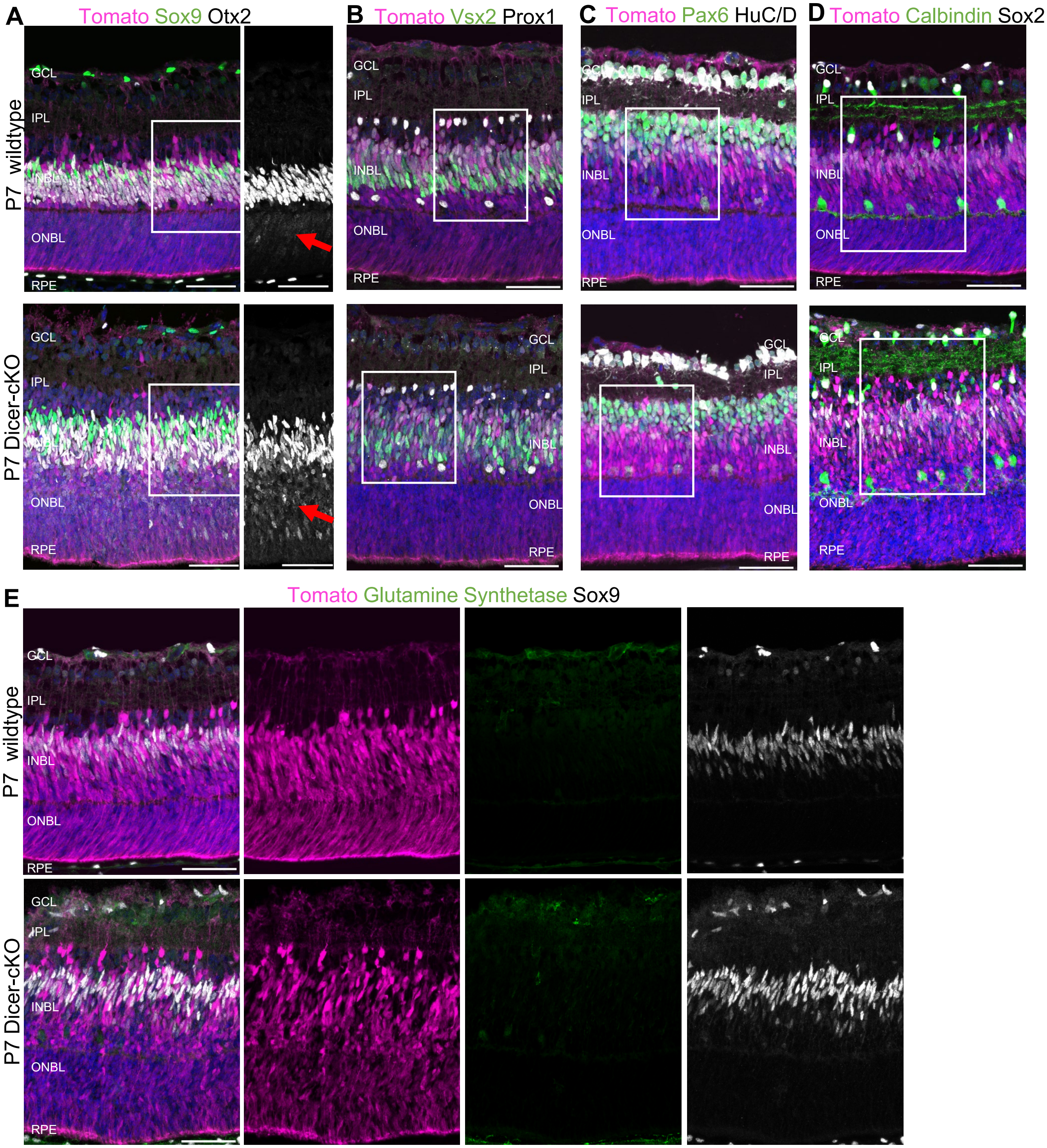


**Supplementary Figure 2. Dicer loss in late RPCs results in altered cellular structures of RPC progenies. A-E:** Immunofluorescent labeling with antibodies against Sox9, Otx2 (A), Vsx2, Prox1 (B), Pax6, HuC/D (C), Calbindin, Sox2 (D), and glutamine synthetase (GS), Sox9 (E) of P7 wildtype or Dicer-cKO central retinal cross sections to characterize Tomato+ RPC progenies in the INL. Insets in B-E are shown in the main Figure 2. Red arrows indicate Otx2+ cells in the ONBL. Scale bars 50 µm. GCL: ganglion cell layer, IPL: inner plexiform layer, INBL: inner neuroblastic layer, ONBL: outer neuroblastic layer, RPE: retinal pigment epithelium.

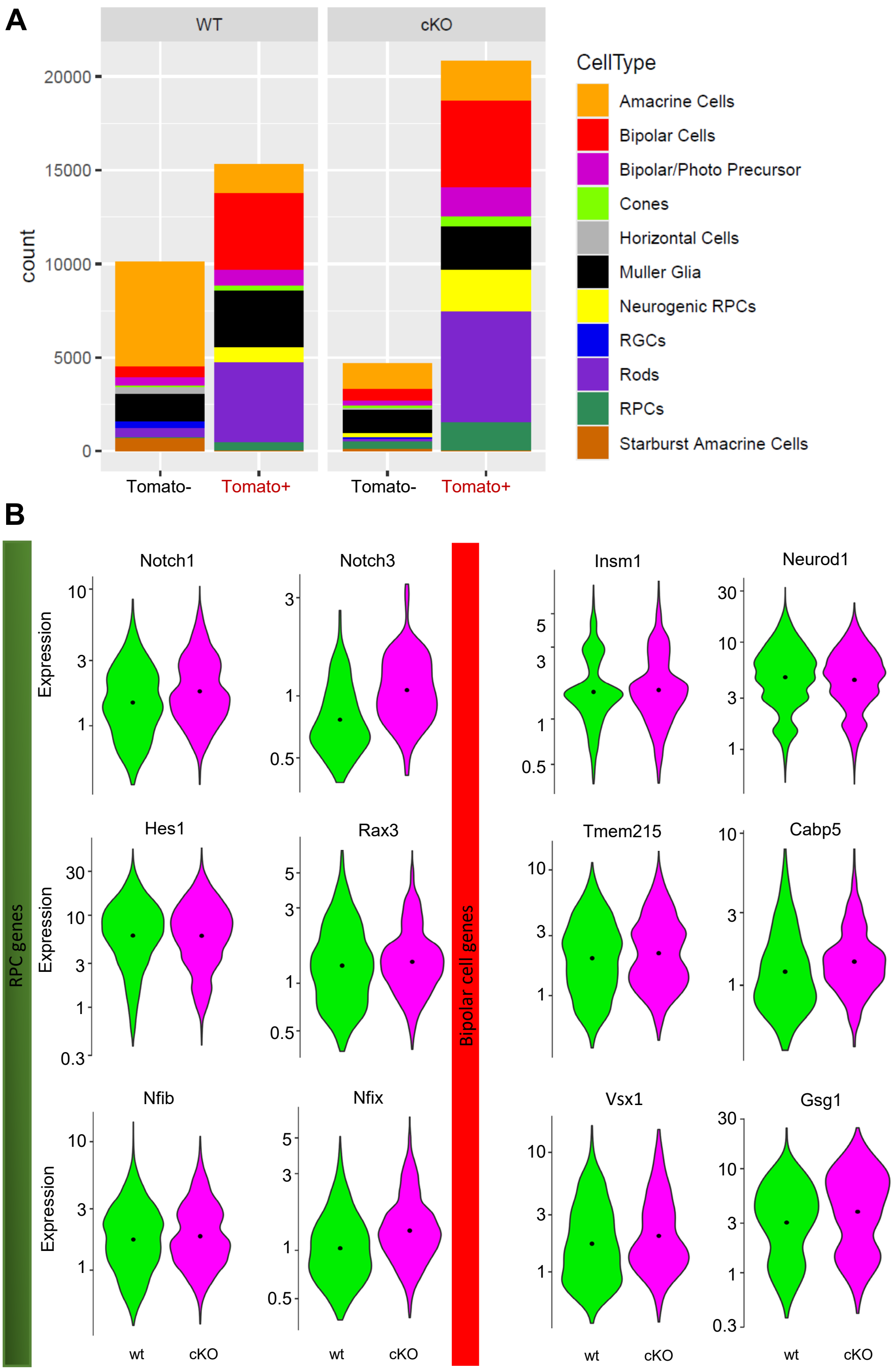


**Supplement Figure 3. scRNA-Seq of P7 progenies shows altered populations and gene expression. A:** Numbers (counts) of scRNA-seq captured cells of FACS-purified P7 progenitor populations of Ascl-Cre:tdTomato wildtype and Dicer-cKO mice, as well as corresponding Tomato-negative fractions, colored by annotated cell type as determined by marker gene expression. **B:** Violin plots of the cellular expression of marker genes of RPCs, Müller glia, and bipolar cells facetted by genotype. RPCs: retinal progenitor cells, RGCs: retinal ganglion cells.

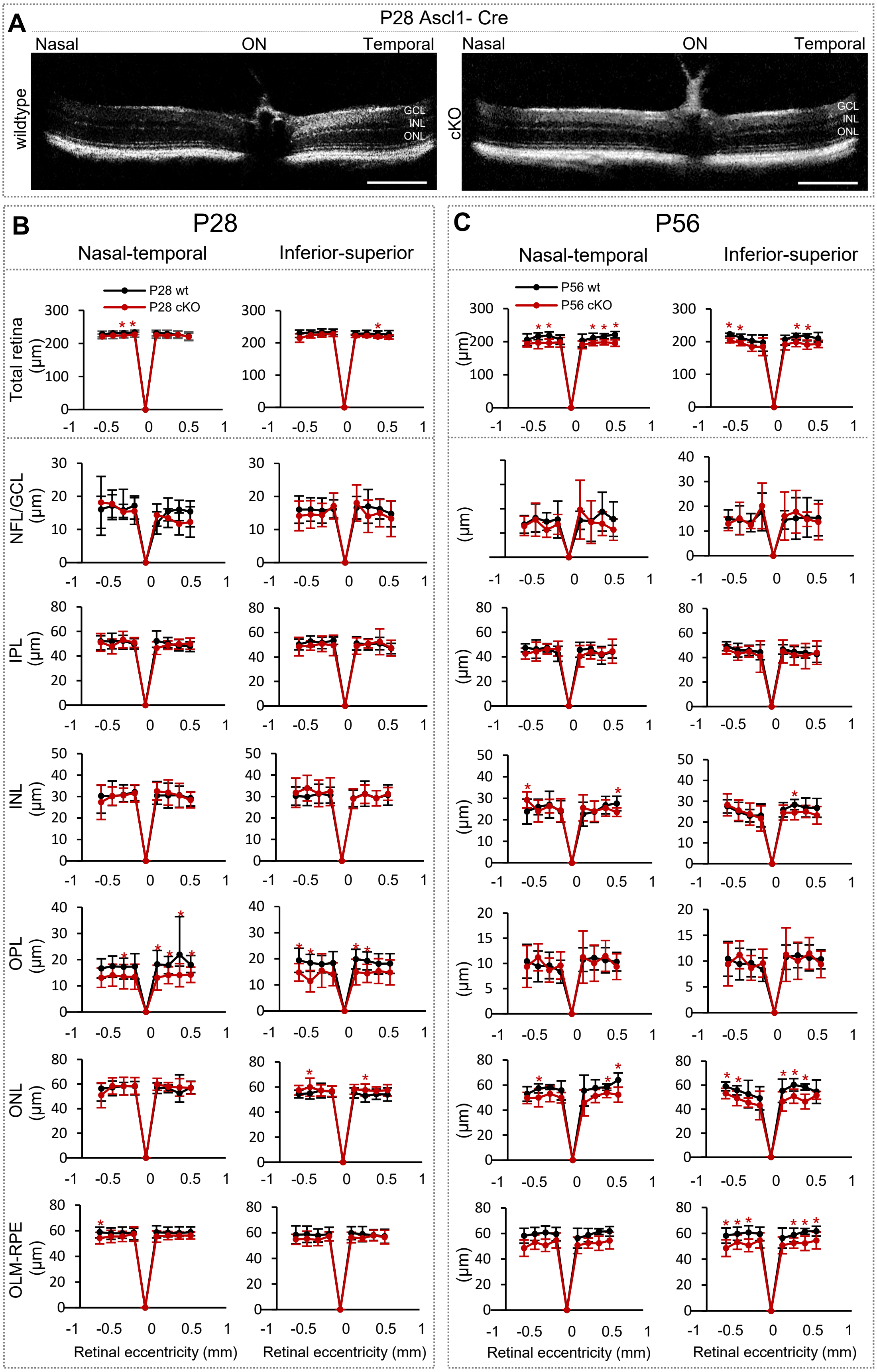


**Supplementary Figure 4: P28 and P56 Dicer-cKO retinas have normal appearance *in vivo* and minimal laminar alterations. A:** Spectral-domain optical coherence tomography (SD-OCT) images of postnatal day P28 wildtype (wt) and Ascl1-Cre:Dicer cKO$_{RPC}$ retinas (cKO). **B-C:** Spider plots of the thickness (µm) of the total retina and individual layers measured at the nasal-temporal or inferior-superior axis, of P28 (B) wt (n=7) and P28 cKO (n=9) mice and (C) p56 wt (n=7) and P56 cKO (n=6) mice. Mean ± S.D. Significant differences are indicated, Mann-Whitney-U-test: *: $p \leq 0.05$. NFL: nerve fiber layer, GCL: ganglion cell layer, IPL: inner plexiform, INL: inner nuclear layer, OPL: outer plexiform layer, ONL: outer nuclear layer, OLM: outer limiting membrane, RPE: retina pigment epithelium, ON: optic nerve.

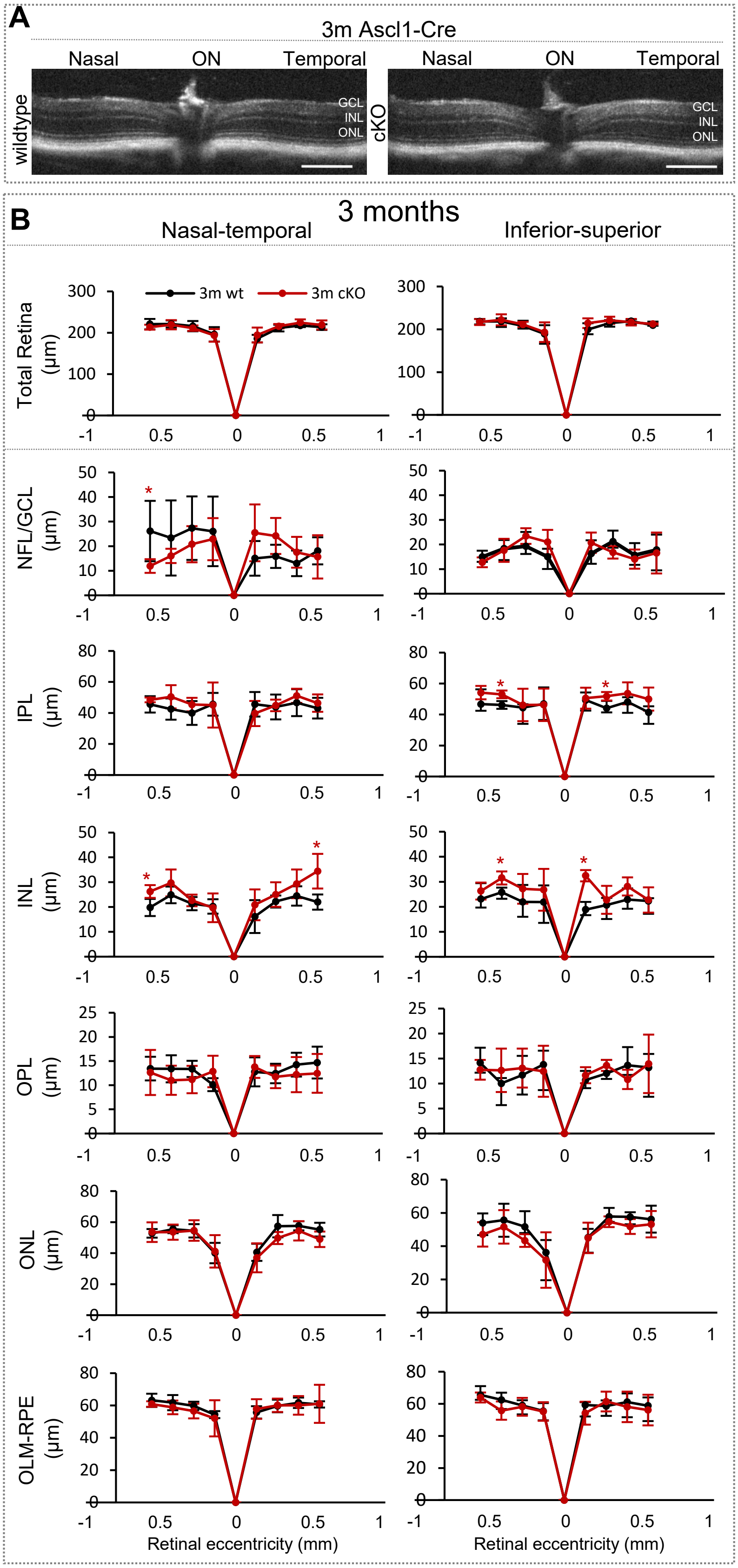


**Supplementary Figure 5: Three-month Dicer-cKO retinas show minimal laminar alterations. A:** Spectral-domain optical coherence tomography (SD-OCT) images of 3 month-old wildtype (wt) and Ascl1-Cre:Dicer $cKO_{RPC}$ mouse (cKO) retinas. **B:** Spider plots of the thickness (µm) of the total retina and individual layers measured at the nasal-temporal or superior-inferior axis, of 3-month old wt (n=5) and 3 month cKO (n=4) mice. mean ± S.D. Significant differences are indicated, Mann-Whitney-U-test: *: $p \leq 0.05$. NFL: nerve fiber layer, GCL: ganglion cell layer, IPL: inner plexiform, INL: inner nuclear layer, OPL: outer plexiform layer, ONL: outer nuclear layer, OLM: outer limiting membrane, RPE: retina pigment epithelium, ON: optic nerve.

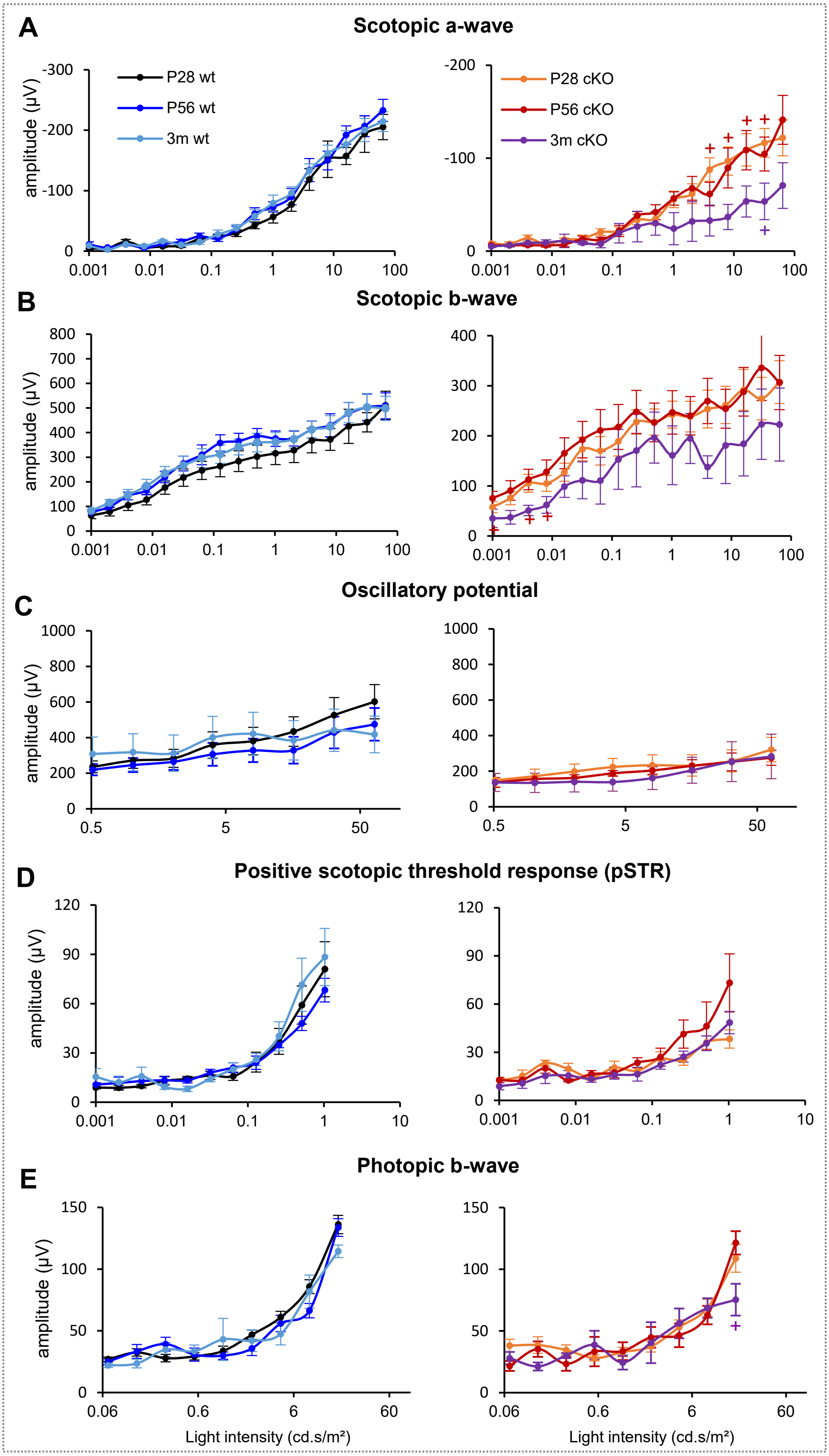


**Supplementary Figure 6: miRNA loss in late RPCs leads to impairments of rod function but only minimal alteration in cone function. A-B:** a-wave (A) and b-wave (B) of full-field scotopic electroretinogram recordings of wildtype (wt) and Ascl1-Cre:Dicer cKO$_{RPC}$ (cKO) mice at postnatal day P28, P56, and 3-months. **C:** Full-field scotopic electroretinogram recordings showing oscillatory potential amplitudes as selected wave forms and intensity dependent graphs for P28, P56, and 3-month old wildtypes (wt) and Dicer-cKO mice. **D:** Positive scotopic threshold response (pSTR) of wt and cKO at postnatal day P28, P56, and 3-months. **E:** Full-field photopic electroretinogram recordings showing b-wave amplitudes as selected wave forms and intensity-dependent graphs of wt and cKO at P28, P56, and 3-months. wt (n=10) and cKO mice (n=6), mean ± SEM. Mann-Whitney-U-test cKO vs cKO: +: $p \leq 0.05$.

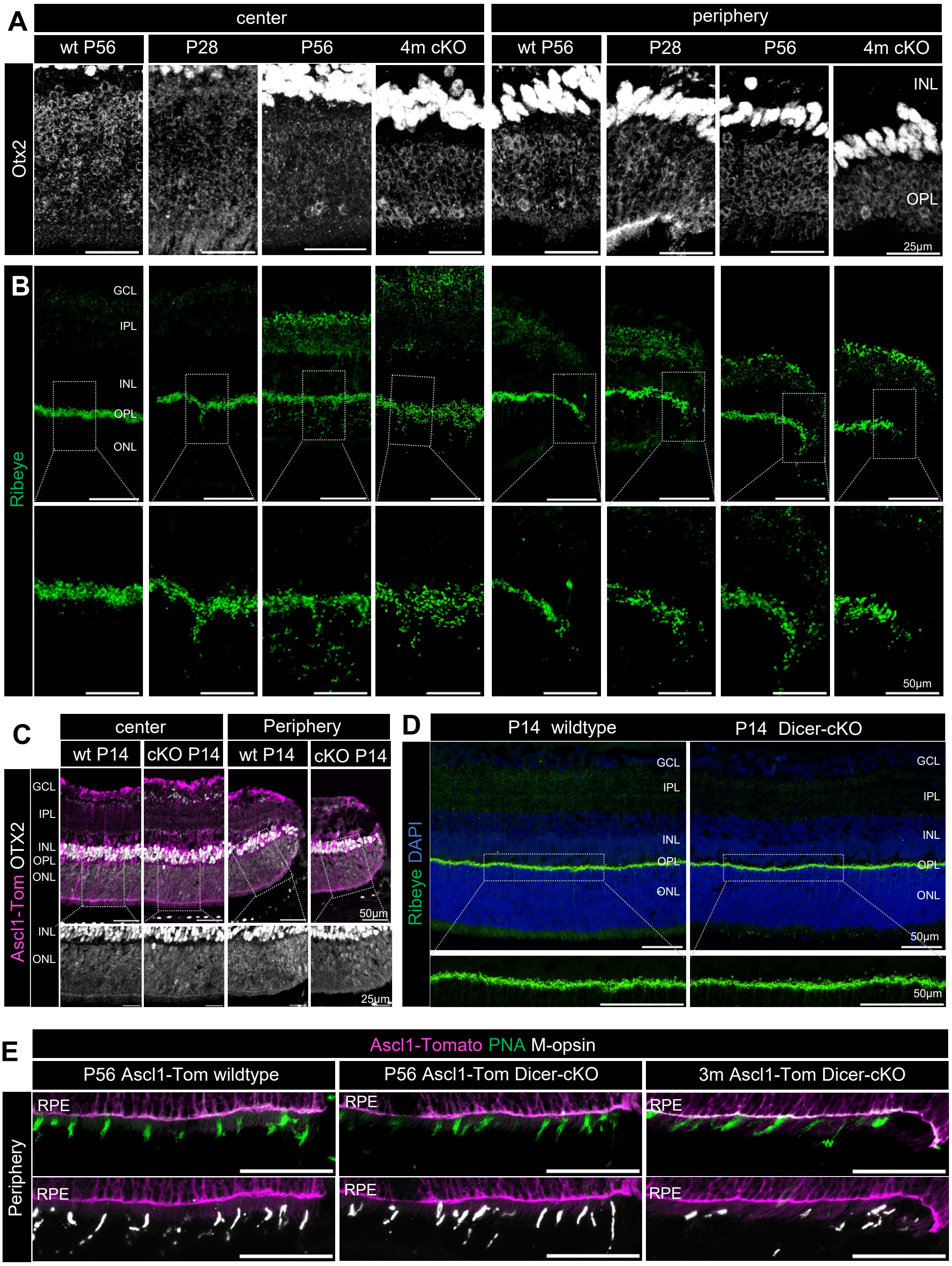


**Supplementary Figure 7: miRNA loss in late RPCs leads to reduced rod function and subsequent rod degeneration. A:** Immunofluorescent labeling using antibodies against Otx2 to label photoreceptor nuclei in the ONL in central or peripheral wildtype (wt) or Ascl1-Cre:Dicer cKO$_{RPC}$ (Dicer-cKO, cKO) retinas. **B:** Ribeye labeling to visualize ribbon synapse connections in central or peripheral retinas of P28 and P56 wildtype or Dicer-cKO retinas. **C:** Immunofluorescent labeling with antibodies against Otx2 in central or peripheral areas of P14 wildtype or Dicer-cKO retinas. **D:** Ribeye labeling in central or peripheral areas, as well as DAPI nuclear staining of P14 wildtype or Dicer-cKO retinas. **E:** Antibodies staining against peanut agglutinin (PNA) or to label entire cones or M-opsin to label L/M cones of peripheral P56 wildtype or Dicer–cKO retinas. Scale bars: 50 µm or 25 µm. GCL: ganglion cell layer, IPL: inner plexiform layer, INL: inner nuclear layer, OPL: outer plexiform layer, ONL: outer nuclear layer, RPE: retinal pigment epithelium.

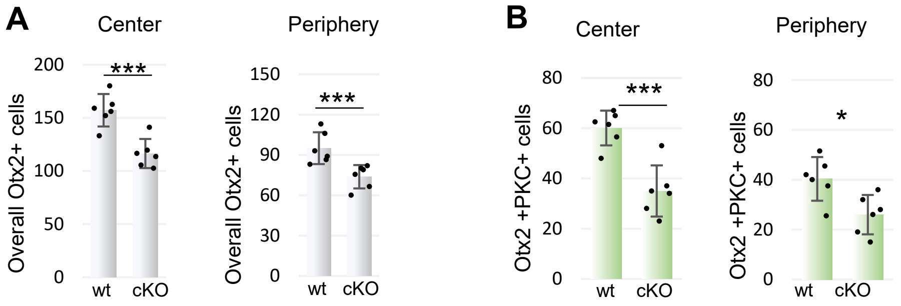


**Supplementary Figure 8. Bipolar cell numbers in P56 et and cKO retinas: A-B:** Absolute overall number of Otx2+ bipolar cells (BCs, A) and Otx2+ PKC+ rod BCs (B) in the central and peripheral retina of P56 wildtypes or Dicer-cKO mice; wt: n=6, cKO: n=6, mean ± S.D., Mann-Whitney-U-test: *: $p \leq 0.05$; **: $p \leq 0.01$; ***: $p \leq 0.005$. Cells per field.

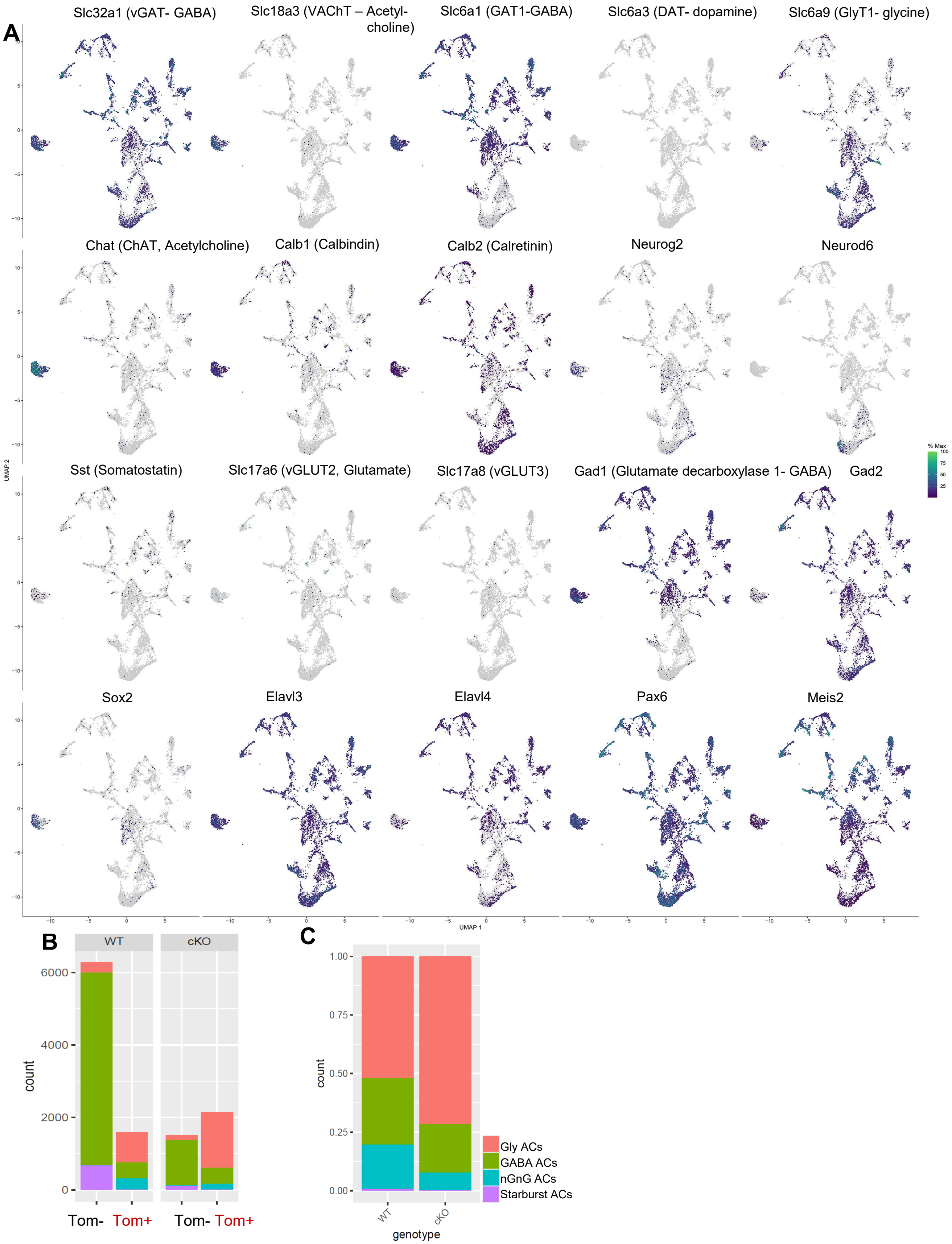


**Supplement Figure 9. scRNA-Seq of P7 progenies shows altered amacrine cell populations. A:** UMAP-dimension reduction of scRNA-seq FACS-purified integrated P7 Tomato+ wildtype (WT) and Dicer-cKO (cKO) amacrine cell (AC) progenies colored by annotated marker gene expression. **B:** AC type counts in scRNA-seq P7 wt and cKO AC Tomato+ progenies and Tomato- non-progenies, colored by annotated cell type as determined by marker gene expression. **C:** AC type proportions in scRNA-seq P7 wt and cKO AC progenies colored by annotated cell type as determined by marker gene expression.

### 3’UTR-seedless region of Elavl3

**Output for STarMir** — Prediction model training data (species): HITS-CLIP (mouse) — Target: Elavl3; miRNA: microRNA

Interactive site viewer for miRNA: all and target: Elavl3

3'UTR-seed | 3'UTR-seedless | CDS-seed | CDS-seedless | 5'UTR-seed | 5'UTR-seedless

| miRNA | Site_Position | Hybrid Conformation | LogitProb† | Score | 3'_bp* | $\Delta G_{hybrid}$ | $\Delta G_{nucl}$* | $\Delta G_{total}$* | Site_Access* | Upstream_Access (15nt)* | Dwstream_Access (15nt)* | Upstream_AU (30nt)* | Site_Location* |
|---|---|---|---|---|---|---|---|---|---|---|---|---|---|
| mmu-miR-25-3p | 2025-2043 | View | 0.810 | 0.536 | 0 | -21.300 | 0 | -3.450 | 0.529 | 0.983 | 0.415 | 0.800 | 0.142 |
| mmu-miR-25-3p | 2025-2039 | View | 0.800 | 0.536 | 0 | -15.800 | 0 | -3.957 | 0.655 | 0.983 | 0.376 | 0.800 | 0.142 |
| mmu-miR-25-3p | 2289-2299 | View | 0.781 | 0.536 | 0 | -20.200 | -1.013 | -7.103 | 0.400 | 0.788 | 0.240 | 0.700 | 0.217 |
| mmu-miR-25-3p | 2176-2188 | View | 0.770 | 0.536 | 0 | -19.200 | -2.154 | -5.358 | 0.383 | 0.639 | 0.492 | 0.767 | 0.185 |
| mmu-miR-25-3p | 2376-2395 | View | 0.768 | 0.536 | 0 | -17.600 | 0 | -5.450 | 0.511 | 0.963 | 0.853 | 0.500 | 0.242 |
| mmu-miR-25-3p | 2289-2304 | View | 0.768 | 0.536 | 0 | -22 | -1.287 | -6.087 | 0.367 | 0.788 | 0.200 | 0.700 | 0.217 |
| mmu-miR-25-3p | 2289-2306 | View | 0.689 | 0.536 | 0 | -24.500 | -1.287 | -0.904 | 0.381 | 0.788 | 0.156 | 0.700 | 0.217 |
| mmu-miR-25-3p | 2533-2550 | View | 0.673 | 0.536 | 0 | -22.700 | -0.020 | -9.655 | 0.330 | 0.507 | 0.646 | 0.533 | 0.287 |

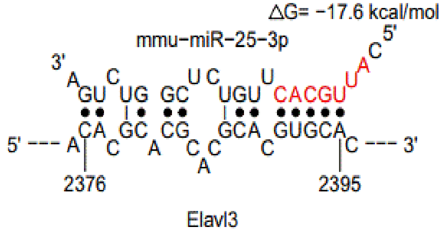


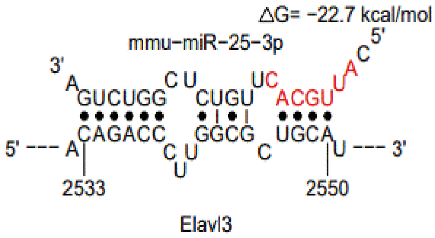


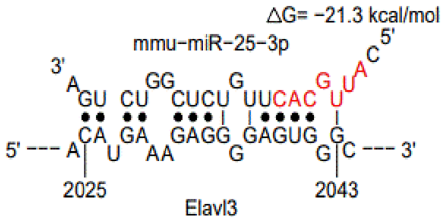


### CDS-seed region of Elavl3

**Output for STarMir** — Prediction model training data (species): HITS-CLIP (mouse) — Target: Elavl3; miRNA: microRNA

Interactive site viewer for miRNA: all and target: Elavl3

3'UTR-seed | 3'UTR-seedless | CDS-seed | CDS-seedless | 5'UTR-seed | 5'UTR-seedless

| miRNA | Site_Position | Hybrid Conformation | LogitProb† | Score | Seed_Position | Seed_Type* | 3'_bp* | $\Delta G_{hybrid}$ | $\Delta G_{nucl}$* | $\Delta G_{total}$* | Site_Access* | Seed_Access* | Upstream_Access (15nt)* | Dwstream_Access (15nt)* | Upstream_AU (30nt)* | Dwstream_AU (30nt)* | Site_ |
|---|---|---|---|---|---|---|---|---|---|---|---|---|---|---|---|---|---|
| mmu-miR-25-3p | 1369-1399 | View | 0.551 | 0.536 | 1394-1399 | offset-6mer | 1 | -27 | -0.294 | -12.139 | 0.441 | 0.507 | 0.484 | 0.365 | 0.400 | 0.467 | 0.855 |

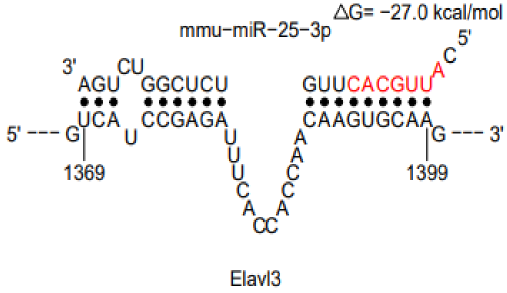


**Supplement Figure 10. Additional predicted binding sites for miR-25.** Screenshot of StarMiR results (tables) and interaction schematics showing additional miR-25-3p binding sites in the 3’UTR and CDS in the *Elavl3* mRNA. miRNA seed sequences are indicated in red. Guide values for ΔG (hybrid stability): < -15 kcal/mol, probability: ≥ 0.5 indicate high hybrid stability/probability. Selected highlighted examples are shown as interaction schematics.

| gene | P7 w1 | P7 wt2 | P7 wt3 | P7 wt4 | P7 cKO1 | P7 cKO2 | P7 cKO3 | P7 cKO4 |
|---|---|---|---|---|---|---|---|---|
| Aqp4 | 463 | 1123 | 821 | 1022 | 21 | 11 | 279 | 17 |
| Ascl1 | 905 | 1243 | 1119 | 1134 | 1850 | 1371 | 1480 | 2161 |
| Bak1 | 33 | 27 | 25 | 31 | 18 | 26 | 30 | 22 |
| Bax | 3322 | 1362 | 1025 | 1471 | 1459 | 1090 | 1217 | 1297 |
| Cabp5 | 7363 | 8027 | 6363 | 5923 | 1003 | 2247 | 2549 | 1617 |
| Calb1 | 260 | 707 | 532 | 570 | 50 | 598 | 615 | 552 |
| Calb2 | 747 | 1830 | 2044 | 962 | 995 | 1820 | 2124 | 1204 |
| Car2 | 65035 | 34155 | 27823 | 60612 | 29335 | 60450 | 53574 | 67692 |
| Casp3 | 4455 | 9286 | 6090 | 6035 | 5604 | 7923 | 6312 | 6328 |
| Casp6 | 1770 | 957 | 857 | 1307 | 1451 | 1789 | 1441 | 1696 |
| Casp7 | 960 | 657 | 524 | 528 | 447 | 691 | 657 | 596 |
| Casp8 | 124 | 156 | 213 | 154 | 111 | 355 | 226 | 335 |
| Casp9 | 1935 | 1425 | 1605 | 1784 | 2423 | 1558 | 1527 | 1548 |
| Casp9 | 1935 | 1425 | 1605 | 1784 | 2423 | 1558 | 1527 | 1548 |
| Casz1 | 3608 | 1808 | 2007 | 2702 | 5183 | 2478 | 2415 | 2487 |
| Ccnd1 | 1901 | 831 | 746 | 1577 | 4608 | 1705 | 1377 | 1772 |
| Dkk3 | 62122 | 66867 | 52061 | 66636 | 72876 | 80585 | 69856 | 86045 |
| Elavl3 | 568 | 539 | 416 | 524 | 940 | 795 | 1053 | 628 |
| Elavl4 | 474 | 1691 | 1883 | 1325 | 645 | 1770 | 2136 | 1883 |
| Glul | 4866 | 3972 | 3273 | 4559 | 3974 | 4607 | 4153 | 4391 |
| Gnat1 | 7988 | 982 | 2591 | 2727 | 1431 | 1370 | 1964 | 782 |
| Gsg1 | 3536 | 1901 | 1337 | 1426 | 1296 | 779 | 854 | 944 |
| Hes1 | 6897 | 3766 | 3186 | 6446 | 3775 | 6674 | 5497 | 12750 |
| Hes5 | 4097 | 3078 | 2436 | 3571 | 2097 | 4577 | 3339 | 6878 |
| Insm1 | 4341 | 3776 | 5683 | 5630 | 2149 | 5723 | 4603 | 3895 |
| Meis2 | 4527 | 5633 | 6541 | 5873 | 6782 | 6486 | 5960 | 5822 |
| Mki67 | 365 | 1286 | 2185 | 607 | 2918 | 4375 | 2138 | 2942 |
| Neurod1 | 17289 | 33342 | 47585 | 60398 | 31479 | 57001 | 51259 | 64913 |
| Neurog2 | 1447 | 2230 | 1437 | 2005 | 2131 | 1807 | 2285 | 4585 |
| Nfia | 2582 | 10110 | 7619 | 8745 | 3417 | 9313 | 9005 | 11303 |
| Nfib | 8202 | 24350 | 18898 | 17728 | 12847 | 21371 | 18302 | 20667 |
| Nfic | 3121 | 846 | 832 | 1580 | 3395 | 1223 | 1487 | 1491 |
| Nfix | 2178 | 1407 | 842 | 2142 | 2056 | 1815 | 2430 | 1882 |
| Notch1 | 4613 | 2308 | 1477 | 3540 | 12170 | 3299 | 3726 | 4973 |
| Notch3 | 226 | 179 | 127 | 254 | 369 | 211 | 206 | 225 |
| Nr2e3 | 122293 | 32227 | 49395 | 53307 | 67437 | 50596 | 48604 | 40599 |
| Nrl | 41994 | 13677 | 28423 | 23027 | 20758 | 23018 | 22588 | 16834 |
| Olig2 | 265 | 255 | 288 | 281 | 702 | 370 | 326 | 496 |
| Otx2 | 49237 | 70426 | 63653 | 49407 | 64394 | 59837 | 47575 | 58020 |
| Pax6 | 6914 | 12443 | 9566 | 12470 | 9859 | 16936 | 16905 | 19142 |
| Pcna | 4225 | 4488 | 5994 | 3858 | 3990 | 6432 | 5823 | 5565 |
| Pcp2 | 1785 | 574 | 207 | 1177 | 109 | 93 | 186 | 118 |
| Pde6b | 79832 | 27069 | 50624 | 42745 | 43809 | 40666 | 43775 | 25862 |
| Prkca | 6784 | 11944 | 8331 | 10395 | 2627 | 4534 | 5128 | 5316 |
| Rax | 9037 | 1959 | 1509 | 4305 | 8091 | 2714 | 3536 | 4361 |
| Rcvrn | 13063 | 1993 | 3483 | 3868 | 1914 | 2473 | 3773 | 1939 |
| Rho | 210769 | 37802 | 85932 | 91998 | 63431 | 68416 | 90446 | 49855 |
| Rlbp1 | 60708 | 30255 | 24088 | 39149 | 37776 | 42692 | 37809 | 44712 |
| Slc1a3 | 44546 | 80362 | 63417 | 75463 | 34539 | 96305 | 79555 | 100466 |
| Snap25 | 62349 | 93462 | 117633 | 95424 | 35215 | 93656 | 84557 | 67788 |
| Sox2 | 1264 | 1812 | 1313 | 2734 | 1260 | 2957 | 2123 | 4242 |
| Sox9 | 6186 | 5875 | 4630 | 7677 | 9887 | 7070 | 8033 | 12869 |
| Sun1 | 7488 | 3580 | 4068 | 5399 | 10112 | 4527 | 5474 | 5085 |
| Syne1 | 130 | 241 | 137 | 238 | 121 | 155 | 149 | 145 |
| Tgs1 | 2217 | 7109 | 7328 | 4371 | 2806 | 5683 | 5088 | 5653 |
| Tmem215 | 13993 | 25853 | 17930 | 18902 | 5208 | 9888 | 10337 | 8836 |
| Trpm1 | 6924 | 4607 | 2201 | 3453 | 618 | 932 | 1554 | 1080 |
| Vsx1 | 4454 | 10353 | 7915 | 6397 | 1616 | 3777 | 3754 | 3025 |
| Vsx2 | 102155 | 75834 | 59592 | 72119 | 54051 | 42040 | 43055 | 44939 |

**Supplementary Table 1:** Expression (normalized counts) of 60 selected genes (alphabetical order) in P7 wildtype (wt) and Dicer-cKO from bulk RNA-seq.

| gene/miR | Accession number | wt | wt2 | wt3 | cko1 | cKO2 | cKO3 | % reduction |
|---|---|---|---|---|---|---|---|---|
| Mirlet7b | MIMAT0000522 | 140279 | 115562 | 117639 | 54744 | 89637 | 67492 | -43 |
| Mir183 | MIMAT0000212 | 58101 | 80167 | 61463 | 10395 | 30107 | 16059 | -72 |
| Mirlet7g | MIMAT0000121 | 34245 | 28671 | 25254 | 12371 | 31046 | 14956 | -34 |
| Mirlet7i | MIMAT0000122 | 18260 | 15204 | 14813 | 6818 | 12077 | 7622 | -45 |
| Mir342 | MIMAT0000590 | 15699 | 14074 | 10122 | 6207 | 13386 | 9176 | -28 |
| Mir204 | MIMAT0000237 | 15898 | 11639 | 11801 | 3210 | 13630 | 10734 | -30 |
| Mir182 | MIMAT0000211 | 7100 | 9398 | 6927 | 2041 | 9029 | 3638 | -37 |
| Mir423 | MIMAT0003454 | 5293 | 5817 | 5007 | 3412 | 5077 | 4633 | -19 |
| Mir130b | MIMAT0000387 | 5453 | 5102 | 4643 | 3795 | 4856 | 3300 | -21 |
| Mir328 | MIMAT0000565 | 5206 | 5173 | 4809 | 4987 | 6823 | 5365 | 13 |
| Mir23b | MIMAT0000125 | 4779 | 4611 | 3805 | 1840 | 4092 | 3094 | -32 |
| Mir99b | MIMAT0000132 | 3913 | 4101 | 3233 | 1294 | 3180 | 2218 | -40 |
| Mir98 | MIMAT0000545 | 4273 | 3363 | 2821 | 704 | 2289 | 956 | -62 |
| Mir674 | MIMAT0003740 | 3610 | 3202 | 2990 | 3405 | 3658 | 3024 | 3 |
| Mir130a | MIMAT0000141 | 3520 | 3241 | 2954 | 5124 | 4031 | 2977 | 25 |
| Mirlet7f | MIMAT0000525 | 4148 | 3037 | 2489 | 1046 | 3119 | 1424 | -42 |
| Mir210 | MIMAT0000658 | 3560 | 2613 | 3055 | 2652 | 2989 | 1932 | -18 |
| Mirlet7c | MIMAT0000523 | 3289 | 2808 | 2355 | 1006 | 2062 | 1456 | -46 |
| Mir151 | MIMAT0004536 | 2558 | 2624 | 1968 | 1504 | 2021 | 1760 | -26 |
| Mir30a | MIMAT0000128 | 2549 | 2138 | 1843 | 1540 | 2605 | 1365 | -16 |
| Mir744 | MIMAT0004187 | 2172 | 2158 | 1931 | 962 | 1487 | 1488 | -37 |
| Mir92b | MIMAT0004899 | 1923 | 1827 | 1801 | 469 | 1430 | 1958 | -31 |
| Mirlet7a | MIMAT0000521 | 1998 | 1965 | 1502 | 601 | 1785 | 1010 | -38 |
| Mir15b | MIMAT0000124 | 1887 | 1853 | 1366 | 635 | 1794 | 1172 | -29 |
| Mir149 | MIMAT0000159 | 1830 | 1531 | 1540 | 876 | 2098 | 1418 | -10 |
| Mir296 | MIMAT0000374 | 1525 | 1449 | 1256 | 1312 | 1549 | 1239 | -3 |
| Mir484 | MIMAT0003127 | 1415 | 1474 | 1106 | 351 | 1330 | 911 | -35 |
| Mir298 | MIMAT0000376 | 1316 | 1373 | 1104 | 1354 | 983 | 686 | -20 |
| Mir181c | MIMAT0000674 | 1016 | 1484 | 1116 | 643 | 909 | 549 | -42 |
| Mir7a | MIMAT0000677 | 1267 | 1147 | 1119 | 720 | 1290 | 849 | -19 |
| Mir532 | MIMAT0004781 | 1300 | 1202 | 1016 | 969 | 1314 | 1052 | -5 |
| Mir23a | MIMAT0000532 | 1231 | 1022 | 820 | 460 | 942 | 737 | -30 |
| Mir26b | MIMAT0000534 | 1171 | 1051 | 773 | 176 | 1064 | 512 | -42 |
| Mir382 | MIMAT0000747 | 1052 | 1080 | 701 | 564 | 843 | 486 | -33 |
| Mir672 | MIMAT0003735 | 1012 | 845 | 786 | 247 | 766 | 447 | -45 |
| Mir106b | MIMAT0000386 | 888 | 883 | 727 | 584 | 716 | 485 | -29 |
| Mir184 | MIMAT0000213 | 800 | 870 | 708 | 247 | 861 | 906 | -15 |
| Mir96 | MIMAT0000541 | 687 | 891 | 741 | 347 | 823 | 352 | -34 |
| Mir140 | MIMAT0000151 | 658 | 665 | 612 | 638 | 855 | 589 | 8 |
| Mir211 | MIMAT0000668 | 994 | 322 | 504 | 573 | 558 | 223 | -26 |
| Mir335 | MIMAT0000766 | 732 | 480 | 599 | 125 | 1001 | 482 | -11 |
| Mir384 | MIMAT0004745 | 759 | 483 | 521 | 133 | 651 | 269 | -40 |
| Mir361 | MIMAT0000704 | 524 | 648 | 437 | 469 | 488 | 382 | -17 |
| Mir34a | MIMAT0000542 | 474 | 623 | 506 | 174 | 306 | 271 | -53 |
| Mir340 | MIMAT0004651 | 602 | 452 | 477 | 123 | 464 | 209 | -48 |
| Mir326 | MIMAT0000559 | 502 | 447 | 422 | 499 | 566 | 276 | -2 |
| Mir30b | MIMAT0000130 | 447 | 442 | 405 | 149 | 598 | 299 | -19 |
| Mir323 | MIMAT0000551 | 466 | 472 | 321 | 136 | 448 | 181 | -39 |
| Mir351 | MIMAT0000609 | 431 | 396 | 381 | 147 | 441 | 387 | -19 |
| Mir676 | MIMAT0003782 | 404 | 435 | 354 | 153 | 326 | 281 | -36 |

**Supplementary Table 2:** Top 50 highly expressed P7 wildtype (wt) miRNAs and their expression levels in the P7 Dicer-cKO retina (highest to lowest wt expression, normalized counts, bulk RNA-seq).

| | GO biological process complete | Mus musculus - REFLIST (21836) | 171 | expected | over/ under | fold Enrichment | raw P-value | FDR |
|---|---|---|---|---|---|---|---|---|
| 1 | cell-cell adhesion involved in synapse maturation (GO:0090125) | 2 | 2 | 0.02 | + | > 100 | 6.10E-05 | 3.99E-03 |
| 2 | positive regulation of neuromuscular synaptic transmission (GO:1900075) | 3 | 2 | 0.02 | + | 85.13 | 1.82E-04 | 9.28E-03 |
| 3 | regulation of neuromuscular synaptic transmission (GO:1900073) | 3 | 2 | 0.02 | + | 85.13 | 1.82E-04 | 9.25E-03 |
| 4 | nucleokinesis involved in cell motility in cerebral cortex radial glia guided migration (GO:0021817) | 3 | 2 | 0.02 | + | 85.13 | 1.82E-04 | 9.22E-03 |
| 5 | ascending aorta morphogenesis (GO:0035910) | 4 | 2 | 0.03 | + | 63.85 | 3.62E-04 | 1.58E-02 |
| 6 | nuclear migration along microtubule (GO:0030473) | 4 | 2 | 0.03 | + | 63.85 | 3.62E-04 | 1.58E-02 |
| 7 | modulation of microtubule cytoskeleton involved in cerebral cortex radial glia guided migration (GO:0021815) | 4 | 2 | 0.03 | + | 63.85 | 3.62E-04 | 1.57E-02 |
| 8 | negative regulation of amacrine cell differentiation (GO:1902870) | 4 | 2 | 0.03 | + | 63.85 | 3.62E-04 | 1.57E-02 |
| 9 | retrograde trans-synaptic signaling by trans-synaptic protein complex (GO:0098942) | 5 | 2 | 0.04 | + | 51.08 | 6.00E-04 | 2.27E-02 |
| 10 | ascending aorta development (GO:0035905) | 5 | 2 | 0.04 | + | 51.08 | 6.00E-04 | 2.26E-02 |
| 11 | positive regulation of basement membrane assembly involved in embryonic body morphogenesis (GO:1904261) | 5 | 2 | 0.04 | + | 51.08 | 6.00E-04 | 2.25E-02 |
| 12 | regulation of basement membrane assembly involved in embryonic body morphogenesis (GO:1904259) | 5 | 2 | 0.04 | + | 51.08 | 6.00E-04 | 2.25E-02 |
| 13 | metanephric nephron tubule morphogenesis (GO:0072282) | 8 | 3 | 0.06 | + | 47.89 | 2.57E-05 | 1.92E-03 |
| 14 | primitive erythrocyte differentiation (GO:0060319) | 6 | 2 | 0.05 | + | 42.57 | 8.96E-04 | 3.12E-02 |
| 15 | vascular endothelial growth factor receptor-2 signaling pathway (GO:0036324) | 6 | 2 | 0.05 | + | 42.57 | 8.96E-04 | 3.11E-02 |
| 16 | NMDA glutamate receptor clustering (GO:0097114) | 6 | 2 | 0.05 | + | 42.57 | 8.96E-04 | 3.10E-02 |
| 17 | cell motility involved in cerebral cortex radial glia guided migration (GO:0021814) | 6 | 2 | 0.05 | + | 42.57 | 8.96E-04 | 3.10E-02 |
| 18 | positive regulation of peptidyl-lysine acetylation (GO:2000758) | 7 | 2 | 0.05 | + | 36.48 | 1.25E-03 | 4.00E-02 |
| 19 | contact inhibition (GO:0060242) | 7 | 2 | 0.05 | + | 36.48 | 1.25E-03 | 4.00E-02 |
| 20 | negative regulation of hepatocyte proliferation (GO:2000346) | 7 | 2 | 0.05 | + | 36.48 | 1.25E-03 | 3.99E-02 |
| 21 | pointed-end actin filament capping (GO:0051694) | 7 | 2 | 0.05 | + | 36.48 | 1.25E-03 | 3.98E-02 |
| 22 | positive regulation of extracellular matrix assembly (GO:1901203) | 14 | 4 | 0.11 | + | 36.48 | 3.42E-06 | 3.41E-04 |
| 23 | angiogenesis involved in coronary vascular morphogenesis (GO:0060978) | 7 | 2 | 0.05 | + | 36.48 | 1.25E-03 | 3.97E-02 |
| 24 | negative regulation of photoreceptor cell differentiation (GO:0046533) | 7 | 2 | 0.05 | + | 36.48 | 1.25E-03 | 3.96E-02 |
| 25 | postsynaptic density protein 95 clustering (GO:0097119) | 7 | 2 | 0.05 | + | 36.48 | 1.25E-03 | 3.95E-02 |
| 26 | common bile duct development (GO:0061009) | 7 | 2 | 0.05 | + | 36.48 | 1.25E-03 | 3.95E-02 |
| 27 | metanephric tubule morphogenesis (GO:0072173) | 11 | 3 | 0.09 | + | 34.83 | 7.43E-05 | 4.74E-03 |
| 28 | commissural neuron axon guidance (GO:0071679) | 15 | 4 | 0.12 | + | 34.05 | 4.63E-06 | 4.42E-04 |
| 29 | detection of cell density (GO:0060245) | 8 | 2 | 0.06 | + | 31.92 | 1.66E-03 | 4.91E-02 |
| 30 | negative regulation of hepatocyte apoptotic process (GO:1903944) | 8 | 2 | 0.06 | + | 31.92 | 1.66E-03 | 4.90E-02 |
| 31 | male sex determination (GO:0030238) | 16 | 4 | 0.13 | + | 31.92 | 6.14E-06 | 5.67E-04 |
| 32 | regulation of photoreceptor cell differentiation (GO:0046532) | 8 | 2 | 0.06 | + | 31.92 | 1.66E-03 | 4.89E-02 |
| 33 | regulation of amacrine cell differentiation (GO:1902869) | 8 | 2 | 0.06 | + | 31.92 | 1.66E-03 | 4.88E-02 |
| 34 | positive regulation of vascular endothelial growth factor receptor signaling pathway (GO:0030949) | 14 | 3 | 0.11 | + | 27.36 | 1.61E-04 | 8.59E-03 |
| 35 | metanephric nephron morphogenesis (GO:0072273) | 19 | 4 | 0.15 | + | 26.88 | 1.28E-05 | 1.08E-03 |
| 36 | regulation of extracellular matrix assembly (GO:1901201) | 20 | 4 | 0.16 | + | 25.54 | 1.60E-05 | 1.29E-03 |
| 37 | angiogenesis involved in wound healing (GO:0060055) | 16 | 3 | 0.13 | + | 23.94 | 2.45E-04 | 1.18E-02 |
| 38 | endothelial cell morphogenesis (GO:0001886) | 17 | 3 | 0.13 | + | 22.53 | 2.96E-04 | 1.37E-02 |
| 39 | peptidyl-tyrosine autophosphorylation (GO:0038083) | 17 | 3 | 0.13 | + | 22.53 | 2.96E-04 | 1.36E-02 |
| 40 | ventricular trabecula myocardium morphogenesis (GO:0003222) | 17 | 3 | 0.13 | + | 22.53 | 2.96E-04 | 1.36E-02 |
| 41 | positive regulation of branching involved in ureteric bud morphogenesis (GO:0090190) | 23 | 4 | 0.18 | + | 22.21 | 2.86E-05 | 2.06E-03 |
| 42 | metanephric nephron tubule development (GO:0072234) | 18 | 3 | 0.14 | + | 21.28 | 3.53E-04 | 1.56E-02 |
| 43 | microtubule anchoring (GO:0034453) | 24 | 4 | 0.19 | + | 21.28 | 3.41E-05 | 2.38E-03 |
| 44 | regulation of branching involved in ureteric bud morphogenesis (GO:0090189) | 25 | 4 | 0.2 | + | 20.43 | 4.04E-05 | 2.76E-03 |
| 45 | metanephros morphogenesis (GO:0003338) | 26 | 4 | 0.2 | + | 19.65 | 4.74E-05 | 3.19E-03 |
| 46 | nuclear migration (GO:0007097) | 33 | 5 | 0.26 | + | 19.35 | 5.52E-06 | 5.19E-04 |
| 47 | pharyngeal system development (GO:0060037) | 34 | 5 | 0.27 | + | 18.78 | 6.43E-06 | 5.87E-04 |
| 48 | metanephric nephron epithelium development (GO:0072243) | 21 | 3 | 0.16 | + | 18.24 | 5.66E-04 | 2.16E-02 |
| 49 | positive regulation of mesenchymal cell proliferation (GO:0002053) | 35 | 5 | 0.27 | + | 18.24 | 7.45E-06 | 6.73E-04 |
| 50 | sex determination (GO:0007530) | 28 | 4 | 0.22 | + | 18.24 | 6.42E-05 | 4.16E-03 |

**Supplementary Table 3:** Gene Ontology (GO) analysis showing the top 50 biological processes of genes identified as P7 miRNA targets and to be upregulated in the P7 cKO.

| | miR ID | Accession number | Nanostring P2 RPC counts | RNA-Seq bulk P7 RPC count |
|---|---|---|---|---|
| **1** | **mmu-miR-9** | **MIMAT0000142** | **63981** | **83** |
| **2** | **mmu-miR-16** | **MIMAT0000527** | **35845** | **40** |
| 3 | mmu-miR-204 | MIMAT0000237 | 25967 | 9191 |
| **4** | **mmu-miR-181a** | **MIMAT0000210** | **25080** | **100** |
| **5** | **mmu-miR-20a** | **MIMAT0000529** | **21395** | **0** |
| **6** | **mmu-miR-25** | **MIMAT0000652** | **20511** | **0** |
| 7 | mmu-miR-15b | MIMAT0000124 | 20032 | 1200 |
| 8 | mmu-let-7g | MIMAT0000121 | 15750 | 19458 |
| **9** | **mmu-let-7d** | **MIMAT0000383** | **15516** | **0** |
| 10 | mmu-miR-96 | MIMAT0000541 | 12811 | 507 |
| 11 | mmu-let-7i | MIMAT0000122 | 12334 | 8839 |
| **12** | **mmu-miR-125a-5p** | **MIMAT0000135** | **9841** | **0** |
| 13 | mmu-let-7a | MIMAT0000521 | 9477 | 1132 |
| **14** | **mmu-miR-15a** | **MIMAT0000526** | **8593** | **81** |
| **15** | **mmu-miR-1944** | **MIMAT0009409** | **8207** | **Not found** |
| 16 | mmu-miR-183 | MIMAT0000212 | 7352 | 18854 |
| 17 | mmu-let-7c | MIMAT0000523 | 7030 | 1508 |
| 18 | mmu-miR-124 | MIMAT0000134 | 7015 | 190 |
| 19 | mmu-let-7b | MIMAT0000522 | 6889 | 70624 |
| **20** | **mmu-miR-19a** | **MIMAT0000651** | **6501** | **0** |
| 21 | mmu-miR-125b-5p | MIMAT0000136 | 4547 | 80 |
| 22 | mmu-miR-301a | MIMAT0000379 | 4339 | Not found |
| 23 | mmu-miR-30c | MIMAT0000514 | 4087 | 40 |
| 24 | mmu-let-7e | MIMAT0000524 | 4076 | 0 |
| 25 | mmu-miR-135a | MIMAT0000147 | 3862 | 30 |
| 26 | mmu-miR-130a | MIMAT0000141 | 3847 | 4044 |
| 27 | mmu-let-7f | MIMAT0000525 | 3836 | 1863 |
| 28 | mmu-miR-19b | MIMAT0000513 | 3405 | 7 |
| 29 | mmu-miR-130b | MIMAT0000387 | 2976 | 3983 |
| 30 | mmu-miR-1937a | MIMAT0009401 | 2592 | Not found |
| 31 | mmu-miR-182 | MIMAT0000211 | 2267 | 4903 |
| 32 | mmu-miR-93 | MIMAT0000540 | 2106 | 0 |
| 33 | mmu-miR-342-3p | MIMAT0000590 | 1897 | 9589 |
| 34 | mmu-miR-106b | MIMAT0000386 | 1689 | 595 |
| 35 | mmu-miR-99b | MIMAT0000132 | 1669 | 2231 |
| 36 | mmu-miR-103 | MIMAT0000546 | 1479 | 20 |
| 37 | mmu-miR-340-5p | MIMAT0004651 | 1380 | 265 |
| 38 | mmu-miR-99a | MIMAT0000131 | 1375 | 185 |
| 39 | mmu-miR-335-5p | MIMAT0000766 | 1352 | 536 |
| 40 | mmu-miR-30d | MIMAT0000515 | 1330 | 0 |
| 41 | mmu-miR-1937c | MIMAT0009429 | 1299 | Not found |
| 42 | mmu-miR-350 | MIMAT0000605 | 1212 | 261 |
| 43 | mmu-miR-872 | MIMAT0004934 | 1043 | 192 |
| 44 | mmu-miR-210 | MIMAT0000658 | 964 | 2525 |
| 45 | mmu-miR-151-5p | MIMAT0004536 | 934 | 1762 |
| 46 | mmu-miR-191 | MIMAT0000221 | 917 | 0 |
| 47 | mmu-miR-148a | MIMAT0000516 | 822 | 32 |
| 48 | mmu-miR-30b | MIMAT0000130 | 770 | 349 |
| 49 | mmu-miR-29c | MIMAT0000536 | 760 | 0 |
| 50 | mmu-miR-30a | MIMAT0000128 | 691 | 1837 |

**Supplementary Table 4.** 50 miRNAs highly expressed in P2 RPCs (Nanostring, *Wohl et. al., 2019*) and P7 RPCs/PCs (bulk RNA-Seq, averaged normalized counts). Top 10 miRNAs found only at P2 are highlighted in dark blue. *Expression levels *per se* cannot be compared as data were obtained from different techniques.

| miRNA | Tools | Ensembl ID | Ensembl transcript | RefSeq ID | miRNA binding site in mRNA | binding region in mRNA | true target predicted probability | thermodynamic stability energy of duplex, < -15 | miRNA seed binding |
|---|---|---|---|---|---|---|---|---|---|
| mmu-miR-20a-5p | miRWalk | ENSMUST00000003501.9 | Canonical-current | NM_010487.2 | 660-676 | CDS | 1 | -20.6 | 0 |
| | STarMir | ENSMUST00000003501.9 | Canonical-current | NM_010487.2 | 2389-2436 | 3'UTR | 0.405 | -15.7 | 1 |
| | DT: TarBase | ENSMUST00000003501.9 | n/a | n/a | n/a | n/a | n/a | n/a | n/a |
| | TargetScan | ENSMUST00000003501.7 | Canonical - outdated | n/a | n/a | n/a | n/a | n/a | n/a |
| mmu-miR-15b-5p | miRWalk | ENSMUST00000003501.9 | Canonical-current | NM_010487.2 | 2585-2609 | 3'UTR | 0.846 | -18.6 | 1 |
| | STarMir | ENSMUST00000003501.9 | Canonical-current | NM_010487.2 | 2586-2601 | 3'UTR | 0.433 | -23.1 | 0 |
| | DT: TarBase | ENSMUST00000003501.9 | n/a | n/a | n/a | n/a | n/a | n/a | n/a |
| | TargetScan | ENSMUST00000003501.7 | Canonical - outdated | n/a | n/a | n/a | n/a | n/a | n/a |
| mmu-miR-25-3p | miRWalk | ENSMUST00000003501.9 | Canonical-current | NM_010487.2 | 1755-1806 | 3'UTR | 0.846 | -21.2 | 1 |
| | STarMir | ENSMUST00000003501.9 | Canonical-current | NM_010487.2 | 1756-1774 | 3'UTR | 0.411 | -25.1 | 0 |
| | STarMir | ENSMUST00000003501.9 | Canonical-current | NM_010487.2 | 1369-1399 | CDS | 0.551 | -27 | 0 |
| | STarMir | ENSMUST00000003501.9 | Canonical-current | NM_010487.2 | 2025-2043 | 3'UTR | 0.810 | -21.3 | 0 |
| | STarMir | ENSMUST00000003501.9 | Canonical-current | NM_010487.2 | 2376-2395 | 3'UTR | 0.768 | -17.6 | 0 |
| | STarMir | ENSMUST00000003501.9 | Canonical-current | NM_010487.2 | 2533-2550 | 3'UTR | 0.673 | -22.7 | 0 |
| | DT: TarBase | ENSMUST00000003501.9 | n/a | n/a | n/a | n/a | n/a | n/a | n/a |
| | TargetScan | ENSMUST00000003501.7 | Canonical - outdated | n/a | n/a | n/a | n/a | n/a | n/a |
| mmu-miR-124-3p | miRWalk | ENSMUST00000003501.9 | Canonical-current | NM_010487.2 | 3147-3164 | 3'UTR | 0.923 | -23.3 | 0 |
| | STarMir | ENSMUST00000003501.9 | Canonical-current | NM_010487.2 | 3148-3163 | 3'UTR | 0.667 | -28.5 | 0 |
| | DT: TarBase | ENSMUST00000003501.9 | n/a | n/a | n/a | n/a | n/a | n/a | n/a |
| | TargetScan | ENSMUST00000003501.7 | Canonical - outdated | n/a | n/a | n/a | n/a | n/a | n/a |

**Supplementary Table 5:** List of predicted P2 RPC miRNAs, prediction tools used, and outcomes for *Elavl3* mRNA interaction. Guide values for ΔG (hybrid stability): < -15 kcal/mol, probability: ≥ 0.5 indicate high hybrid stability/probability. CDS: coding sequence, outdated Ensembl IDs are shown in red, matching sequences across different tools are shown in green. DT: Diana Tools.
